# A gene regulatory network for specification and morphogenesis of a Mauthner Cell homolog in non-vertebrate chordates

**DOI:** 10.1101/2024.01.15.575616

**Authors:** Kwantae Kim, Katarzyna M. Piekarz, Alberto Stolfi

## Abstract

Transcriptional regulation of gene expression is an indispensable process in multicellular development, yet we still do not fully understand how the complex networks of transcription factors operating in neuronal precursors coordinately control the expression of effector genes that shape morphogenesis and terminal differentiation. Here we break down in greater detail a provisional regulatory circuit downstream of the transcription factor Pax3/7 operating in the descending decussating neurons (ddNs) of the tunicate *Ciona robusta.* The ddNs are a pair of hindbrain neurons proposed to be homologous to the Mauthner cells of anamniotes, and Pax3/7 is sufficient and necessary for their specification. We show that different transcription factors downstream of Pax3/7, namely Pou4, Lhx1/5, and Dmbx, regulate distinct “branches” of this ddN network that appear to be dedicated to different developmental tasks. Some of these network branches are shared with other neurons throughout the larva, reinforcing the idea that modularity is likely a key feature of such networks. We discuss these ideas and their evolutionary implications here, including the observation that homologs of all four transcription factors (Pax3/7, Lhx5, Pou4f3, and Dmbx1) are key for the specification of cranial neural crest in vertebrates.

## Introduction

During the development of multicellular organisms, specialized cell types and their immediate progenitors deploy the coordinated expression of genes encoding various proteins that shape their morphogenesis or confer their unique functional properties (Brunet and King, 2017). Key features of this “terminal selection” include modularity of gene regulatory networks (GRNs) and *cis*-regulatory elements (Hobert, 2008; Hobert and Kratsios, 2019), and the combinatorial activity of transcription factors (Zeitlinger, 2020). While modularity allows for a variety of mechanisms to control similar gene expression in distinct cell types or subtypes (Hatleberg and Hinman, 2021), combinatorial activity ensures a greater variety of unique gene expression profiles, often through synergistic effects (Farley et al., 2015; Farley et al., 2016). It is thought that both modularity and combinatorial features (“mix and match”) drive the diversification and specialization of cell types in multicellular development and evolution (Arendt et al., 2016).

Solitary tunicates in the genus *Ciona* have emerged as model organisms for the study of transcriptional regulation in animal development (Di Gregorio and Levine, 2002). As the sister group to vertebrates within phylum Chordata (Delsuc et al., 2006; Putnam et al., 2008), tunicates share many evolutionary innovations with their close vertebrate relatives, including novel genes, cell types, and developmental processes (Fodor et al., 2021). One potential homologous neuronal subtype shared by tunicates and vertebrates is the Mauthner cell, or M-cell. The M-cells of anamniotes such as fish and amphibians are giant reticulospinal neurons arising from the hindbrain and control fast escape responses to auditory stimuli (Hale et al., 2016). The descending decussating neuron (ddN) in the motor ganglion (MG) of the tunicate larva has been proposed as an M-cell homolog based on their developmental origins in the hindbrain, unusual descending, decussating (i.e. contralateral) axon projection, and connectivity within the neuronal network for motor control in swimming (**Figure 1A**)(Ryan et al., 2017). Although their roles in larval swimming or escape responses have yet to be experimentally determined, the ddNs are conserved in distantly related tunicate species, hinting at a deeply conserved function (Lowe and Stolfi, 2018). In contrast, the developmental cell lineage and gene expression profile of the ddNs in *Ciona robusta* are known in great detail, while relatively little is known about the regulation of M-cell development in vertebrates. Therefore, comparative studies on tunicate ddNs might provide some leads on understanding the development and evolution of the Mauthner cell.

**Figure 1.**
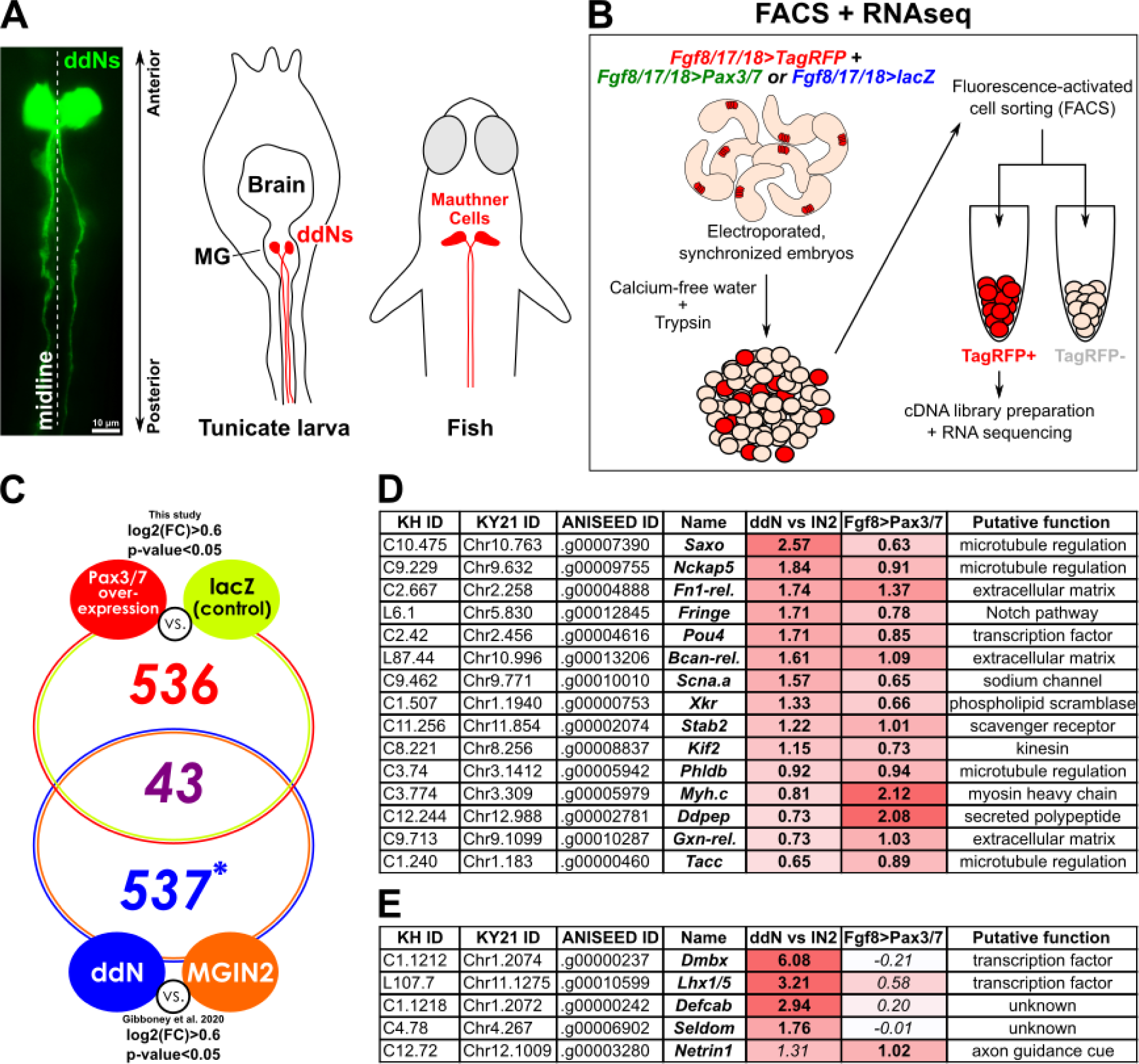
Bulk RNA sequencing of FACS-isolated cells to identify ddN-specific transcripts upregulated by Pax3/7 overexpression. A) The descending decussating neurons (ddNs) of a Ciona larva at 19.5 hours post-fertilization (hpf at 20°C, ∼ Hotta stage 29) labeled by expression of elecroporated *Dmbx/Defcab>Unc-76::YFP* reporter plasmid. Scale bar = 10 μm. Right: comparative diagrams of ddNs and their putative vertebrate homologs, the Mauthner cells (species depicted as an example is *Danio rerio*). B) Schematic diagram of FACS-RNAseq workflow (see text for details). C) Venn diagram of transcripts in common between significantly enriched (Log2 fold-change > 0.6 and p-value < 0.05) transcripts upregulated by Pax3/7 overexpression from the current study, and in the ddN relative to MGIN2 (MG interneuron 2) from Gibboney et al. 2020. Asterisk indicates number of microarray probesets, not transcript models. D) Selected examples of transcripts from the overlap shown in (C). Log2(FC) values color-coded based on rank. E) Selected examples of transcripts not included in the overlap shown in (C) but notable due to previously having been confirmed as enriched in the ddNs by *in situ* hybridization or reporter gene expression. Italicized numbers indicate p > 0.05.

Previous work on the ddNs revealed that the transcription factor Pax3/7 is necessary and sufficient for the specification of ddN fate and its unique morphology. Tissue-specific CRISPR/Cas9 mutagenesis of *Pax3/7* abolished ddN fate (Kim et al., 2022), while overexpression of Pax3/7 throughout the MG resulted in ectopic ddN-like neurons that also projected axons across the midline (Stolfi et al., 2011). Furthermore, microarray-based transcriptome of isolated ddNs revealed several novel candidate effectors of ddN development and function (Gibboney et al., 2020). However, it remains unknown how Pax3/7 and downstream regulators shape the transcriptional profile of the ddNs. Here we use a combination of CRISPR/Cas9 mutagenesis, ectopic overexpression, and *cis-*regulatory reporter analyses to study in greater detail the GRN operating downstream of Pax3/7 in the ddNs of *Ciona.* We find that different branches of this transcriptional network regulate distinct effector genes. Some branches are shared with distinct neuron types in other parts of the *Ciona* larval nervous system, highlighting the modular and combinatorial nature of such networks. We discuss the functional and evolutionary implications of these findings in the context of chordate neural development.

## Materials and methods

### *Ciona* handling, electroporation, fixing, and imaging

*Ciona robusta (intestinalis* Type A) adults were collected in the San Diego, California area by M-REP Consulting. Specimens were kept in artificial salt water at 34 ppt salinity under constant light. Gametes were obtained from dissected adults and fertilized *in vitro* as previously described (Christiaen et al., 2009b). Zygotes were dechorionated and electroporated as previously described (Christiaen et al., 2009a). Unless precisely specified in the **Supplemental Sequences File,** plasmid amounts electroporated ranged from 35-100 μg per 700 μl of total electroporation volume, except for those encoding Histone 2B (H2B) fusion fluorescent proteins, which were electroporated at much lower dose, around 10-25 μg per 700 μl. Unless otherwise noted, GFP/mCherry/YFP(Venus) proteins are tagged with Unc-76 (Dynes and Ngai, 1998), which is routinely used in *Ciona* to ensure even labeling of cell bodies and axons. Embryos were raised at 20°C unless otherwise specified. Embryos, larvae, and juveniles were fixed in MEM-FA solution (3.7% formaldehyde, 0.5M NaCl, 0.1M MOPS pH 7.4, 1 mM EGTA, 2 mM MgSO4, 0.1% Triton-X100) and washed first in 1X PBS/0.4% Triton-X100/50mM NH4Cl, then in 1X PBS/0.1% Triton-X100 before mounting in 1X PBS/2% DABCO/50% glycerol. Pax3/7 and mCherry immunostains were performed as previously described (Kim et al., 2022) using mouse anti-Pax3/7 antibody graciously provided by Nipam Patel (Davis et al., 2005), rabbit anti-mCherry (BioVision antibody 5993), and Alexa Fluor 555 anti-mouse or rabbit secondary antibodies (Thermo Fisher). Slides were imaged on a Leica DMI8 or DM IL LED inverted epifluorescence microscope with a K3M or DFC3000 G monochrome digital camera. Cell measurements were performed in Leica LAS X software. Statistical tests were performed with the help of GraphPad Prism (see **Supplemental Table 2** for details of all the statistical tests of proportions).

### CRISPR/Cas9 methods

Tissue-specific CRISPR/Cas9-mediated mutagenesis in F0 was performed as generally described (Gandhi et al., 2017; Stolfi et al., 2014). Design of single-chain guide RNAs (sgRNAs) were done in CRISPOR at crispor.tefor.net (Haeussler et al., 2016), selecting for high Doench ’16 and MIT specificity scores, avoiding known SNPs based on the GHOST “Joined 2008” genome assembly and annotation (Satou et al., 2008). Cloning sgRNA expression vectors with the U6 promoter (Nishiyama and Fujiwara, 2008) was performed using oligo annealing/ligation or one-step overlap PCR/in-fusion recombination (Gandhi et al., 2018), or more recently by custom *de novo* synthesis and cloning by Twist Bioscience. Validation of sgRNA efficacies were performed as recently described (Johnson et al., 2023a) by Amplicon-EZ Illumina sequencing of 150-450 bp PCR products amplified from pooled CRISPR-treated larvae, by Genewiz from Azenta Life Sciences (New Jersey, USA). For validation, 75 μg of sgRNA plasmid was co-electroporated with 25 μg of *Eef1a>Cas9* (Stolfi et al., 2014) per 700 μl of total volume. Validation plots are shown in **Supplemental Figure 3**.

### Cell sorting and bulk RNA sequencing

Embryo dissociation and fluorescence-activated cell sorting (FACS) were performed as previously described (Kim et al., 2020), using TagRFP (Merzlyak et al., 2007) to label the desired cells and eGFP to counterselect (see **Supplemental Sequences File** for detailed plasmid mixtures). Two biological replicates for perturbation and negative control conditions (*Fgf8/17/18>Pax3/7* and *Fgf8/17/18>lacZ,* respectively) were raised to 13.5 hours at 18°C and dissociated following established methods (Wang et al., 2018). TagRFP+/eGFP-negative cells were isolated on a BD FACS Aria cell sorter directly into lysis buffer for the RNAqueous Micro RNA extraction kit (Thermo Fisher). RNA was extracted following the kit’s instructions. Following previously published methods (Sharma et al., 2019; Wang et al., 2019), the SMART-seq v4 Ultra Low Input RNA kit (Takara) was used to prepare cDNAs starting from 1.5 µg of total RNA for each sample and sequencing libraries were prepared subsequently prepared using the Ovation Ultralow System V2 (NuGen). Libraries were diluted to 4 nM and pooled with unrelated samples for sequencing with an Illumina NextSeq 500, at 2x75 mid-output. Raw FASTQ files can be accessed at NCBI under accession number PRJNA1056038.

Bulk RNAseq processing and analysis were performed in Galaxy (usegalaxy.org)(Afgan et al., 2022) as previously described (Johnson et al., 2023b). Briefly, raw RNAseq reads were inspected using FastQC Read Quality Reports (Galaxy Version 0.73+galaxy0) and MultiQC (Galaxy Version 1.11+galaxy0). Cutadapt (Galaxy Version 4.0+galaxy0) was used to filter and trim reads, with minimum read length set to 20. Mapping was performed using RNA STAR (Galaxy Version 2.7.8a+galaxy0) with SA pre-indexing string length set to 12, using the “HT” genome assembly and “KY21” gene models from the Ghost Database (http://ghost.zool.kyoto-u.ac.jp/download_ht.html)(Satou et al., 2019; Satou et al., 2022), and counts were obtained using featureCounts (Galaxy Version 2.0.1+galaxy2) with minimum mapping quality per gene set to 10. Differential expression analysis was performed using DESeq2 (Galaxy Version 2.11.40.7+galaxy1). KY21 and older “KH” gene model versions were linked using the Ciona Gene Model Converter (https://github.com/katarzynampiekarz/ciona_gene_model_converter) and this was used to cross-reference results with previously published microarray data on ddN enrichment (Gibboney et al., 2020).

## Results

### Identifying genes downstream of Pax3/7 in the ddNs

It was previously shown that Pax3/7 is sufficient to confer a ddN-like identity to other MG neurons when overexpressed (Stolfi et al., 2011). However, aside from genes encoding other transcription factors such as *Dmbx, Pou4,* and *Lhx1/5,* no other direct or indirect transcriptional targets of Pax3/7 have been identified (Imai et al., 2009). In order to understand the regulation of ddN identity and morphogenesis, we sought to identify candidate effector genes specifically upregulated by Pax3/7 overexpression in the MG. To this end, we carried out fluorescence-activated cell sorting (FACS) to isolate MG progenitor cells from dissociated late tailbud stage 24 embryos (**Figure 1B**). An *Fgf8/17/18>TagRFP* fluorescent reporter plasmid was used to label cells of the A9.30 lineage that gives rise to most of the MG, while the *Fgf8/17/18* promoter was used to overexpress Pax3/7 or a neutral “control” *lacZ* transgene in these cells. Cells were selected by FACS and processed for bulk RNA sequencing (RNAseq). See materials and methods for more experimental details.

Differential gene expression analysis was carried out to identify those transcripts specifically upregulated by *Fgf8/17/18>Pax3/7* in the FACS-isolated MG cells. This identified 536 genes that showed at least 0.6 log2 fold-change significant increase in expression (p<0.05) in response to Pax3/7 overexpression (**Figure 1C, Supplemental Table 1**). To narrow our search, we compared this set to the subset of transcripts enriched in ddNs relative to another MG neuron subtype, MG Interneuron 2 (MGIN2), which had been previously determined by microarray analysis (Gibboney et al., 2020). This resulted in a shorter list of only 43 genes shared between the two datasets (**Figure 1C, Supplemental Table 1**). This represented our list of candidate genes upregulated downstream of Pax3/7 specifically in the ddNs.

A shorter list revealed interesting candidate regulators and effectors of ddN morphogenesis and differentiation from among the larger list of 43 genes (**Figure 1D**). Of these, a few were previously confirmed by *in situ* hybridization to be upregulated in ddNs during their development, including *Saxo, Nckap5, Scna.a,* and *Pou4* (Gibboney et al., 2020). Also significantly upregulated by Pax3/7 was *Fn1-related,* which had been previously detected in the Pax3/7+ sister cells of the ddNs (Gibboney et al., 2020). *Netrin1* was also significantly upregulated by Pax3/7. Although the enrichment of *Netrin1* transcripts in the ddNs had narrowly missed the statistical significance cutoff in the previously published microarray analysis, this gene was subsequently shown by *in situ* hybridization to be expressed in the ddNs and shown to be required for optimal ddN axon guidance across the midline (Gibboney et al., 2020). Curiously, known Pax3/7 targets *Lhx1/5* and *Dmbx* were not signficantly upregulated by Pax3/7 (**Figure 1E**), suggesting that overexpression of Pax3/7 may have activated negative feedback and/or feed-forward regulatory programs in our assay. This suggests that many key regulators and effectors downstream of Pax3/7 are transiently upregulated and/or their expression levels are limited by such feedback/forward mechanisms.

After screening novel candidate *cis*-regulatory sequences potentially controlling the expression of these 43 genes in the ddNs, we obtained new fluorescent reporter plasmids for two of them: *Saxo* and *Ddpep (<u>dd</u>N-expressed <u>pep</u>tide). Saxo* (ANISEED unique gene identifier Cirobu.g00001698) is the sole *Ciona* homolog of human *SAXO1/SAXO2* (Dardaillon et al., 2020), which encode microtubule-stabilizing/bundling proteins. These are part of a larger class of microtubule-binding Saxo proteins that are conserved from unicellular eukaryotes to vertebrates (Cuveillier et al., 2020; Dacheux et al., 2012; Dacheux et al., 2015; Erickson et al., 2023; Gui et al., 2022; Leung et al., 2023). *Saxo* was the top most-enriched gene transcript in the ddNs in our overlap list, and modestly upregulated by Pax3/7 overexpression (**Figure 1D**). A *cis*-regulatory element from the first intron of *Ciona Saxo* was sufficient to drive reporter gene expression in the ddNs, Bipolar Tail Neurons (BTNs), and all epidermal neurons (**Figure 2, Supplemental Figure 1**). Expression in ddNs and BTNs had been previously confirmed by *in situ* hybridization (Gibboney et al., 2020; Kim et al., 2020).

**Figure 2.**
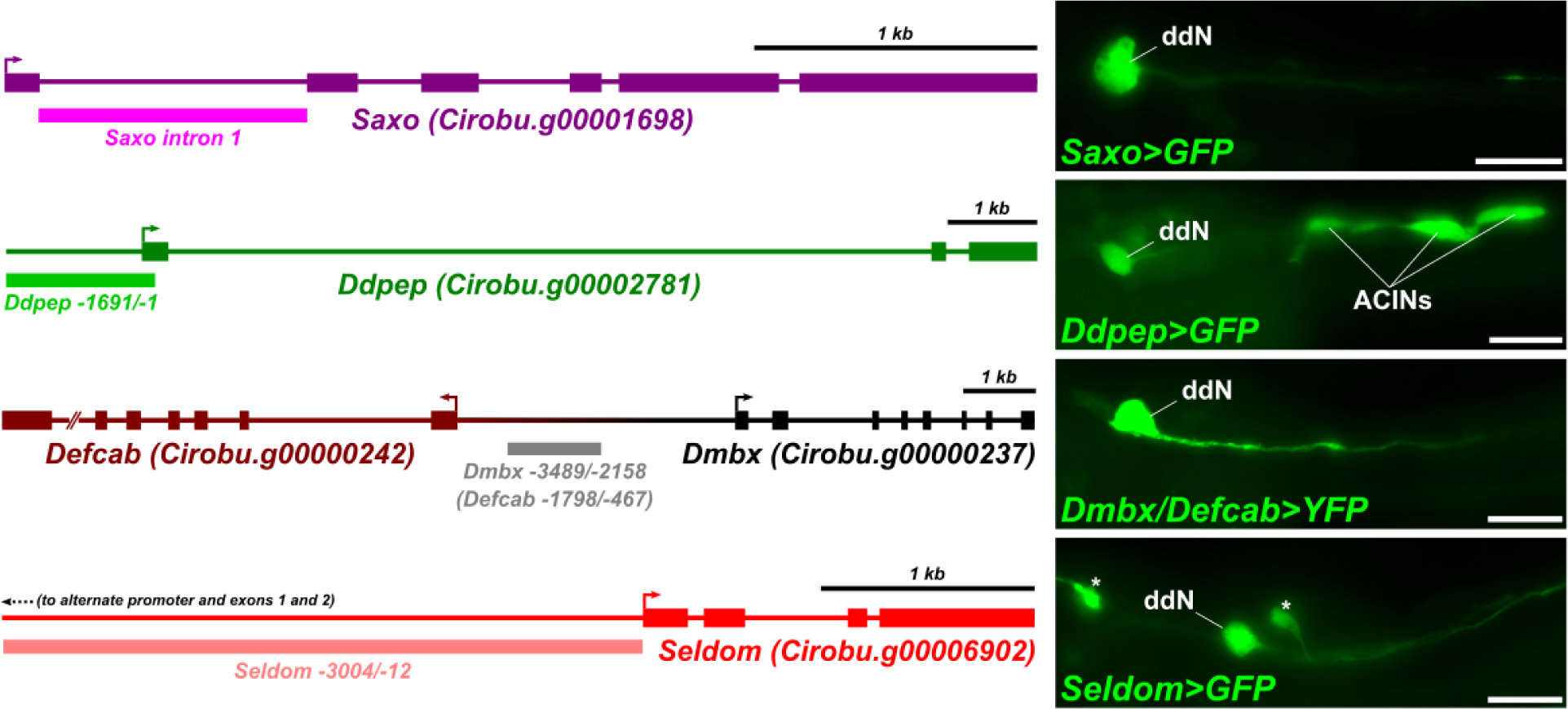
*Cis*-regulatory reporters used in this study. Diagrams of the ddN-expressed genes selected for in-depth study, depicting *cis*-regulatory sequences used to make fluorescent reporter plasmids that mark the ddNs (right panels). All larvae raised at 20°C to 18.5 hpf/stage 28 (*Dmbx/Defcab, Seldom)* or 19 hpf/stage 28 (*Ddpep, Saxo).* All scale bars = 25 µm.

In contrast to the highly conserved *Saxo* gene, *Ddpep* (Cirobu.g00002781) encodes a short peptide that has only been identified in tunicates (Kawada et al., 2011), with no detectable homology or conserved domains/motifs shared with other organisms. Although its transcripts are only moderately enriched in ddNs relative to MGIN2, *Ddpep* was the top gene upregulated by Pax3/7 overexpression, after the pan-neural/muscle gene *Myh.c* (Chiba et al., 2003; Kim et al., 2020). A ∼1.7 kbp fragment immediately 5’ to the *Ddpep* transcribed region was sufficient to drive reporter gene expression in the ddNs (**Figure 2**). Interestingly, *Ddpep>GFP* also labeled the ascending contralateral inhibitory neurons (ACINs) of the tail (**Figure 2**)., whose axons also cross the midline like the ddNs (Horie et al., 2010; Ryan et al., 2016; Takamura et al., 2010).

We thus selected *Saxo* and *Ddpep* reporters for a more in-depth investigation of ddN-specific regulation, in addition to two reporters previously published: *Dmbx/Defcab* and *Seldom.* As was previously shown, *Dmbx* (Cirobu.g00000237) and *Defcab* (*<u>d</u>dN-expressed <u>EFCAB</u> family member,* Cirobu.g00000242) share a common *cis-*regulatory element in their intergenic region that drives expression in ddNs (**Figure 2**)(Gibboney et al., 2020). *Seldom (<u>Sel</u>1 <u>dom</u>ain-containing protein,* Cirobu.g00006902) was previously referred to by its KyotoHoya identifier number (*KH.C4.78*). A *Seldom* reporter was used to label mechanosensory neurons in the papillae (Johnson et al., 2023b), but is also highly expressed in the ddNs, and in assorted neurons in the brain and periphery (**Figure 2**). We reasoned that these four reporters would provide a diverse sampling of *cis*-regulatory logic for further study.

### MG-specific CRISPR-mediated mutagenesis of *Pax3/7*

To verify whether the expression of the selected reporter plasmids in the ddNs depends on Pax3/7, we performed tissue-specific CRISPR/Cas9-mediated mutagenesis in the MG in F0 larvae (**Figure 3A**), as routinely carried out in *Ciona* (Gandhi et al., 2018). We used the *Fgf8/17/18* promoter to drive expression of Cas9 (*Fgf8/17/18>Cas9*) in the A9.30 lineage, which gives rise to most of the neurons in the MG including the ddNs (Cole and Meinertzhagen, 2004; Imai et al., 2009; Stolfi and Levine, 2011), and previously published single-chain guide RNA (sgRNA) expression plasmids targeting *Pax3/7* (Kim et al., 2022). Previously, this MG-specific CRISPR mutagenesis of *Pax3/7* resulted in significant loss of the *Dmbx/Defcab* reporter expression in the ddNs (Kim et al., 2022). Here we assayed the expression of the three remaining ddN reporters (*Saxo, Ddpep,* and *Seldom*), comparing between *Pax3/7* CRISPR and “negative control” CRISPR larvae electroporated instead with an sgRNA that does not target any known *Ciona* sequences (Stolfi et al., 2014). As expected, CRISPR/Cas9-mediated mutagenesis of *Pax3/7* in the MG reduced the frequency of expression for all three reporters (**Figure 3B,D**). This suggests all are indeed downstream of Pax3/7 in the ddNs.

**Figure 3.**
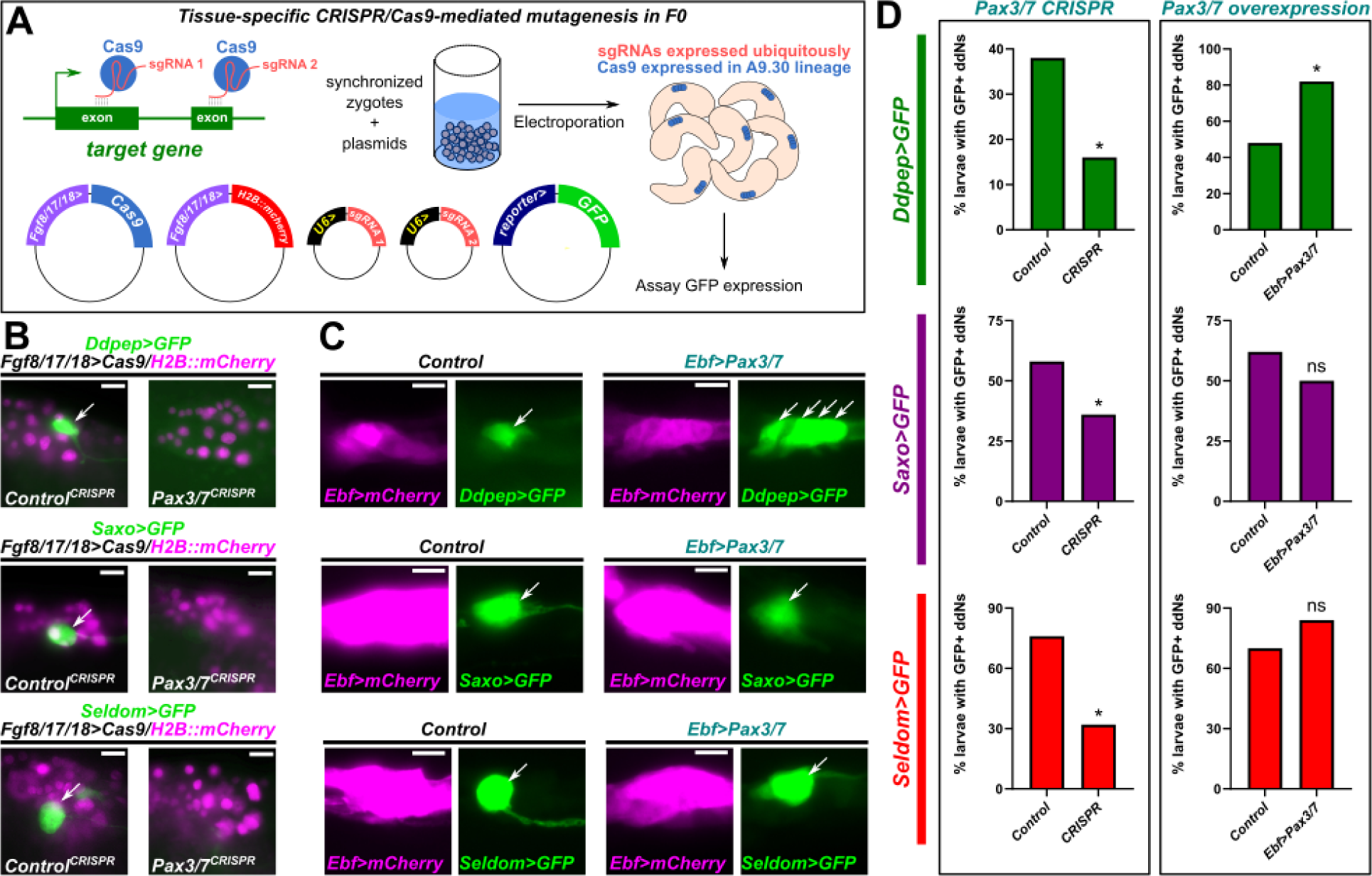
*Pax3/7* regulates gene expression in the ddNs. A) Schematic diagram of tissue-specific CRISPR/Cas9-mediated mutagenesis as carried out in this study. In this example, the *Fgf8/17/18* promoter is used to drive expression of Cas9 in the A9.30 cell lineage that gives rise to the ddN. B) Representative images of the effects of CRISPR/Cas9-mediated mutagenesis of *Pax3/7* on *Ddpep, Saxo,* and *Seldom* reporter expression in the ddNs (arrows). Developmental times are 17 hpf/stage 27 for *Ddpep*, 18 hpf/stage 28 for *Saxo* and *Seldom.* C) Effects of Pax3/7 overexpression in the motor ganglion, assaying the same reporters as in (B). Pax3/7 overexpression resulted in ectopic expression of *Ddpep* reporter (extra arrows) but not for *Saxo* or *Seldom* reporters. D) Scoring frequency of reporter expression in *Pax3/7* CRISPR and overexpression experiments, n= 50 for all conditions. Only larvae expressing *Fgf8/17/18>H2B::mCherry* or *Ebf>mCherry* reporters were scored, as a way to exclude unelectroporated animals from the analysis. Developmental times are 17.5 hpf/stage 27 for *Saxo* and *Seldom,* and 19.5 hpf/stage 29 for *Ddpep.* All larvae raised at 20°C. All scale bars = 10 μm. * p <0.05, ns = not significant. See **Supplemental Table 2** for all statistical tests.

Previously, Pax3/7 was shown to be sufficient to activate ectopic expression of the *Dmbx/Defcab* reporter throughout the MG (Serrano-Saiz et al., 2018). To test whether Pax3/7 is also sufficient to activate our new ddN reporters, we used the *Ebf* promoter to drive ectopic expression of Pax3/7 in all neurons of the MG (Kratsios et al., 2012). Interestingly, this resulted in significantly higher frequency expression of the *Ddpep>GFP*, but not that of *Saxo>GFP* or *Seldom>GFP* (**Figure 3C,D**). For *Ddpep>GFP,* ectopic reporter expression could be seen in additional MG cells (**Figure 3C**). In contrast, *Saxo* reporter expression was modestly downregulated by Pax3/7 overexpression (**Figure 3D**). This mildly repressive effect was statistically significant when using the *Fgf8/17/18* promoter to overexpress Pax3/7 more specifically in the MG (**Supplemental Figure 2**). This suggests that there may be an indirect regulatory pathway (with potentially an incoherent/negative feed-forward loop inserted) between Pax3/7 and *Saxo* and *Seldom*.

### Pou4 is required for expression of *Saxo* and *Seldom*

Which transcription factor(s) might be regulating *Saxo* and *Seldom* downstream of Pax3/7? One obvious candidate was the POU-homeodomain transcription factor Pou4, which is expressed specifically in the ddNs downstream of Pax3/7 (Imai et al., 2009). *Pou4* is the sole *Ciona* homolog of the *POU4/Brn3* class of genes encoding highly conserved transcription factors that regulate neuronal specification and differentiation in various animal species (Chacha et al., 2022; Collum et al., 1992; Gold et al., 2014; Leyva-Díaz et al., 2020; Ninkina et al., 1993; Serrano-Saiz et al., 2018; Tang et al., 2013; Tournière et al., 2020). Pou4 was already shown to be required for *Seldom* reporter expression in *Ciona* papilla neurons (PNs)(Johnson et al., 2023b), while the *Saxo* reporter is expressed in all neuron types that also express *Pou4* (Candiani et al., 2005). Therefore we reasoned that Pou4 was a likely candidate to regulate *Seldom* and *Saxo* in the ddNs as well.

As expected, MG-specific CRISPR/Cas9-mediated mutagenesis of *Pou4* abolished the ddN-specific expression of both the *Saxo* and *Seldom* reporters in a significant proportion of larvae (**Figure 4A**). In contrast, overexpression of Pou4 using the *Fgf8/17/18* promoter resulted in strong ectopic expression of both reporters throughout the MG in all electroporated embryos (**Figure 4B**). We used the *Fgf8/17/18* promoter in this case because the *Ebf>mCherry* reporter appeared to be repressed by Pou4, preventing accurate scoring of expression (data not shown). On the other hand, Pou4 was not required for *Dmbx/Defcab* or *Ddpep* reporter expression (**Figure 4A**), nor sufficient to induce or increase their expression (**Figure 4C**). We therefore conclude that Pou4 is necessary and sufficient for the expression of *Seldom* and *Saxo* in the ddNs, but not that of *Dmbx, Defcab,* or *Ddpep*.

**Figure 4.**
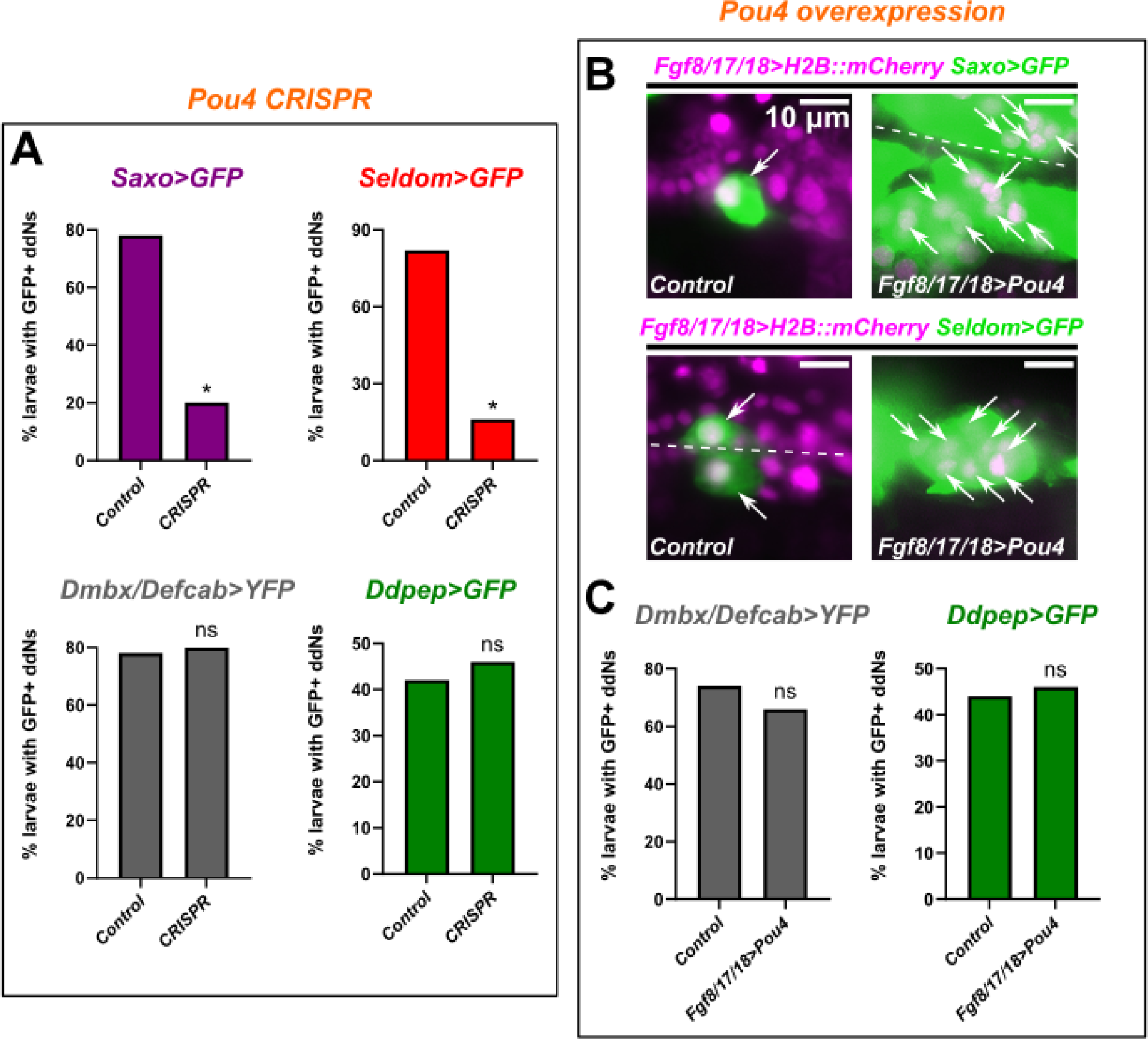
Regulation of *Saxo* and *Seldom* in ddNs by Pou4. A) Plots showing frequency of *Saxo, Seldom, Dmbx/Defcab,* and *Ddpep* reporter plasmids in the ddNs of H2B::mCherry+ larvae (17.5-18 hpf/stage 27-28) subjected to CRISPR/Cas9-mediated disruption of *Pou4* in the MG or in a parallel negative control condition. B) Representative images of larvae in which Pou4 was overexpressed ectopically throughout the MG (A9.30 cell lineage) using the *Fgf8/17/18* promoter. Pou4 overexpression results in massive ectopic activation (extra arrows) of *Seldom* and *Saxo* reporters in all larvae, but not *Dmbx/Defcab* or *Ddpep* reporters. Single or paired arrows always indicate a ddN or ddN left/right pair as in the wild type. Dashed line indicates the neural tube midline. Larvae fixed at 19.5 hpf/stage 29 (*Saxo*, *Seldom,* and *Dmbx/Defcab*) and 17.5 hpf/stage 27 *(Ddpep*). C) Scoring plots showing lack of increased activation of *Dmbx/Defcab* and *Ddpep* reporters upon electroporation of *Fgf8/17/18>Pou4* (only H2B::mCherry+ larvae scored). See text for experimental details and **Supplemental Table 2** for all statistical tests. * p < 0.0001, ns = not significant. All larvae raised at 20°C, all n = 50 except *Saxo* reporter in *Pou4* CRISPR (n = 60), all scale bars = 10 μm.

### Lhx1/5 overexpression results in ectopic *Dmbx/Defcab* activation

Similarly to Pou4, the LIM-homeodomain transcription factor Lhx1/5 is also expressed in the ddNs downstream of Pax3/7 (Imai et al., 2009). *Lhx1/5* is the *Ciona* ortholog of vertebrate *Lhx1* and *Lhx5* genes, homologs of which play significant roles in neurodevelopment throughout metazoa (Avraham et al., 2009; Currie and Pearson, 2013; Pillai et al., 2007; Serrano-Saiz et al., 2018; Sheng et al., 1997; Taira et al., 1992; Zhao et al., 1999). To test whether Lhx1/5 might be regulating downstream effector gene expression in the ddNs, we sought to disrupt it in the MG with newly validated *Lhx1/5-*targeting sgRNAs (**Supplemental Figure 3**). Although we observed high indel rates elicited by our *Lhx1/5-*targeting sgRNAs, the expression frequency of our ddN reporters was not profoundly affected by *Lhx1/5* CRISPR (**Figure 5A**). Thus, we were unable to confirm tissue-specific disruption of *Lhx1/5* and a requirement for this gene in the absence of a clear phenotype.

**Figure 5.**
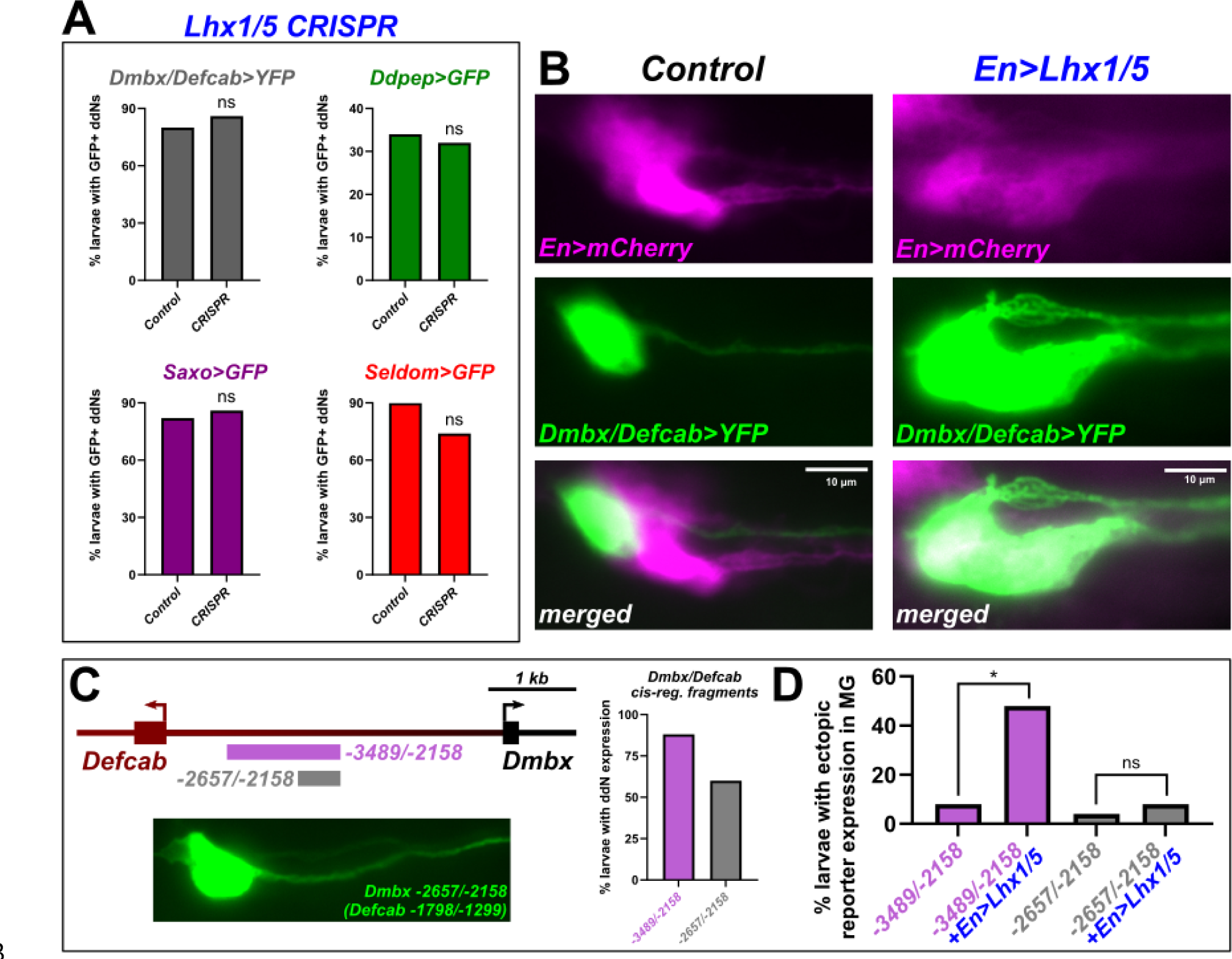
Lhx1/5 upregulates the activity of a *Dmbx/Defcab cis*-regulatory element that is dispensable for precise gene expression in ddNs. A) Plots showing modest or no reduction of ddN reporter plasmid expression upon MG-specific CRISPR-based disruption of *Lhx1/5* in mCherry+ larvae (18.5 hpf/stage 28). B) Overexpression of Lhx1/5 in the whole anterior MG using the *En* promoter results in ectopic expression of the standard *Dmbx/Defcab* reporter used in previous figures Quantification shown in panel D, alongside the smaller *Dmbx/Defcab* reporter (see text for details). Larvae fixed at 20 hpf/stage 29. C) Top: diagram showing large (-3489/-2158) and small (-2657/-2158) shared *Dmbx/Defcab* intergenic fragments tested. Coordinates given relative to start codon of *Dmbx* transcript. See text and **Supplemental Sequences File** for details. Bottom: image showing expression of a GFP reporter plasmid using the smaller fragment, labeling the ddN at 20 hpf/stage 29. Right: quantification of GFP expression frequency with reporter plasmids using the large and small fragments, at 20 hpf/stage 29. D) Plot showing the effect of *En>Lhx1/5* in increasing the frequency of ectopic reporter expression in the MG, but only with the larger *Dmbx/Defcab* fragment, indicating the smaller fragment does not respond to Lhx1/5 overexpression. All larvae at 20 hpf/stage 29, raised at 20°C. All n = 50, all scale bars = 10 µm. See text for experimental details, and **Supplemental Table 2** for all statistical tests. * p < 0.0001, ns = not significant.

Conversely, we overexpressed Lhx1/5 in the anterior MG (from which the ddNs are specified) using the *Engrailed (En)* promoter (Stolfi et al., 2011). We used this promoter because broader overexpression with Lhx1/5 using the *Ebf* or *Fgf8/17/18* drivers was producing non-specific developmental defects that made it difficult to score ddN expression (data not shown). Upon overexpression of Lhx1/5, we observed ectopic expression of the *Dmbx/Defcab* reporter in the MG (**Figure 5B**). Although we cannot not rule out a requirement for Lhx1/5 in activating transcription of *Dmbx* or *Defcab,* we hypothesized that Lhx1/5 might be playing an auxiliary role instead. Consistent with this idea, we found that Lhx1/5 overexpression caused ectopic expression of the larger *Dmbx/Defcab* regulatory element (*Dmbx -3489/-2158)*, but not that of a smaller region (*Dmbx -2657/-2158)* centered on the minimal, Pax3/7-dependent activation element previously identified (Stolfi et al., 2011)(**Figure 5C,D**). Although we observed higher frequency of normal reporter plasmid expression with the larger fragment compared to the smaller fragment, the smaller fragment lacking Lhx1/5-responsive element is still active in the ddNs (**Figure 5C**). In contrast, *En>Lhx1/5* did not result in ectopic expression of any of the other ddN reporters tested (**Supplemental Figure 4**) Taken together, these results indicate that Lhx1/5 is likely not required for initial activation of *Dmbx* or *Defcab,* but rather augments or maintains their expression through a separate *cis*-regulatory module adjacent to the activation module.

### Pax3/7 is required for *Ddpep* expression in ACINs

As was mentioned above, some of the ddN reporters are also expressed in other neuron types in the *Ciona.* For instance, *Seldom* is expressed in PNs and BTNs, *Saxo* is expressed in neurons of the peripheral nervous system: namely PNs, BTNs, and various epidermal neurons, and *Ddpep* is expressed in ACINs. We therefore asked if the regulation of these reporters in these other neuron types was similar to their regulation in ddNs. Because Pou4 was already shown to be required for activation of the *Seldom* reporter in PNs (Johnson et al., 2023b), we focused on *Ddpep* and *Saxo*.

Because Pax3/7, but not Pou4 or Lhx1/5, was sufficient and necessary for *Ddpep* reporter expression in the ddNs, we hypothesized that this regulatory logic might also operate in the ACINs. Immunostaining with a pan-bilaterian Pax3/7 antibody (Davis et al., 2005) and *Pax3/7* reporter plasmids (Horie et al., 2018; Kim et al., 2022) suggested that ACINs also express Pax3/7 (**Supplemental Figure 5A,B**). Because ACINs derive from the A9.29 cell lineages (Nishitsuji et al., 2012), we used the *Ephrin A.b* promoter (**Supplemental Figure 5C**) to drive expression of Cas9 in these cells to disrupt *Pax3/7* in ACIN progenitors. Indeed, *Ddpep* reporter expression in the ACINs was largely abolished with *Pax3/7* CRISPR in these cells (**Figure 6A,B**). We conclude that Pax3/7 regulates *Ddpep* expression not only in the ddNs, but in ACINs as well.

**Figure 6.**
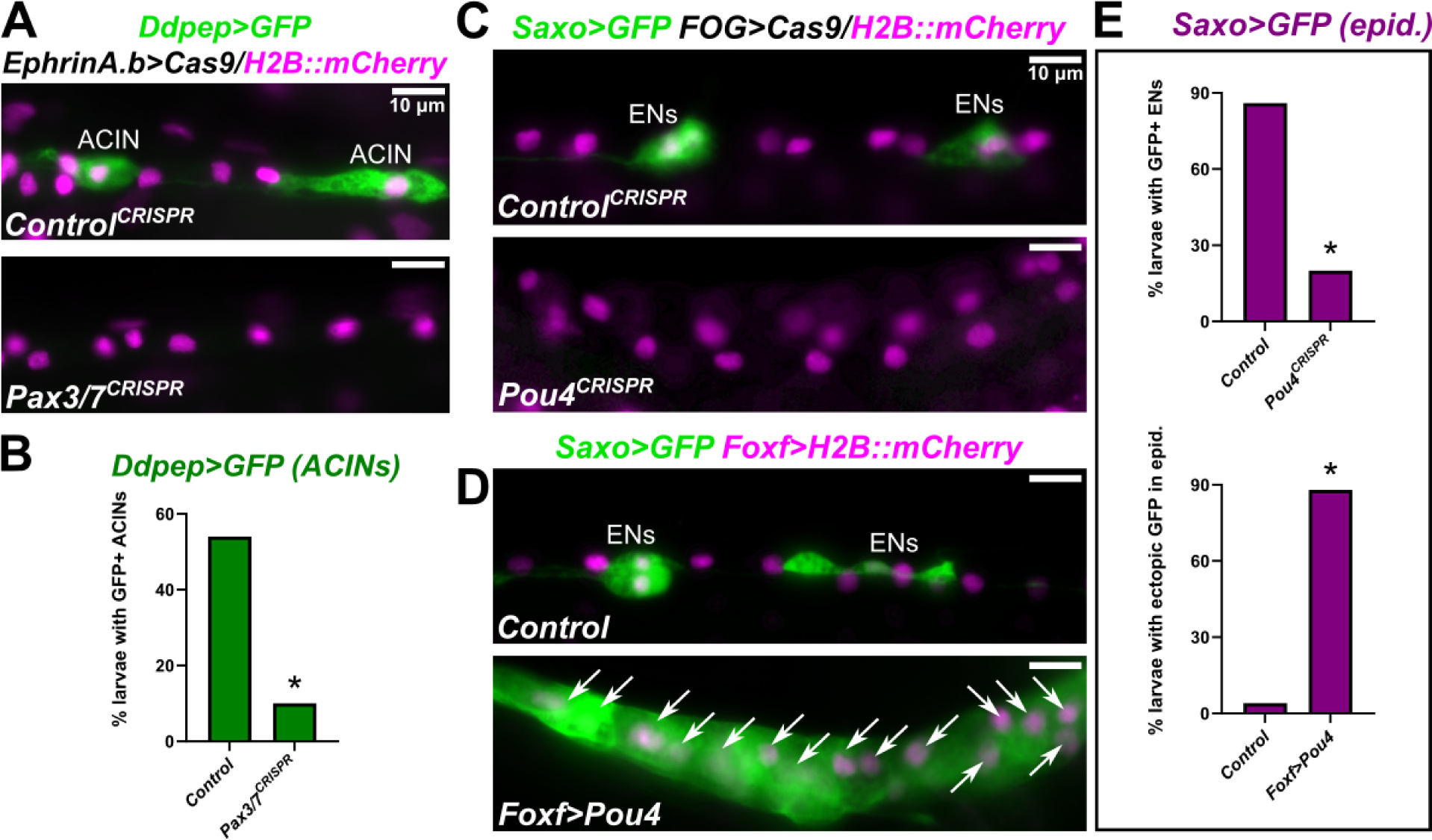
Pax3/7 is required for *Ddpep* expression in ACINs and Pou4 is required for *Saxo* expression in epidermal neurons. A) Loss of *Ddpep* reporter expression in the ascending contralateral inhibitory neurons (ACINs) at 18 hpf/stage 28 by targeted CRISPR/Cas9-mediated mutagenesis of *Pax3/7* using the *EphrinA.b* promoter to drive expression of Cas9 in the ACIN lineages. B) Plot comparing frequency of *Ddpep* reporter expression between *Pax3/7* CRISPR and negative control conditions in H2B::mCherry+ larvae as depicted in panel A. C) Loss of *Saxo* reporter plasmid in epidermal neurons (ENs) of the tail at 19 hpf/stage 28 by targeted CRISPR/Cas9-mediated mutagenesis of *Pou4* using the *FOG* promoter to drive expression of Cas9 in animal pole lineages, which give rise to all epidermal neuron progenitors. D) Overexpression of Pou4 in the epidermis using the *Foxf* promoter results in ectopic expression of the *Saxo* reporter (arrows) in all larvae at 19 hpf/stage 28. E) Plots showing frequency of GFP expression in H2B::mCherry+ ENs (top) and ectopic GFP expression in H2B::mCherry+ epidermis (bottom), in the conditions depicted in panels C and D, respectively. All larvae raised at 20C, all n = 50, all scale bars = 10 µm. See text for experimental details and **Supplemental Table 2** for statistical tests. p < 0.0001.

### Pou4 activates *Saxo* expression in the peripheral nervous system

Next, we investigated the regulation of *Saxo* in peripheral neurons (PNs, BTNs, and epidermal neurons). Because *Pou4* is also expressed in all peripheral neurons (Candiani et al., 2005; Chacha et al., 2022; Tang et al., 2013), we sought to test whether the regulatory connection between Pou4 and *Saxo* is truly ddN-specific or more widespread in the *Ciona* larval nervous system. To this end, we first carried out CRISPR/Cas9-mediated mutagenesis of *Pou4* in the epidermis, using the *FOG* promoter (Rothbächer et al., 2007) to drive Cas9 in animal pole lineages. This significantly reduced the frequency of *Saxo* reporter expression in epidermal neurons (**Figure 6C,E**). Conversely, overexpression of Pou4 throughout the epidermis using the *Foxf* promoter (Beh et al., 2007) resulted in strong ectopic *Saxo* reporter expression throughout this tissue (**Figure 6D,E**). From these results, we conclude that Pou4 is sufficient and necessary for the expression of *Saxo* in neurons of the epidermis.

Because of the strong effect of Pou4 gain- or loss-of-function on *Saxo* reporter expression in all tissues assayed, we predicted that Pou4 directly activates this expression through binding to specific sites in our *Saxo cis*-regulatory element (located in the first intron). We searched for predicted Pou4 binding sites using JASPAR (Rauluseviciute et al., 2023) and looked for sequences conserved in the related *Ciona savignyi* species. This revealed two conserved potential Pou4 binding sites (**Figure 7A**). To test these sites, we generated *Saxo>GFP* reporter plasmids bearing point mutations predicted to disrupt potential binding by Pou4. Co-electroporating these with the wild-type *Saxo>mCherry* reporter showed that mutating the first site (mPou4 site 1) did not discernably alter expression (**Figure 7B**). However, GFP expression was nearly abolished by mutating the second site (mPou4 site 2), while mutating both further reduced reporter activity (**Figure 7B**). These data suggest that Pou4 might directly activate the transcription of *Saxo* through binding to a specific site in its intronic *cis*-regulatory element.

**Figure 7.**
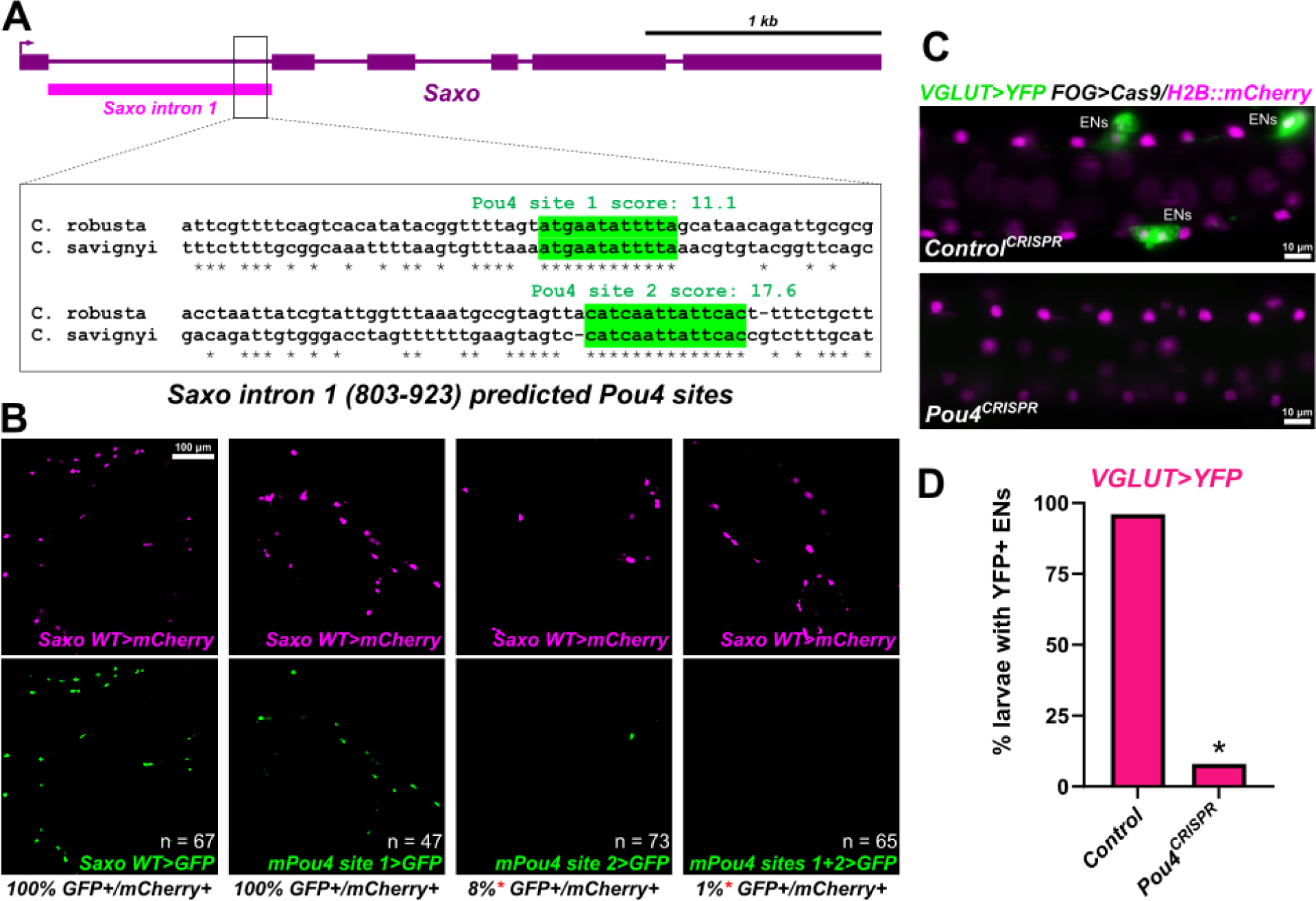
Regulation of *Saxo* and *VGLUT* expression in epidermal neurons by Pou4. A) Diagram of intronic *Saxo cis*-regulatory element, with region of conservation between *Ciona robusta* and *Ciona savignyi* magnified in detail. Putative Pou4-binding sites with scores predicted by JASPAR highlighted in green. B) Representative images of larvae (19 hpf/stage 28) co-electoporated with an mCherry reporter plasmid using the wild type (WT) *Saxo* regulatory element and GFP reporter plasmids with WT or mutated *Saxo* elements carrying point mutations predicted to disrupt the binding of Pou4 (“mPou4” mutations) at one or both sites. Quantification below each panel represents the frequency of GFP co-expression in each mCherry-expressing neuron. The number of neurons scored is represented as “n” in each GFP panel. C) Animal pole lineage-specific disruption of *Pou4* results in frequent loss of *VGLUT* reporter in epidermal neurons (ENs) at 18.5 hpf/stage 28. D) Plot of *Pou4* CRISPR effect on *VGLUT* reporter expression as represented in panel C, n = 50 (H2B::mCherry+ larvae) for either condition. All scale bars = 10 μm unless otherwise specified, all larvae raised at 20°C, see text for experimental details and **Supplemental Table 2** for statistical tests. * p < 0.0001, ns = not significant.

Although Pou4 factors play many conserved roles in neuronal specification and differentiation (Leyva-Díaz et al., 2020), no data on the regulation of *Saxo* by Pou4 in other organisms. However, we noticed that *Saxo* reporter expression in the peripheral nervous system greatly overlaps that of *Slc17a6/7/8* (also known as *Vesicular glutamate transporter,* or *VGLUT),* as all epidermal neurons in the *Ciona* larva are glutamatergic (Horie et al., 2008). As part of their complex regulatory roles, Pou4 factors also activate the expression of *VGLUT* homologs in *C. elegans* and mouse (Serrano-Saiz et al., 2018). To test if regulation of *VGLUT* by Pou4 is conserved in *Ciona,* we looked at *VGLUT* reporter expression in epidermal *Pou4* CRISPR larvae as above. Near complete loss of *VGLUT* reporter expression was observed in the *Pou4* CRISPR condition (**Figure 7C,D**), demonstrating that Pou4 performs an evolutionarily conserved role of specifying glutamatergic neuron identity in the *Ciona* larva. Interestingly, the ddNs are not glutamatergic and do not express *VGLUT*, suggesting that the Pou4-dependent *cis*-regulatory logic for *VGLUT* is more complex than that for *Saxo*.

### Ddpep overexpression misroutes MG axons

Although we were able to identify more precisely the regulatory mechanisms shaping the transcriptional profile of the ddNs, not much is known about the functions of downstream genes that do not encode transcription factors, i.e. “effector genes”. In the ddNs, a role for *Defcab* promoting axon growth was shown previously, while *Netrin1* was implicated in polarizing ddNs (Gibboney et al., 2020). Here we investigated the potential functions of two of the genes whose regulation we studied in detail above: *Ddpep* and *Saxo,* as well as a third gene that we identified by RNAseq as being strongly upregulated by Pax3/7 but for which we were not able to generate a reporter plasmid: *Nckap5* (Cirobu.g00009755).

*Ddpep* encodes a relatively short, secreted polypeptide sequence (198 a.a.) that is processed into smaller peptides (Kawada et al., 2011). Although Ddpep orthologs are found in other tunicate species (**Supplemental Figure 6**), we have not been able to identify any homologs outside the tunicates. The expression of *Ddpep* in both ddNs and ACINs is interesting because these two are the only neuron types in the MG whose axons cross the midline (Ryan et al., 2016). It has been suggested that *Ddpep* encodes a neuropeptide (Kawada et al., 2011), but we hypothesize that Ddpep might act as an autocrine cue to direct axon growth towards and/or across the midline, like how ddN-expressed Netrin1 has been proposed to act (Gibboney et al., 2020). This hypothesis was supported by the observation that Ddpep protein tagged with GFP (Ddpep::GFP) accumulates on the surface of the ddN axon growth cones as they begin to decussate (**Figure 8A**). Furthermore, we also detected *Ddpep* expression in the posterior tip of the notochord by *in situ* hybridization (**Supplemental Figure 7A**), which was recapitulated by a GFP reporter plasmid utilizing an intronic *cis*-regulatory element from the first intron of *Ddpep* (**Supplemental Figure 7B**). Its localized expression in notochord cells of the tail tip supports a putative function for Ddpep in axon guidance and not as a neurotransmitter.

**Figure 8.**
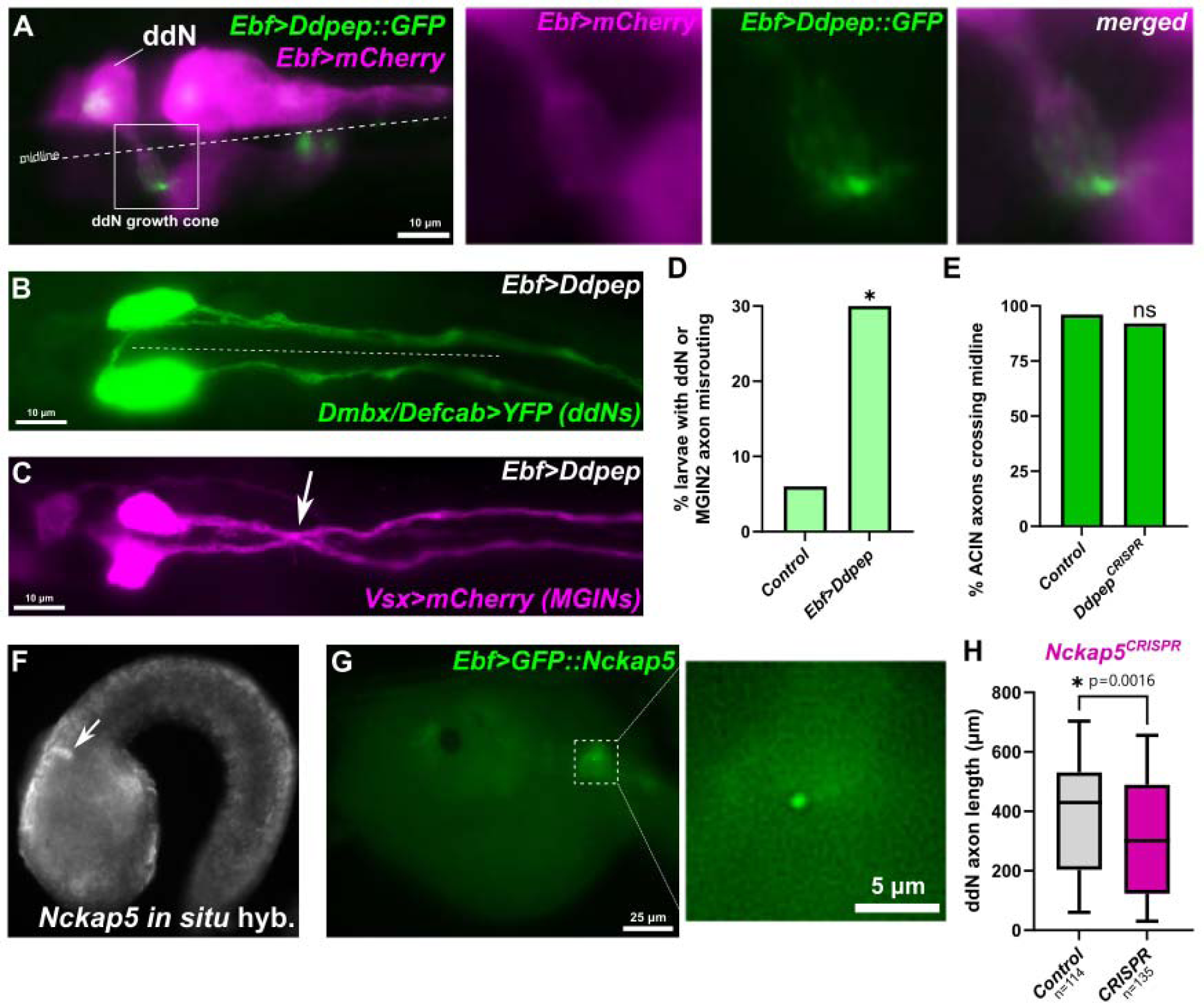
Testing the functions of effector genes in the ddNs. A) Mid-tailbud embryo at 12.5 hpf/stage 21 expressing Ddpep::GFP in the ddNs and other neurons. Inset shows Ddpep:GFP accumulation at growth cone tip of a ddN axon as it crosses the midline. B) Example of ddN axon trajectory defects in the ddNs upon overexpression of Ddpep. The axons of the ddNs are avoiding the midline (dashed line), instead of crossing over like normally. Note also a second axon from the left ddN projecting anteriorly then crossing the midline and turning posteriorly. C) Example of MGIN2 axon trajectory defects upon overexpression of Ddpep. Arrow indicates MGIN2 axons crossing or following the midline, when normally they remain ipsilateral. Both larvae in panels B and C were fixed at 18.5 hpf/stage 28. D) Plot of frequency of abnormal axon trajectories observed in ddNs or MGIN2s upon overexpression of Ddpep as in panels B and C, and a negative control. For each condition, n = 50, * p = 0.0033. E) Scoring of ACIN axon crossing phenotype upon CRISPR/Cas9-mediated mutagenesis of *Ddpep,* showing no significant effect. For either condition, n = 50. F) Fluorescent, whole-mount *in situ* mRNA hybridization showing *Nckap5* expressed in the ddN (arrow). Same embryo (12 hpf/stage 24), different objective as in Gibboney et al. 2020. G) GFP::Nckap5 localization in one small spot in an unidentified MG neuron. The spot is likely around the centrosome, based on known localization of Nckap5 orthologs in human cells. H) Quantification of axon lengths measured in larvae at 18 hpf/stage 28, comparing between a negative CRISPR control condition and MG-specific CRISPR-mediated mutagenesis of *Nckap5.* Number of axons measured indicated by “n” under plot, p = 0.0016 by two-tailed Mann Whitney test. All animals raised at 20°C, all scale bars = 10 µm unless otherwise specified. See text and supplemental sequence file for experimental details and **Supplemental Table 2** for statistical test information.

To disrupt *Ddpep* by CRISPR/Cas9 we designed and validated sgRNAs targeting this gene (**Supplemental Figure 3**). However, MG-specific CRISPR mutagenesis of *Ddpep* did not result in any discernible ddN defects (data not shown). *Ddpep* was also targeted by CRISPR/Cas9 in the ACIN lineage, but ACIN axons still crossed the midline at rates comparable to those in the negative control (**Supplemental Figure 7C**). However, when we overexpressed Ddpep throughout the MG (*Ebf>Ddpep),* this caused an increase in misguided axon trajectories for ddNs and MGIN2s, as visualized by *Dmbx/Defcab* and *Vsx* reporter plasmids respectively (**Figure 8B-D**). More precisely, ddN axons were observed to avoid crossing the midline more frequently, while MGIN2 axons crossed or abutted the midline more often, though other axon guidance defects were also observed (data not shown). Based on these results, we speculate that Ddpep functions as a tunicate-specific autocrine axon guidance cue to promote midline crossing in ddN and ACIN axon growth cones. Although Ddpep is sufficient to misroute axons when overexpressed, we were unable to show that it is necessary for ddN or ACIN axon guidance. It is possible that Ddpep acts redundantly in parallel to other cues (e.g. Netrin1) to provide robustness to axon polarization.

### Testing Saxo function in ddNs and papilla neurons

SAXO (formerly known as FAM154) proteins belong to a larger family of STOP/MAP6-related proteins that localize to centrosomes and cilia to mediate stabilization of microtubules through their Mn domains (Dacheux et al., 2015; Erickson et al., 2023; Leung et al., 2023). Because we had previously demonstrated a role for the centrosome-localized Defcab in ddN axon outgrowth, we hypothesized that Saxo may be playing a similar role as well. However, MG-specific CRISPR mutagenesis of *Saxo* using validated sgRNAs did not affect axon length or trajectory (**Supplemental Figure 8A**). Because Saxo is also expressed in the ciliated mechanosensory PNs of the papillae, we hypothesized that it might play a role in PN development or mechanotransduction. Knocking out *Pou4* in the papillae abrogates PN differentiation and mechanically stimulated induction of metamorphosis (Sakamoto et al., 2022). To test whether *Saxo* is required for PN function, we targeted it using CRISPR specifically in the papilla territory with *Foxc>Cas9* (Johnson et al., 2023b). However, *Saxo* CRISPR did not result in a significant reduction in the proportion of larvae undergoing metamorphosis, unlike eliminating the PNs completely with *Pou4* CRISPR (**Supplemental Figure 8B-D**). Based on these data, we have yet to identify a function for Saxo in any *Ciona* neurons.

### Nckap5 is a centrosome-enriched effector of ddN axon growth

From the list of ddN-enriched genes upregulated by Pax3/7, we found a conserved gene encoding the *Ciona* ortholog of another class of known centrosome-enriched, microtubule-stabilizing proteins: Nckap5 (Cirobu.g00009755). Although *Nckap5* expression in the ddNs was previously confirmed by *in situ* hybridization (**Figure 8E**)(Gibboney et al., 2020), we were unable to identify any *cis*-regulatory sequences that control this expression. *Ciona Nckap5* is orthologous to human *NCKAP5* and *NCKAP5L*, the latter encoding a protein also known as Cep169 that bundles and stabilize microtubules at centrosomes (Mori et al., 2015a; Mori et al., 2015b). We found that a *Ciona* Nckap5::GFP fusion localized to the centrosome in neurons (**Figure 8F**), suggesting a potentially conserved function. To test whether *Nckap5* plays a role in ddN axon growth, we designed and validated sgRNAs targeting it for MG-specific CRISPR/Cas9 mutagenesis (**Supplemental Figure 3**). We noticed that *Nckap5* CRISPR had a detrimental effect on ddN axon growth, as quantitative comparison revealed a significant reduction in average ddN axon length in *Nckap5* CRISPR larvae relative to control larvae (**Figure 8G**), suggesting this gene may be required for optimal ddN axon outgrowth, similar to *Defcab,* which also encodes a protein that localizes to the centrosome (Gibboney et al., 2020).

## Discussion

Here we have partially dissected the GRN controlling the specification, differentiation, and morphogenesis of tunicate ddNs, which may be homologous to vertebrate Mauthner cells. We have revealed that this network consists of different nodes or branches downstream of the ddN fate-determining transcription factor Pax3/7 (**Figure 9**). Pax3/7 is required for the expression of other transcription factors, namely Pou4, Lhx1/5, and Dmbx. These in turn act in different ways to regulate several aspects of ddN development. First, Pax3/7 and Pou4 activate the transcription of multiple effector genes including *Ddpep, Saxo, Seldom, Defcab,* and possibly *Netrin1* and *Nckap5.* In contrast, Lhx1/5 does not appear to be required for the expression of any of the effector genes surveyed, but might help maintain optimal transcript levels of *Dmbx* and *Defcab* through a shared *cis*-regulatory element that is not required for initial activation of either gene. Finally, Dmbx was previously shown to promote ddN differentiation through transcriptional repression of unknown targets (Stolfi et al., 2011).

**Figure 9.**
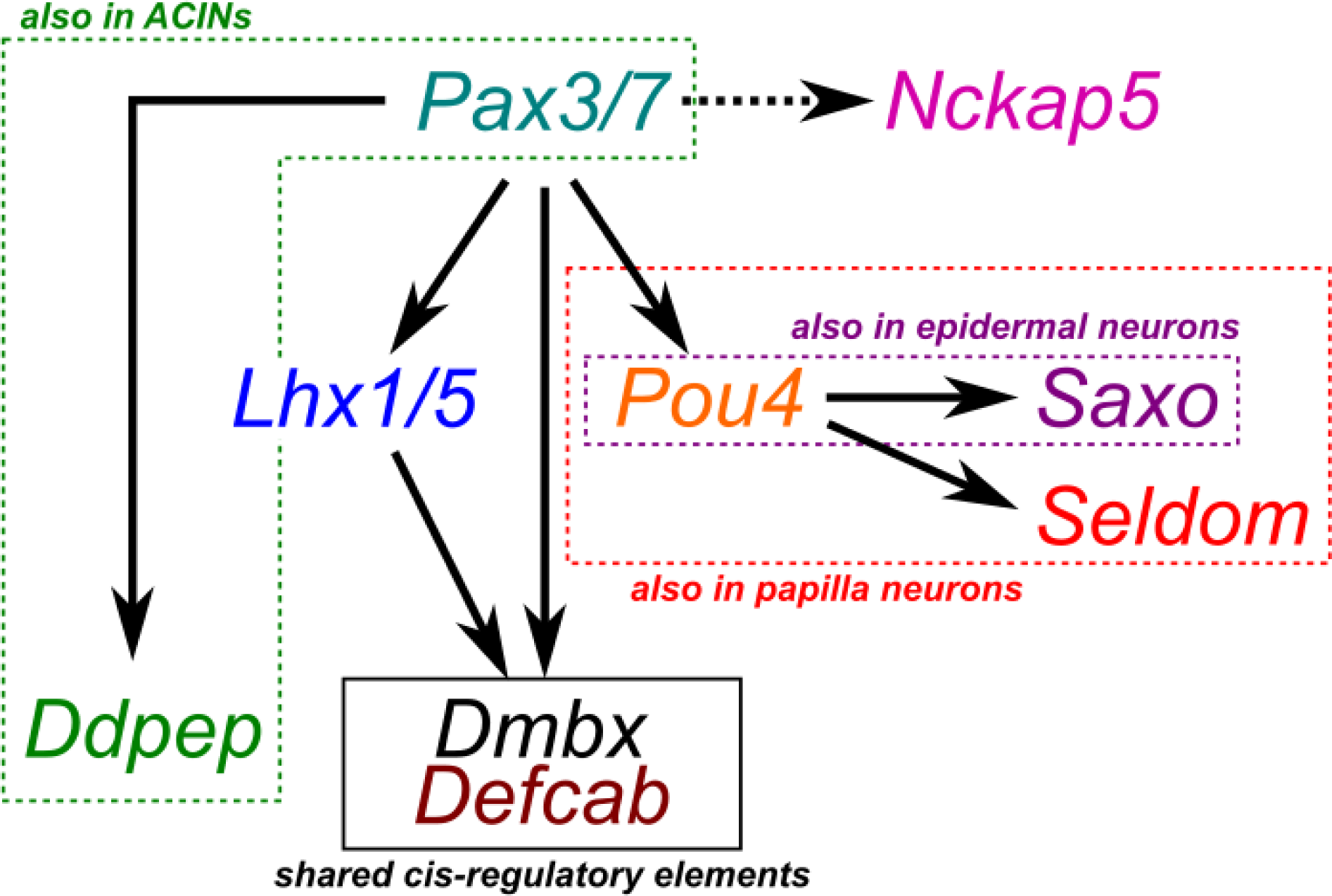
A summary diagram of a ddN gene regulatory network. Diagram for a ddN-specific gene regulatory network downstream of Pax3/7. Some branches are shared with different neuron types, such as ACINs, epidermal neurons, or papilla neurons (a sub-type of epidermal neurons). Dashed arrow indicates regulatory connection supported by RNAseq data, but not validated by reporter plasmids or *in situ* mRNA hybridization.

Interestingly, orthologs of *Pax3/7* and its major regulatory targets (*Lhx1/5, Pou4,* and *Dmbx*) are also crucial for specification of cranial neural crest cells in vertebrates (Simoes-Costa and Bronner, 2016). Although the precise regulatory relationships between these factors appears to be different between vertebrates and *Ciona,* this may hint at a deeply conserved regulatory “kernel” might have been key to the evolutionary diversification of neural crest derivatives along the rostral-caudal axis of vertebrates (Martik et al., 2019; Rothstein et al., 2018).

Our data also suggest a role for the tunicate-specific secreted molecule Ddpep in the regulation of axon guidance in ddNs and possibly in ACINs. Pax3/7 is required for expression of *Ddpep* in both neuron types, which are also the only MG neurons that project their axons contralaterally across the midline. Although CRISPR mutagenesis of *Ddpep* did not result in any noticeable axon guidance defects, we speculate that activation of *Ddpep* by Pax3/7 in both neuron types might act in conjunction with partially redundant/overlapping pathways to ensure robust midline crossing. If true, this would mean that different nodes of the ddN regulatory network may carry out specific morphogenetic behaviors and can be shared with other neurons arising from distinct cell lineages.

Similarly, we found that Pou4 regulates *Saxo* in the ddNs and in the neurons of the peripheral nervous system. Saxo proteins are known to localize to cilia in neural cells in *Ciona* and vertebrates (Dacheux et al., 2015; Erickson et al., 2023; Kim et al., 2020). The ciliated PNs of the papillae mechanically sense the attachment of swimming *Ciona* larvae to suitable substrates and this is sufficient and necessary to trigger metamorphosis into the sessile adult phase (Sakamoto et al., 2022; Wakai et al., 2021). However, targeting this gene specifically in the sensory papillae did not result in a significant metamorphosis defect. Although we found that the ancient regulatory connection between *Pou4* and *VGLUT* is evolutionarily conserved in *Ciona,* it is not known if Pou4 factors regulate *Saxo* orthologs in vertebrate ciliated and/or sensory cells.

In contrast to the PNs and other epidermal neurons, the ddNs and the BTNs are not ciliated. We hypothesized that Saxo, like Defcab, might regulate polarized axon outgrowth due to its proposed role as a microtubule stabilizer and its localization to the centrosomes. However, we did not observe any significant polarity or axon growth defects in ddNs or BTNs, casting doubt on this hypothesis. It is possible that Saxo functions redundantly with Nckap5, a similarly microtubule-stabilizing, centrosome-enriched protein that we show here is required for axon outgrowth in the ddNs. It is also possible that Saxo carries out another unknown function in ddNs/BTNs for which we did not assay. Alternatively, its expression might not be important for ddN or BTN development, and is instead but a consequence of its direct activation by Pou4 and its adaptive value may come from its functions in another Pou4+ neuron type. One additional possibility is that our CRISPR/Cas9 approach to targeting genes like *Saxo,* encoding proteins made of numerous domain repeats (12 Mn domains in Saxo), might not be effective at completely knocking out their function, as the resulting truncated polypeptides might still retain enough domains to remain partially functional.

Finally, although we identified *Seldom* as another gene that is activated by Pou4 in both ddNs and PNs, the function of Seldom protein is unknown. Although we could not find Seldom orthologs in mouse or humans, we have identified putative orthologs in diverse invertebrates and chordates, including some mammals (see **Supplemental Sequences File**). Future work will be required to fully characterize the function and evolutionary history of this potentially conserved gene, and the relative contributions of other effectors to ddN morphogenesis.

## Supporting information

Supplemental Sequences File

## Acknowledgments

We thank the lab for insightful discussion and comments over the course of this work. We thank Shohon Rafique for identifying candidate sgRNAs, and Serina Suzuki, Mana Morita, Susanne Gibboney, and Lindsey Cohen for technical assistance. We thank Christina Cota, Liang Han, Farzaneh Najafi, and Shuyi Nie for comments and suggestions, and Florian Razy-Krajka and Wei Wang for scientific ideas and guidance. This study was funded by NSF IOS grant 1940743, NIH grants R01HD104825 and K99HD084814, and funds from the Nakatani Foundation RIES program.

## Notes

### Competing Interest Statement

The authors have declared no competing interest.

