## Supplemental Figures for "A gene regulatory network for specification and morphogenesis of a Mauthner Cell homolog in non-vertebrate chordates"

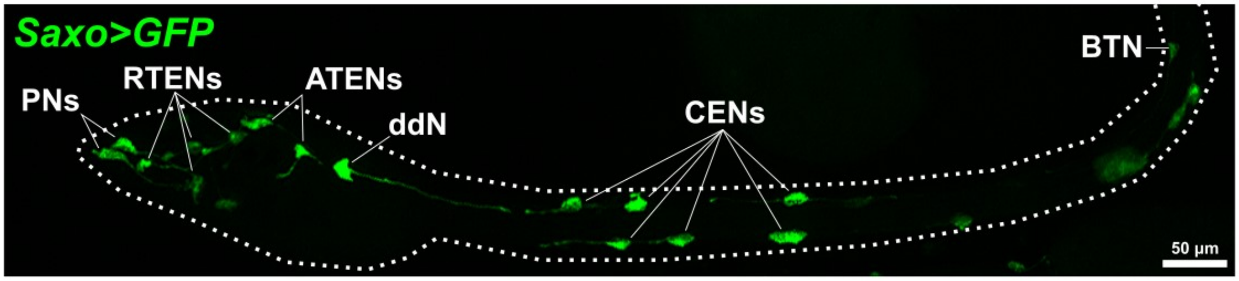


Supplemental Figure 1.

Larva at 19 hpf/stage 28 electroporated with the *Saxo* reporter, showing expression in ddNs, bipolar tail neurons (BTN), caudal epidermal neurons (CENs), apical trunk epidermal neurons (ATENs), rostral trunk epidermal neurons (RTENs), and papilla neurons (PNs). Larval body outline indicated by dashed line.

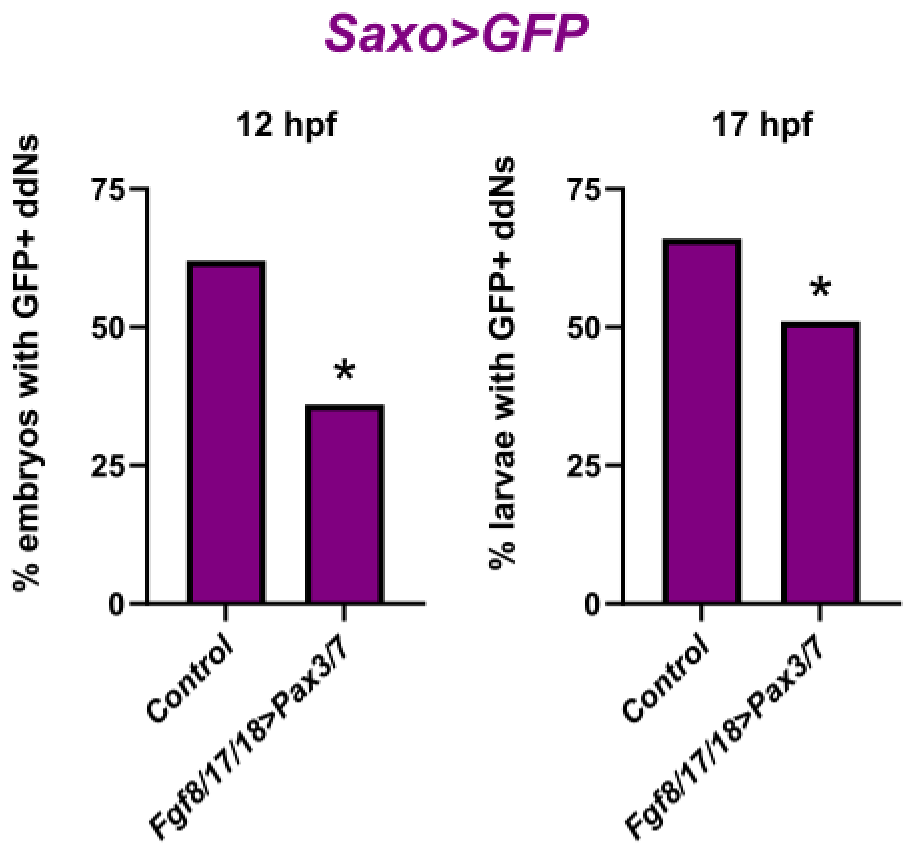


Supplemental Figure 2.

Plots showing reduced frequency of *Saxo* reporter expression in individuals electroporated with *Fgf8/17/18>Pax3/7* relative to negative control animals, assayed at two different developmental stages, 17 hpf/stage 27 larvae and 12 hpf/stage 24 mid-tailbud embryos. See text for experimental details and **Supplemental Table 2** for statistical test information. * $p < 0.01$. All animals raised at 20°C, $n = 50$ for all conditions except *Fgf8/17/18>Pax3/7* at 17 hpf ($n = 60$).

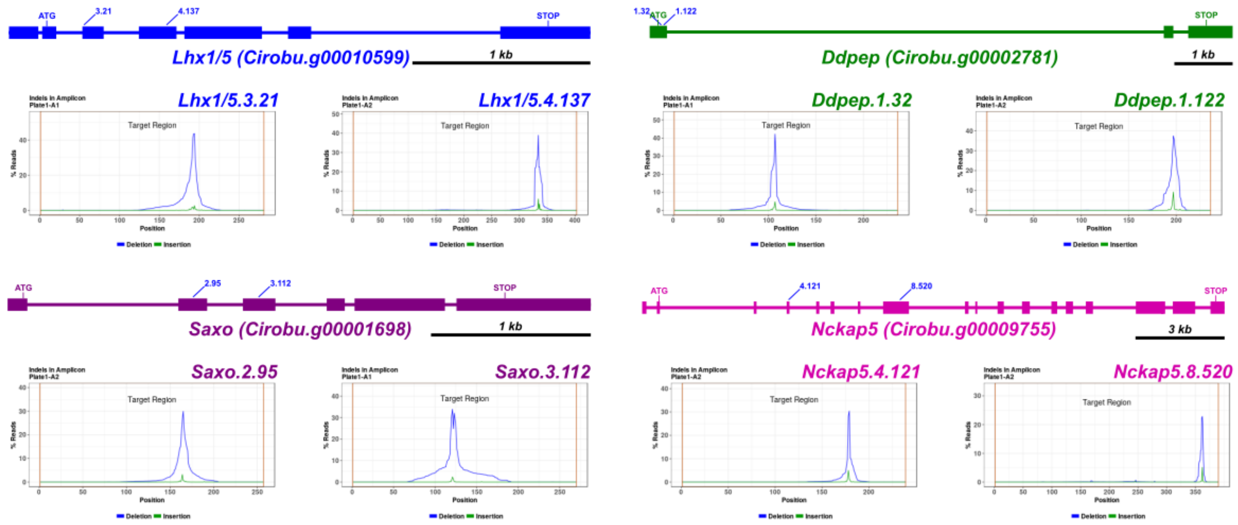


Supplemental Figure 3.

Schematic diagrams indicating target sites of sgRNAs designed and validated to target *Lhx1/5, Ddpep, Saxo,* and *Nckap5* for CRISPR/Cas9-mediated mutagenesis in this study. Plots show indel coverage and frequency for each validated sgRNA, as determined by E-Z amplicon Illumina sequencing (GENEWIZ by Azenta). See text for validation protocol and details.

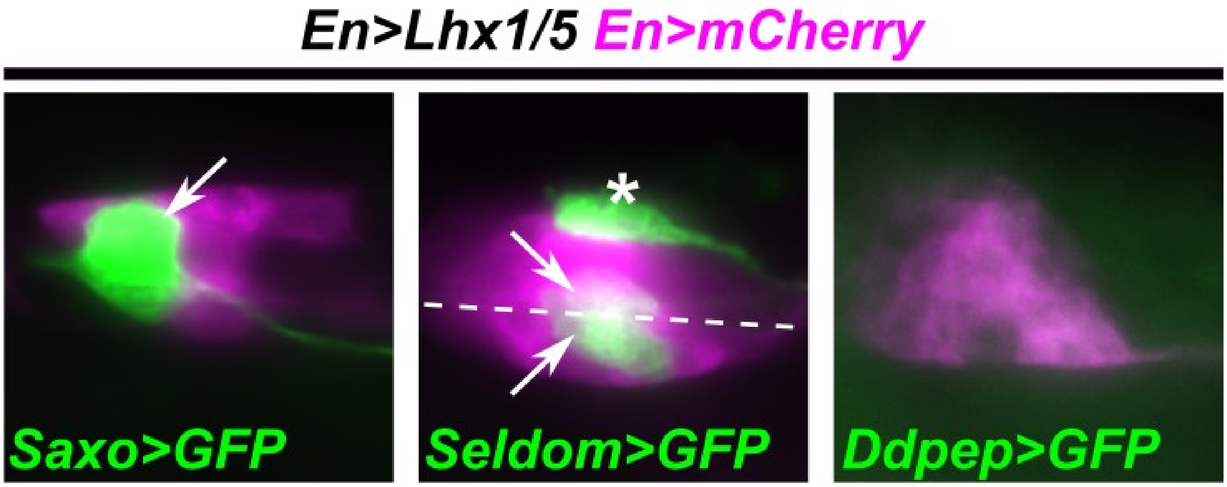


Supplemental Figure 4.

Lack of ectopic activation of *Saxo, Seldom,* and *Ddpep* reporters upon Lhx1/5 overexpression in MG. Arrows indicate a single ddN or ddN left/right pairs. Dashed line indicates midline. Asterisk indicates AMG neuron labeled by the Seldom reporter. Note reduced expression of *Ddpep* reporter upon overexpression of Lhx1/5. Larvae raised at 20°C to 19.5 hpf/stage 29.

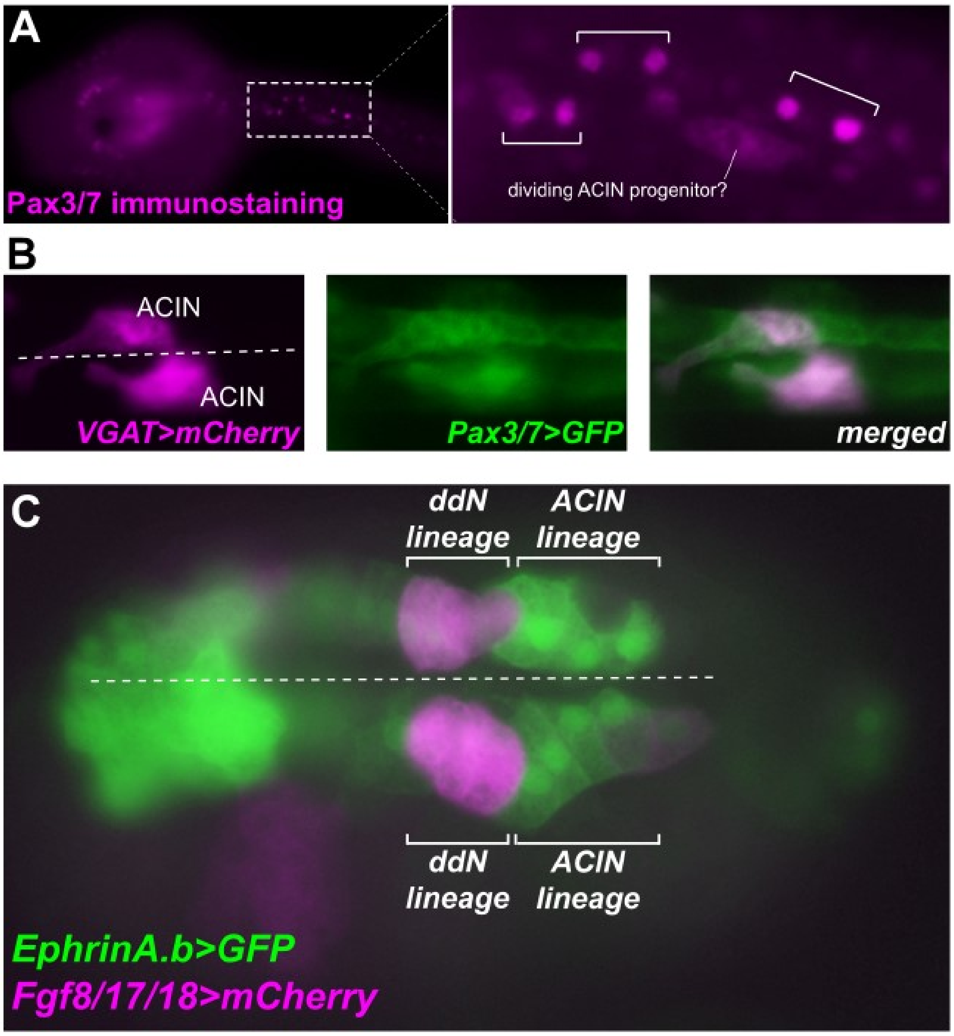


Supplemental Figure 5.

A) Immunostaining for Pax3/7 protein labels certain nuclei in the anterior part of the tail at 12 hpf 20°C (~stage 24). These nuclei are consistent with belonging to that of the ACIN lineage based on position and numbers, following the description of Nishitsuji et al. 2012. Brackets in magnified inset denote each ACIN-ependymal precursor sibling pairs, while one progenitor cell appears to be completing cell division. B) Expression of *Pax3/7* reporter in the ACIN lineage suggested by co-expression of *Pax3/7* and *VGAT* reporters in hatched larvae (~stage 27). Both left and right embryo halves shown, along with the embryonic midline (dashed line). C) An EphrinA.b reporter labels the lateral rows of the neural tube including the A9.29 lineage that gives rise to the ACINs (~stage 18). The ddNs arise from the A9.30 lineage labeled by the *Fgf8/17/18* reporter primarily. Both left and right halves shown, along with the midline (dashed line).

```
CLUSTAL format alignment by MAFFT (v7.490)

Cirobu.Ddpep    MTSQRSTTRTSVVVSVLVLLFW------------------------------CQVFYTSK
Stclva.Ddpep    MTQKVFKFN--AYWIIAGFLFGILLHVGPSKE--AKLEKSAIKQVLKNYEKVVDAQSSLT
Harore.Ddpep    MWQTRLGSNL-VYVALLCFLLCKPVMVASKRQVNAKPQKGDLQNGIQQFHTDSQATIIAD
                * .     .  .   :  :*:                                :.    

Cirobu.Ddpep    ADGATVEL--NREAKVRSKRHVADMLITEKVSRVKRGHNKNLLLSRILQLTDASTGGHCQ
Stclva.Ddpep    DASSEVLF--------RGKRHTADHLMNDKYSRTLKSHARKQILSTLLKLVGSTAGGPCQ
Harore.Ddpep    EAGKTTEFSLGESGRSRQKRHTADHLISDKYSRSLKSQARKQILSKLLNLVGSSPGNPCE
                  .  . :        * ***.** *:.:* **  :.: :: :** :*:*..::.*. *:

Cirobu.Ddpep    CILQQKA---------------------VKQGCECNEPILYLRMIRKITDPKM---KLGH
Stclva.Ddpep    CESHSHLDDQTSRLKSRQDAISKRVH--TEPSFDCNCDMYSL--VSYLAERLSQGGEDEE
Harore.Ddpep    CLFAQSTSDKDSSSRLIQRDVFKRLDSISSSSFKCNCDISSL--IKFILSKL----KFTK
                *   .                        . . .**  :  *  :  : .      :  .
                          Peptide from Kawada et al. 2011
Cirobu.Ddpep    GSDATELDRLLARMKPNMKRAVLHLAINEFQRLLREVEVERTRRGRIIRQLIRRRRKLRH
Stclva.Ddpep    STGSDPLSGYLSRLSPSVKARLVHFLNEESMRIKRNEMLRKLRRMRLHRQFLHRRRLLRN
Harore.Ddpep    NSNTDPLGRYMSMLSEATKTRLAHFLHEEELRLKKNEILRKLRIRRMHRMMMHRRRLLHK
                .:.:  *.  :: :.   *  : *:  :*  *: ::  :.: *  *: * :::*** *::

Cirobu.Ddpep    HPRSAFDGKQLFSF
Stclva.Ddpep    DPNAAFRGKQFLSF
Harore.Ddpep    DPRAAFRGKQFFSF
                .*.:** ***::**
```

Supplemental Figure 6.

Alignment of Ddpep ortholog protein sequences from different tunicate species. Cirobu = *Ciona robusta,* Stclva = *Styela clava,* Harore = *Halocynthia roretzi.* Peptide detected by mass spectrometry in Kawada et al. 2011 indicated in red lettering, although this portion is not conserved in other tunicates. See supplemental sequences file for protein sequences in FASTA format.

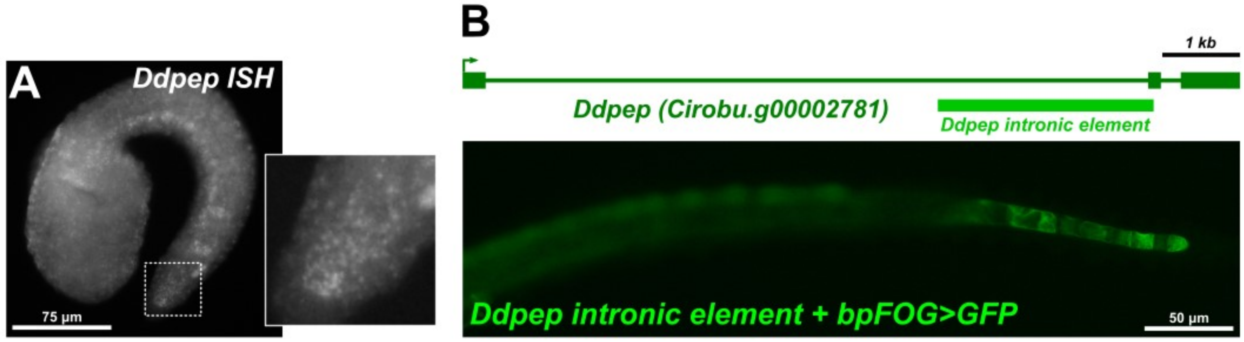


Supplemental Figure 7.

A) Whole-mount fluorescent *in situ* mRNA hybridization showing *Ddpep* expression in the posterior tip of the notochord (magnified inset) at 9.5 hpf/stage 21. Expression can also be seen in ddNs and endodermal strand. B) Top: schematic diagram of intronic *Ddpep cis*-regulatory fragment that drives notochord tip expression image underneath. Bottom: larva at 19 hpf/stage 28 electroporated with the *Ddpep* intronic fragment reporter showing expression in notochord tip cells.

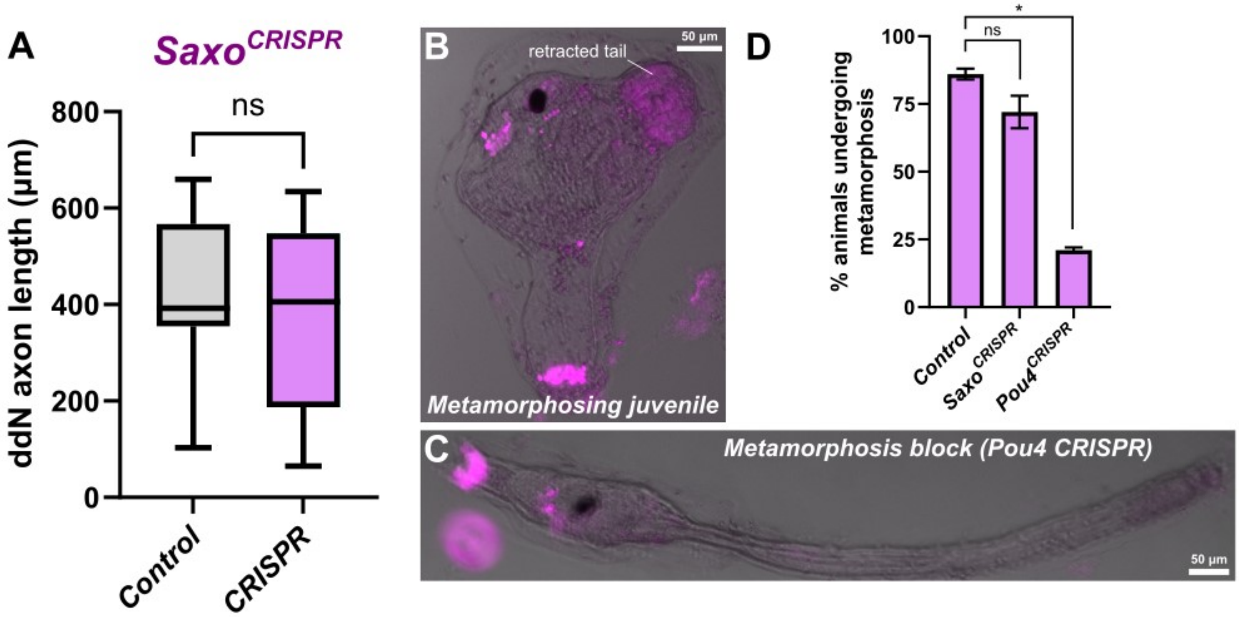


Supplemental Figure 8.

MG-specific CRISPR/Cas9-mediated mutagenesis of *Saxo* does not cause significant reduction in average ddN axon length. Larvae fixed and imaged at 19.5 hpf, 20°C (stage 29). For control CRISPR, n = 23, for *Saxo* CRISPR, = 20. Non-significance (ns, p = 0.4616) was determined by two-tailed Mann Whitney test. E) Example of a normal, metamorphosing juvenile (negative control condition) at 50.5 hpf. Magenta color indicates expression of *Foxc>H2B::mCherry.* F) Example of a larva at 50.5 hpf in which metamorphosis has been blocked, in this case by papilla-specific disruption of *Pou4* by CRISPR/Cas9. Magenta indicates *Foxc>H2B::mCherry* expression. G) Plot showing quantification of metamorphosis defects in different CRISPR conditions, by assaying tail retraction (or no tail retraction) in H2B::mCherry+ individuals as in panel E and F. Experiments were performed and scored in duplicate, with n = 50 in each condition and each duplicate. See Supplemental Table 2 for statistical test details. * p < 0.0001, ns = not significant.
