## Supplemental Sequences File for "A gene regulatory network for specification and morphogenesis of a Mauthner Cell homolog in non-vertebrate chordates"

**Supplemental Sequences File, Kim et al.**

**Partial plasmid sequences**

>Fgf8/17/18 -4835/+12 “sec” from Imai et al. 2009

mutation to disrupt endogenous Fgf8/17/18 start codon

gcatagatttaagccacctcgtgaagtgggcgggtttaattaaaacaaacagagctccagcaatcagtgcaatcaagacgtttgtcgtaggttaaaatattgctgctaaagccgcttctgatcagataaattatgataaaatcaaaccgcatttgataacatctgctccgtattgttttgccggtaagtatggacatagacattaaaaaatgtggaataaatatgaaaaacgcggtataatgatggactataggtatatgcctggtttgaaccgaaaatgcaaaggtcacagatctaaagctgttgaacgcaaggtgttcaagcgaaaatttctgtgaaactgctgtttggtatttttccgatgctaattcggtgcgttgagagaaccccatggaatgctgtaatatgctagcgtttgttgttacaataaacatacctaagtttaagcgttataaaaatctagcaatacaccaaacgtacatgtacagaaaagtcgtggtttaccaataataaataagttatgtgtatatatatagcaccggtattctattcttatctcgcaacctcagtttactggactgaaaacaaatcgggcttgtctccctttgttctgagtcttgtccacaacaaagcgtcagatcgcgctggtttaatgatctcgaatttcagatcatcagttttcagtttcccaaatattccaactatccagtaccgctagcacagggtacacaaataggctgtgtatcggctctagcgacttgacctcgtcggtgtttgtgcaaatttataatcccgccagcctaaacagggcaagggccgctgaggccgcattgacatacaagtacgcgatgcttgcaaaatgccagtccgcaatttctcgcaaaaaaacgtgactgcactctcggggtagcgcattgttccggcggctcctgcctgcacggcgttttcatgaatcacgcggctgtctaatctacgatcaacggccttagccaaaaacccggatcggtaaaagttaacactgttgcgaactcaatacctcgaaattcttgaggttactgcggttactcgggccggattatccgtcaaaaataagcgaagtcgggttcagtttacgtttctcctaactattcccaactgaaaatgacaattaaaaaaaggatcatgatattgtgattagcatgtcatccagcaacgagctgtgtatatgtaggagcatgttttccgaaaggattactattaattttgaagtttgatttcagaaaatcgattatgagcaaacaagtaatgtgcggaaacgattataatacctttttacgaagcagtgtaccagttaaaacagtaaatcaaattctgttactaaagcagtcgatgtttaaacggctgggtgtatctcgctttcaagagtcggcgtcggtgtccggaaaccccgcttctcccagaggtagtgttgtgaaaaatgccggcccagctattctcattgtttagcctccgataatgggaaatacaacgcgtgtctgagataccgctttcccattcctcaagtacgttgttcgcctgcctaccgtccattgactgctggttaattacatcaacttgttttatgggaaacgcttatcgcacacagagagcgtgacgcacagaatcagagagcacatgtaatataaagttgtatgtaaacaagggagttttagtccttagtgagcgccatacgattgcgatcggatgtttatcagcctcggctagtgcacagtttaaacgttctgatgttagttgccgagccctctcagcaagtatgtatacagctgtttattcgggcctctgtagttctagtgttaaggaaggcagactcgactgcacgctacaagcgacagcgctttttccatcaatccaatcacatttgttgagccccttacgggcacttgaggtgacggcgctaatttctaagcagtttcgattgtgtgggcagcttggccgcagagcaaagcacagaaaacgcggcgcttcgcattgcagcagcgccgaacactttgccaaggttgcagtatatgaatacagtagcgccgcgaaagacggagagtttttaaagcacaataagcgtgcctgcggtctgtacatacatacagcaaaattttcggcgttgtgtaaagttggtcgcgacaacaatagccctctcggtgtttggtgcttgaagcggcactacaacagcccgctatcataggccgggaattagggcaagtggcggcttagcgagaatttaattttccttgttaagcaaaatcgtgtgaagtgcttctgtcccacgactgttcacttcctgcgacaacgtatagtgagatcgtacttttgtagccaagataccaatatcaataataatattcagctttgcccgctaggttgaggtaaagggtagatattagttgagaatagttgcgtaatatgattaagctcgtgtaccggcttgctttatatatttttgctaaacaaacaagtgtagcgattaaaatcactcaggcctaagccaagggtaaaaccggttaattcggaggttttagaaacagttattttttgcggtttcactaaacttagaataaatacccaagaatattattatggtttataatttttttttaagttttttttttaaatatccattttttggtgattgtgattgatttgtaataagatatttttaacttcgatattaatgtttgcttaaaattacgaaatagagccgtctgctgcgaaggtttgtaaagtgaaacaattagtgggctactgcagttctactttctgtacaaatatacgctttatcacgctgtcaccttgacagacgttattgtaggaaagtttttgtgctgaacaatagccttgtaattagttgcgggggggcctctcggttcaaggccgaatagatacgacctctgatcagcgggtcgaaaacgaacgtaacaatagggaattatttcgtgctgtgacgtcggccgagtggggctgggaaccatttctaaaacattcctggcataatgatacgatgtaagcgcaatatttcacaaacgaaacaagtttacgtgactgtaactgcataagcgttgctttgtcactggaggcgccaaccgcacgataagatatttcagcgacggtcgcaatgaaaacggacaccctattacatgtatgacggccgttaatacctctgtcgctttgacaccttccaattgtcgataacataacccgcaacttgatattctgtgaatattatgacacccggcgccttttgcctgaattgtgtaatcgtgccgcaatattcgaaattgtcctgggtttatttttaaggcagacgtcagagaaattataattctttacatcagatctgataaaccgctcgatttcacggatgtgttgaacaagcgccgtttcaaactgtcgcagttttatttttatcttcgtataaatttacagcggaattcattgaatgctaaactgttgtataaagaagagtgcggaaagacgggccttgttgggacatactatcaaactattctgatcgtgttttaaacaattaacaacggtatatggtattcgcgagaaaacggtcttataatttttgaatattctttgtttactaccgaataggacaagaaaatagaatgaataggtgtcccatctttccccaccctatactatatatgaaccacacgtactccaaacacacttaacagcgtgacatatttcgctattgcgccagtttaaaacacttctgtcttgagttcaataggcatacgtgtcatagttactacacacgttgtaacttttcccgtatcggaagtttttaaaaaaaaataggactctcctatgacgaattatattaattgctcccctggcgtggtttattttgtcttttcccacttattttcctttgcccacgttttatacagttacctttaatttgctacgcgtttgctcgacgggcggcatgagcgttttggggattatgacgtaacaaaacggtttttaatctcttcctctatgttacgacataatagcacctgtggaaaaacacaaacctgttttgttgggaagctgtatggttgtaaagggatgtgtgtttcgttttgttggtctctgctaaatacacgagtgcaatttcagtcaattacgaataccaacgtatgaaaagaatgcgcagattttgctaattggtttgtttaccctcgttcgtgcttggaattctgcaaattgttgttgtgtgggttagaacaatccaatcaccctgaaagttttttttaaatgcagaagaaaaccattaattttgtgcccaaaacaccgtaagctgtagcgagcacaagttcgatatgtgttcatcaaaaatcggcggactttttctcatcgtcgcttttttcccgcggctccagcaacgcgtttgttaaaagcgcgttgcctaccgttatttacattaaacgatgccgatcgctgggcggttcgttatttttactcgtctggtaatggacaaaataacaacccagtcgtgtgcttagattttagcggtatggagcgccacgcgccgaccctcgggaaggacatgctgcacagcgggaaaccgagctttaggatccggcagataattttattcggagtcgatgttagaattgttataagttgttcaggataatagcaaagtgaaagcagaaataaatttaaactttatgctttgtatatttattcagtaagtagttgaataatgtcaattcggatttaatgcattgcgagtataaatagtaaatccgaatatcaaagtgatttctaagggacattattcttctcttggattacatacgaaaatAcgaccctcc

>Fgf8/17/18 -4835/+12 from Imai et al. 2009

(original version with endogenous start codon intact)

gcatagatttaagccacctcgtgaagtgggcgggtttaattaaaacaaacagagctccagcaatcagtgcaatcaagacgtttgtcgtaggttaaaatattgctgctaaagccgcttctgatcagataaattatgataaaatcaaaccgcatttgataacatctgctccgtattgttttgccggtaagtatggacatagacattaaaaaatgtggaataaatatgaaaaacgcggtataatgatggactataggtatatgcctggtttgaaccgaaaatgcaaaggtcacagatctaaagctgttgaacgcaaggtgttcaagcgaaaatttctgtgaaactgctgtttggtatttttccgatgctaattcggtgcgttgagagaaccccatggaatgctgtaatatgctagcgtttgttgttacaataaacatacctaagtttaagcgttataaaaatctagcaatacaccaaacgtacatgtacagaaaagtcgtggtttaccaataataaataagttatgtgtatatatatagcaccggtattctattcttatctcgcaacctcagtttactggactgaaaacaaatcgggcttgtctccctttgttctgagtcttgtccacaacaaagcgtcagatcgcgctggtttaatgatctcgaatttcagatcatcagttttcagtttcccaaatattccaactatccagtaccgctagcacagggtacacaaataggctgtgtatcggctctagcgacttgacctcgtcggtgtttgtgcaaatttataatcccgccagcctaaacagggcaagggccgctgaggccgcattgacatacaagtacgcgatgcttgcaaaatgccagtccgcaatttctcgcaaaaaaacgtgactgcactctcggggtagcgcattgttccggcggctcctgcctgcacggcgttttcatgaatcacgcggctgtctaatctacgatcaacggccttagccaaaaacccggatcggtaaaagttaacactgttgcgaactcaatacctcgaaattcttgaggttactgcggttactcgggccggattatccgtcaaaaataagcgaagtcgggttcagtttacgtttctcctaactattcccaactgaaaatgacaattaaaaaaaggatcatgatattgtgattagcatgtcatccagcaacgagctgtgtatatgtaggagcatgttttccgaaaggattactattaattttgaagtttgatttcagaaaatcgattatgagcaaacaagtaatgtgcggaaacgattataatacctttttacgaagcagtgtaccagttaaaacagtaaatcaaattctgttactaaagcagtcgatgtttaaacggctgggtgtatctcgctttcaagagtcggcgtcggtgtccggaaaccccgcttctcccagaggtagtgttgtgaaaaatgccggcccagctattctcattgtttagcctccgataatgggaaatacaacgcgtgtctgagataccgctttcccattcctcaagtacgttgttcgcctgcctaccgtccattgactgctggttaattacatcaacttgttttatgggaaacgcttatcgcacacagagagcgtgacgcacagaatcagagagcacatgtaatataaagttgtatgtaaacaagggagttttagtccttagtgagcgccatacgattgcgatcggatgtttatcagcctcggctagtgcacagtttaaacgttctgatgttagttgccgagccctctcagcaagtatgtatacagctgtttattcgggcctctgtagttctagtgttaaggaaggcagactcgactgcacgctacaagcgacagcgctttttccatcaatccaatcacatttgttgagccccttacgggcacttgaggtgacggcgctaatttctaagcagtttcgattgtgtgggcagcttggccgcagagcaaagcacagaaaacgcggcgcttcgcattgcagcagcgccgaacactttgccaaggttgcagtatatgaatacagtagcgccgcgaaagacggagagtttttaaagcacaataagcgtgcctgcggtctgtacatacatacagcaaaattttcggcgttgtgtaaagttggtcgcgacaacaatagccctctcggtgtttggtgcttgaagcggcactacaacagcccgctatcataggccgggaattagggcaagtggcggcttagcgagaatttaattttccttgttaagcaaaatcgtgtgaagtgcttctgtcccacgactgttcacttcctgcgacaacgtatagtgagatcgtacttttgtagccaagataccaatatcaataataatattcagctttgcccgctaggttgaggtaaagggtagatattagttgagaatagttgcgtaatatgattaagctcgtgtaccggcttgctttatatatttttgctaaacaaacaagtgtagcgattaaaatcactcaggcctaagccaagggtaaaaccggttaattcggaggttttagaaacagttattttttgcggtttcactaaacttagaataaatacccaagaatattattatggtttataatttttttttaagttttttttttaaatatccattttttggtgattgtgattgatttgtaataagatatttttaacttcgatattaatgtttgcttaaaattacgaaatagagccgtctgctgcgaaggtttgtaaagtgaaacaattagtgggctactgcagttctactttctgtacaaatatacgctttatcacgctgtcaccttgacagacgttattgtaggaaagtttttgtgctgaacaatagccttgtaattagttgcgggggggcctctcggttcaaggccgaatagatacgacctctgatcagcgggtcgaaaacgaacgtaacaatagggaattatttcgtgctgtgacgtcggccgagtggggctgggaaccatttctaaaacattcctggcataatgatacgatgtaagcgcaatatttcacaaacgaaacaagtttacgtgactgtaactgcataagcgttgctttgtcactggaggcgccaaccgcacgataagatatttcagcgacggtcgcaatgaaaacggacaccctattacatgtatgacggccgttaatacctctgtcgctttgacaccttccaattgtcgataacataacccgcaacttgatattctgtgaatattatgacacccggcgccttttgcctgaattgtgtaatcgtgccgcaatattcgaaattgtcctgggtttatttttaaggcagacgtcagagaaattataattctttacatcagatctgataaaccgctcgatttcacggatgtgttgaacaagcgccgtttcaaactgtcgcagttttatttttatcttcgtataaatttacagcggaattcattgaatgctaaactgttgtataaagaagagtgcggaaagacgggccttgttgggacatactatcaaactattctgatcgtgttttaaacaattaacaacggtatatggtattcgcgagaaaacggtcttataatttttgaatattctttgtttactaccgaataggacaagaaaatagaatgaataggtgtcccatctttccccaccctatactatatatgaaccacacgtactccaaacacacttaacagcgtgacatatttcgctattgcgccagtttaaaacacttctgtcttgagttcaataggcatacgtgtcatagttactacacacgttgtaacttttcccgtatcggaagtttttaaaaaaaaataggactctcctatgacgaattatattaattgctcccctggcgtggtttattttgtcttttcccacttattttcctttgcccacgttttatacagttacctttaatttgctacgcgtttgctcgacgggcggcatgagcgttttggggattatgacgtaacaaaacggtttttaatctcttcctctatgttacgacataatagcacctgtggaaaaacacaaacctgttttgttgggaagctgtatggttgtaaagggatgtgtgtttcgttttgttggtctctgctaaatacacgagtgcaatttcagtcaattacgaataccaacgtatgaaaagaatgcgcagattttgctaattggtttgtttaccctcgttcgtgcttggaattctgcaaattgttgttgtgtgggttagaacaatccaatcaccctgaaagttttttttaaatgcagaagaaaaccattaattttgtgcccaaaacaccgtaagctgtagcgagcacaagttcgatatgtgttcatcaaaaatcggcggactttttctcatcgtcgcttttttcccgcggctccagcaacgcgtttgttaaaagcgcgttgcctaccgttatttacattaaacgatgccgatcgctgggcggttcgttatttttactcgtctggtaatggacaaaataacaacccagtcgtgtgcttagattttagcggtatggagcgccacgcgccgaccctcgggaaggacatgctgcacagcgggaaaccgagctttaggatccggcagataattttattcggagtcgatgttagaattgttataagttgttcaggataatagcaaagtgaaagcagaaataaatttaaactttatgctttgtatatttattcagtaagtagttgaataatgtcaattcggatttaatgcattgcgagtataaatagtaaatccgaatatcaaagtgatttctaagggacattattcttctcttggattacatacgaaaatgcgaccctcc

>Fgf8/17/18 -2401/+3 START

atgattaagctcgtgtaccggcttgctttatatatttttgctaaacaaacaagtgtagcgattaaaatcactcaggcctaagccaagggtaaaaccggttaattcggaggttttagaaacagttattttttgcggtttcactaaacttagaataaatacccaagaatattattatggtttataatttttttttaagttttttttttaaatatccattttttggtgattgtgattgatttgtaataagatatttttaacttcgatattaatgtttgcttaaaattacgaaatagagccgtctgctgcgaaggtttgtaaagtgaaacaattagtgggctactgcagttctactttctgtacaaatatacgctttatcacgctgtcaccttgacagacgttattgtaggaaagtttttgtgctgaacaatagccttgtaattagttgcgggggggcctctcggttcaaggccgaatagatacgacctctgatcagcgggtcgaaaacgaacgtaacaatagggaattatttcgtgctgtgacgtcggccgagtggggctgggaaccatttctaaaacattcctggcataatgatacgatgtaagcgcaatatttcacaaacgaaacaagtttacgtgactgtaactgcataagcgttgctttgtcactggaggcgccaaccgcacgataagatatttcagcgacggtcgcaatgaaaacggacaccctattacatgtatgacggccgttaatacctctgtcgctttgacaccttccaattgtcgataacataacccgcaacttgatattctgtgaatattatgacacccggcgccttttgcctgaattgtgtaatcgtgccgcaatattcgaaattgtcctgggtttatttttaaggcagacgtcagagaaattataattctttacatcagatctgataaaccgctcgatttcacggatgtgttgaacaagcgccgtttcaaactgtcgcagttttatttttatcttcgtataaatttacagcggaattcattgaatgctaaactgttgtataaagaagagtgcggaaagacgggccttgttgggacatactatcaaactattctgatcgtgttttaaacaattaacaacggtatatggtattcgcgagaaaacggtcttataatttttgaatattctttgtttactaccgaataggacaagaaaatagaatgaataggtgtcccatctttccccaccctatactatatatgaaccacacgtactccaaacacacttaacagcgtgacatatttcgctattgcgccagtttaaaacacttctgtcttgagttcaataggcatacgtgtcatagttactacacacgttgtaacttttcccgtatcggaagtttttaaaaaaaaataggactctcctatgacgaattatattaattgctcccctggcgtggtttattttgtcttttcccacttattttcctttgcccacgttttatacagttacctttaatttgctacgcgtttgctcgacgggcggcatgagcgttttggggattatgacgtaacaaaacggtttttaatctcttcctctatgttacgacataatagcacctgtggaaaaacacaaacctgttttgttgggaagctgtatggttgtaaagggatgtgtgtttcgttttgttggtctctgctaaatacacgagtgcaatttcagtcaattacgaataccaacgtatgaaaagaatgcgcagattttgctaattggtttgtttaccctcgttcgtgcttggaattctgcaaattgttgttgtgtgggttagaacaatccaatcaccctgaaagttttttttaaatgcagaagaaaaccattaattttgtgcccaaaacaccgtaagctgtagcgagcacaagttcgatatgtgttcatcaaaaatcggcggactttttctcatcgtcgcttttttcccgcggctccagcaacgcgtttgttaaaagcgcgttgcctaccgttatttacattaaacgatgccgatcgctgggcggttcgttatttttactcgtctggtaatggacaaaataacaacccagtcgtgtgcttagattttagcggtatggagcgccacgcgccgaccctcgggaaggacatgctgcacagcgggaaaccgagctttaggatccggcagataattttattcggagtcgatgttagaattgttataagttgttcaggataatagcaaagtgaaagcagaaataaatttaaactttatgctttgtatatttattcagtaagtagttgaataatgtcaattcggatttaatgcattgcgagtataaatagtaaatccgaatatcaaagtgatttctaagggacattattcttctcttggattacatacgaaaATG

>HA::Pax3/7 coding sequence from Stolfi et al. 2011

ATGGCATACCCATACGACGTGCCAGACTACGCAGAACTAGCGCATCCAGGGTCTAATTTCCGACCAGCTTTTCCGTTGGAAGGTCAAGGGCGAGTAAACCAACTCGGTGGCATGTTCATAAACGGTCGCCCTCTTCCCAGTCATGTACGACATAAGATAGTGGAGATGGCAGCTCATGGTGTGCGACCGTGTGTCATAAGTCGACAGCTAAGGGTCTCACATGGATGTGTCAGTAAAATTTTATGCAGATACCAAGAAACAGGATCAATTAAACCGGGAGCAATCGGTGGCAGTAAACCAAAAGTAGCCACCGCAGATGTCGATAATAAAATAGAAGAATACAAAAAAGAAAATCCGGGAATCTTTAGCTGGGAAATCAGAGAACGACTTATCAAGGAAGGTATATGTGATCGAAGTAACGTTCCTTCGGTCAGTTCTATTAGTCGAACTCTGCGTGCAAAAGGTTGTGACGTCGAAAACGAATCAGAATCCAGCGCAAGACTCGATCCAGGTAACCGCTCTTCTTCTTCCGGAGGAGAGCCGAACGAAGTCGGAGGATCCGACAGTGAGTCAGAGCCAGATCTTCCTCTTAAAAGGAAGCAAAGAAGAAGTAGGACGACTTTTAGCGCGGAGCAGCTGGACGAGCTAGAGAGATGCTTTGAAAGAACTCATTATCCTGATATTTATACAAGGGAAGAACTTGCTCAAAGAACAAGACTAACTGAGGCTCGAGTTCAGGTGTGGTTCAGCAACAGACGGGCAAGATGGCGGAAGCAAATGGCGGCTCAACAATTCCATGGCATTCCAGCCCATCACCCCCACCACCTATCACCCCACCTCGGGTACCCACACACAATGCCCGGGTCAGTGGGCCAAGCTGCACATAACTATATGTTGCAGACAAGTGGGGCACATCATGTACAGAGCATGCATGAAAGCTTTGCACATACGGCGTCTTCTCATGATCACGGTTCCTCTCTCCATCGCCCCCATGCGTCTGTACTTACGTCACCAGTTCACCACGCGTCACAAGCCATTAGATACGACACCACAGATCCACTTAATCAAAGTGCAGCCGCTGCTTATGGTTTAATGGGAGCATCAGCTGCAGCAGCGGCCGCAGCAGCTCGCTATAGTCAGTATCCATCGGCCGCTGCTTTCGGCGACGCTTACCCGATACATTATCACCACCACTCAACGTCCCCACCTAGCGGTAGCTCGATTTTCGGTTTTCCGTCACACACCCGACAAACTGCAACAAACCCAGTCGAGCTCACCACCGGGAACCCCCATTCCGGCCCCAACTCTTCACCCATCGATGTGGGTTCACCCGGAGAAGCGGGGAATCACCCCCTGCACCAGCAACACTTTGTTTTAGCACAGAGTGCACGAAGAGCCCACCAAGCAAGCAATCAACGCAATCACCACAACCAACACCACAGCGGGGAAACTACCACCGCACCTTCAACGTCACCAGCATCAAATGGAGATGGACACCCAACAACTGGAATTGAATCGCAATCGCGAGCATATTCAGTACTAGCAGGTCAACAACACCACAACACTGACTTGCATAGCTCAACCTCCGTAGCGGGCCAGCAATCGACAGCTATGCTCTCTTCGAGCGTCGACCACTCAAGCAATGTATTCGCTGGACAGGAGAGGGGTGCGCCGTCATCTCGAGCCGGATACGGTGTTGGCCCATCAGGATTTGCAACTGGTGAAGTACTACCAACATCAAGTGCTGCTTCACATCAATCTTACGCATCATGTCAATACTCACCGTACGGACAAGCCAGTGGTGATTATGGAAGTGCCGGAATAGCTGCTCTACGCATGAAATCCCGGGAACACAGCGCTTCAATTGGACTTATCCCTGTCGGCGGAGGACCGAGCGTTCAGCATGCATATTGA

>MRF -904/-1 from Christiaen et al. 2009

ggcttacgcatctcgagcgaaccagtacgagtgaaaattataacttcaaagtctgcccggcacgtgacttagagtctcccgcgagaccggcgctccggagttctggaatgcagaagaagaagaagcattgtcacctactgtgacgtcataaacgagtgtcttaaatagcgcctgcatcaagccatgcattctataacatcgctagccagccacgcatggtaaagttatcaagcgatagccagtcaatgatatatagtagccaatgtctgttataaccaggcatggtcacatagcgtgttttaatacaagtagatctacatggtatatagttagcggcgaactaaagttttaaactgcagcaaattgcagacataaggtcttgcaaaagtagcctacgagtattacattagccattcggtgatcaagttattttacacaggaatcatgcttaatgttgagtttggaagttcgtaataatatgttagttgcaattaaactgcaagccttatcgttgcacgaagtatattatataccagactttactatatatagaatcagaaagaatacagttacgaatccccgacgaaacaagtttctaatattaaatcgtgaaacattataaaagtgaatggattaaacttatcacgcttaaattatatcaagtcttcagggtatatacttttcgccggttctaaattatttttcgcaagacttgttttttatacaaaatcttaacctaaacgaatgtgcagttataggtttaatattatgtgttacttgcaattacagtgcaaggagaaccgttgcaaaattacaaccataagttgcaagatattaagattatgtattgacctcatatattttgtatttcagaaatctagccggtagtttgacatatttatacg

>Hand-related -2053/+30 based on Davidson and Levine 2003

5’UTR START CODING SEQUENCE

taagtccgtcaaagacactctattttcgtcactttaaatttgttacagacaaattattattatccttcttaaagtagtaaccggttcgcttatcgatataatttcgaatctttagagaacagtccttaagattcgtataaactcatcgcatcgtttgttttgataacgagtggcataggcctcattacgtcataatcaattagaacgacttccttattacgtaggatctgtgaacaatcaaaggtttgactttacacggttacagtttgaccttatggaattctgagacacgaatacacataaaacaaacatacactacaacttgaacaccacaaccaatatttaaagtaaaaattattgggaaaaacttgttgcaaaaaaaataactcgtttcgtattaaatattttttaaaccacactttactgcatttgtgtaatatttaatttaaaaagtttccgtagacatttaagtctgcgaaattttattacgattattttacatttatgtttatacagccaagtttttggccaagatagttcagtgtgttaaaatcgacgccggttttgtattatatttagcccagcttttacacccgccacataatagtaaaccttgcaccaaccttctatccgtcccattcacctgccttcataccagctatcatggcgggttaaacggctcgcatttaaccgctgcctcgtaaaatgtgacctcgtcagtacttccgcattaaaaggggcgaaaagttaccacctcgttaagtaggaactgaaactaatataacgttttgaatcagacccagtggtctactatttaatgtctaaattgtcagccatacataaccaaaattttaagaatttccacaaaggtatatcaaggtaactcgtaagcgggcacatagtgcatgaaacagaaaacccgtgttttaaccactctcattgccgcgccaaggcagaataaagagctacaatttattacattcatttaagtgtatacgatgaataaacaacagttaaagttacgagaattaacctaaacttttaacgtcataccaaactggaaaaaacgaccctggtatacggcaacccgttaaaaacgtttgacctgatttttttaatagtaacagttatacgattaggtttagggccgaaacaactgtctaaaaggtttgttaggtttaaattatatattgctatattccaggcgcttggaaacggcattcaaagcttttaaggggttttataaatcacttaacagatatacttatttacgtatgcgaaaaaaattgaaaccctaagactttactaataaacttaatagttagaagcgcgaatttatcctaaaaagaaagctgaaactatgaaggtttttttcatattgatacatgatagtctttactgcatttgggatcgccgttaggacgataatcctccacagttatctaaacaagtgtgtctcccctgacgcacaatggttgcaacatatgtcaaaattgagcattacagtaaaagttgtttcagtcatttgtgttatctgcagaacagactgctcggtggggaattaatattagcttctcaccgtcctactatgggattgcttctctcgtctcgagatactctgtttgcggtgcacctcgcacaaaagaacgattaatataatgtaaatagtaagtaattccaaacgcaaaccgaacaagcgtcgcgttaattctcgaaacgatactgtgacgtcacagaagcgttttcctaattagtcgcatcgccttttttcaacgtcggacgatcgcgtgaaacggtgtatataaaacgaaggcgcgttttaatttatctaagcagagttgtagagtcatggtatagcatagagcctaattggtatctatatagccgtatgtgacaaatgcgttaagaagcagaaaaaatacgtttagtggcattgtttggtgtggttagccggggacgcagttttagtttcacgtggcaaatttaaagtctggtgtgaaatttttcgctattttaacagttttactttcacagcaaCACAGTAGCTTAACATGACAACAGTAGTTATGCGCACGATGACG

>Saxo intron 1 + basal promoter of FOG (-1)

(bpFOG from Rothbächer et al. 2007)

gtaagcagttaaccaaagtgtggttttgtacaattttaggaattgaatcattacattaggcttcctttaaaaaggctattactggccaaatattagaattgtatatatgcttaaatgtttataatttaatcctaaactgctgtagaaattaattaccatgtaacctatacattaaaatatcaaatttgccataaacaacgatttctcatacatttttagtggcaatttattttgcccaaactcgcgtgatgccaaacgcccttgtataaccttcagatattagtataacagcttatacaagcgtcccatttgtgtagcgagcacaatttcctttcagtaccattcaatactcccctcatttaatgtagcaaatgtaaacagtttttaaagtcttcacaaaagcgccagactaataacacagctctttccccaagctccaacaacaacataccgtgttctggttttctctttcacgcgatttcaattaaacgataagaatgcatttatcctgcttgcgtagactatttatagttcggttttgataacatgtaccgtgttttatggtgtgtagtgtatagtaatttattttaatacacgacacgccaagaacggccgataaaccaagtttttttttctatttctggcagaactaattgaatagttcataaattgctgcgaaattttaagtggggactgagtggccgcctatgaagaattcatcataaagttatatttaacgtattgttgtaacgtccaagttgttgctggttaatgggccatttcgtgaatcgtagacgcatttcattcgttttcagtcacatatacggttttagtatgaatattttagcataacagattgcgcgacctaattatcgtattggtttaaatgccgtagttacatcaattattcacttttctgcttttgagtggccataagagaccattcatattgtctcagctcgagtatctgcaggtcgactctagaggatccggcaaagcttcgtgtattgtaccggcccattgtcaatcatgcaaacttgatattatattgacaagagaagaaggcagtttaaattaaaactctaaagtagagagacattaatctcagctgacaaggcaggtggtcacagtaagttcatttaaatagttggccaacaatagcctttccaagaaagtatttttgttccaggtctatacaaaaataacacacaac

>Saxo intron 1 mPOU4 site 1

gtaagcagttaaccaaagtgtggttttgtacaattttaggaattgaatcattacattaggcttcctttaaaaaggctattactggccaaatattagaattgtatatatgcttaaatgtttataatttaatcctaaactgctgtagaaattaattaccatgtaacctatacattaaaatatcaaatttgccataaacaacgatttctcatacatttttagtggcaatttattttgcccaaactcgcgtgatgccaaacgcccttgtataaccttcagatattagtataacagcttatacaagcgtcccatttgtgtagcgagcacaatttcctttcagtaccattcaatactcccctcatttaatgtagcaaatgtaaacagtttttaaagtcttcacaaaagcgccagactaataacacagctctttccccaagctccaacaacaacataccgtgttctggttttctctttcacgcgatttcaattaaacgataagaatgcatttatcctgcttgcgtagactatttatagttcggttttgataacatgtaccgtgttttatggtgtgtagtgtatagtaatttattttaatacacgacacgccaagaacggccgataaaccaagtttttttttctatttctggcagaactaattgaatagttcataaattgctgcgaaattttaagtggggactgagtggccgcctatgaagaattcatcataaagttatatttaacgtattgttgtaacgtccaagttgttgctggttaatgggccatttcgtgaatcgtagacgcatttcattcgttttcagtcacatatacggttttagtatgaGCattttGgcataacagattgcgcgacctaattatcgtattggtttaaatgccgtagttacatcaattattcacttttctgcttttgagtggccataagagaccattcatattgtctcag

>Saxo intron 1 mPOU4 site 2

gtaagcagttaaccaaagtgtggttttgtacaattttaggaattgaatcattacattaggcttcctttaaaaaggctattactggccaaatattagaattgtatatatgcttaaatgtttataatttaatcctaaactgctgtagaaattaattaccatgtaacctatacattaaaatatcaaatttgccataaacaacgatttctcatacatttttagtggcaatttattttgcccaaactcgcgtgatgccaaacgcccttgtataaccttcagatattagtataacagcttatacaagcgtcccatttgtgtagcgagcacaatttcctttcagtaccattcaatactcccctcatttaatgtagcaaatgtaaacagtttttaaagtcttcacaaaagcgccagactaataacacagctctttccccaagctccaacaacaacataccgtgttctggttttctctttcacgcgatttcaattaaacgataagaatgcatttatcctgcttgcgtagactatttatagttcggttttgataacatgtaccgtgttttatggtgtgtagtgtatagtaatttattttaatacacgacacgccaagaacggccgataaaccaagtttttttttctatttctggcagaactaattgaatagttcataaattgctgcgaaattttaagtggggactgagtggccgcctatgaagaattcatcataaagttatatttaacgtattgttgtaacgtccaagttgttgctggttaatgggccatttcgtgaatcgtagacgcatttcattcgttttcagtcacatatacggttttagtatgaatattttagcataacagattgcgcgacctaattatcgtattggtttaaatgccgtagttacaCcaattGCtcacttttctgcttttgagtggccataagagaccattcatattgtctcag

>Saxo intron 1 mPOU4 sites 1 & 2

gtaagcagttaaccaaagtgtggttttgtacaattttaggaattgaatcattacattaggcttcctttaaaaaggctattactggccaaatattagaattgtatatatgcttaaatgtttataatttaatcctaaactgctgtagaaattaattaccatgtaacctatacattaaaatatcaaatttgccataaacaacgatttctcatacatttttagtggcaatttattttgcccaaactcgcgtgatgccaaacgcccttgtataaccttcagatattagtataacagcttatacaagcgtcccatttgtgtagcgagcacaatttcctttcagtaccattcaatactcccctcatttaatgtagcaaatgtaaacagtttttaaagtcttcacaaaagcgccagactaataacacagctctttccccaagctccaacaacaacataccgtgttctggttttctctttcacgcgatttcaattaaacgataagaatgcatttatcctgcttgcgtagactatttatagttcggttttgataacatgtaccgtgttttatggtgtgtagtgtatagtaatttattttaatacacgacacgccaagaacggccgataaaccaagtttttttttctatttctggcagaactaattgaatagttcataaattgctgcgaaattttaagtggggactgagtggccgcctatgaagaattcatcataaagttatatttaacgtattgttgtaacgtccaagttgttgctggttaatgggccatttcgtgaatcgtagacgcatttcattcgttttcagtcacatatacggttttagtatgaGCattttGgcataacagattgcgcgacctaattatcgtattggtttaaatgccgtagttacaCcaattGCtcacttttctgcttttgagtggccataagagaccattcatattgtctcag

>Ddpep -1690/-1

agtgcgaactacttaagtgttatattcaatgaagaataattcaaattgaaccactttgctttataaactcaagttcaaggggtagggatgatacccattagggtttcaagagcattttgtgattattagccaagtgcttctagtccatacaaaatcaatatcaacttcaaaatcctttttaaagttttataaaataattgaataaaaggacatcatgtacttatgtttaatctgtctagacatatgaggtgtgtaacttttttattcaaattgtagccacaagtgactaaataaatcacttgataacatatcagtagcaaatagctgacgacattttaggctaaagggcttttattgcgtcagtccatctcaggtacagccgtgtgtaaacaatcacattctttgatcatcgcttaaccttaacgcatggttaggtgaaacacttagaaacactgagagggcaggtagtagttttatcattgtacatggcgtcgggaggctgtttcagtggtttcttatcaagaattatacatccgtagtaagcccatgaagcttgagtgtatttataaaacataggtatatcatatgagtaagttgaatacagcagcctcgtgttcgttggtaagactatattgtatctagtgaattaggcttaattagttataccacgtgtactccattaccaagtcggtaatattctcaagggataaacgtgccctcaaacagacagtatcttgaaatttggtagagcatcaaacagaagtagaatctgctgtaaaacgaacgatttcacggtcaagccgctgcataagatttaacattatgataaaattgccaccaaatacgacgtcaatacgtaaaatatagagatagtataaaccaaacattacaccatgaaactggtttaagtaggattgaatctgcggtgacaacatgttatggtgcaaagccctgtcaaatattaaccaacctacacgctgaggtctgtcgcagacaaaagcagcgaaaaattcaacggtctattagcaggcgagcatttggtatttgcttttgcccccacacatttcgcttgtgttgattcaatttagtttcgtctaaaccaccagcaagctctagagtcgggtgtggggtgatttatggagctgtctttcgttgaacctaaaatatcctaatgtctttttcgagttcatgaactgtttttgcgctctgtgaaaacgggaaattttatcgagtttccggtacagctagcgttgtagagctaaacgtgtgagagaatgtccgtaaaccagtttgaaccgcggctgaggtactcccttagactgtttaagaacacagttggttttgtaagacctgtcagtctcgggcttggtagagaaatgcctttttcttctgcctgcggtcgcctatgtcaagacataaggcgggctgatcgttcgcaccatacaggggtttagtcgcttcgcgtgaagagcaatgaaacaactgatttacctgatttattattttgatttataaaaagctgatcttcctttgcagatatacggttgtagaacatactgaattccgtgcattgataacagggtagtggtaaaagttatccgggcctgtttcatcgaatttcaaagtaaaagcaaactctttccgcgttcctggaagctttattcaaagcgctctattctatgttgcgtta

>Ddpep intronic element (+6141/+9016)

cagttacaatgtgggcaggtgtggttattcatacttgtacctttagcatacatactgagaaccgaatggccagatttgaatcaaatattcggccatttttaataatcagaccatattgccgcttcaagatgcggatgttgtgtttcaatatctccgtttagtttgttgggcgtgtgcgggcattccgtttgccatgataaccaaagctacgtgactgaaagcaaattttctcttccgattccggatggcgattgaagtaattagtttcttggcataaaggatttgtctttctgtcagcagtcgtgattttaaagtcgattccgattagttgaagcctgtgatagttgaacagctgtcgatagagatcgcgtgttacgcgcttcacaattatccggcgattcgtctccaacaaaagctcttgtcagtttaggagtgtttgttcaagaaaagcaccattgaacggattaattaagcccggaattttcctttgaattattaaagggaacaacccgctcagccaatgccaattattatgatgcaattaggccgtatatctcaatgcttcaaacattgtatatcttgctgtcacttgttcatgtccgactgtcaattattcctgttgtggcagacaagcagacattgaaatatatgaccttgtctatatgcagatcgttcttgatttaaaaacgatccaaatatagttaacatgtggacgtttgcaagttgcaatgatgtagaagcagaattgtgaacttattggagttttggtgttgtccaccttatgtcaaaatcaccaacgtcagcaaagtatctgattcacgtgacctcgtgaccactgatgcgtgtcgagctagtgaaaaacaagccttcagtgctggcggtcgcttagggtcagtgacaacatggcatactttgagctccactcagctctgagtaaagttgacattgcccggatatgtgtttgtcttcgtcggccaaagtgggccatattgtaatgagtttcaacttgagtttgcccagcggtaaatgcggctttgaaggaatccaaacagatttgggaaccgcaaatattatgctcaaagccaacccttaaatgcgagtgcggtgacgcgttatcagggtggtggaatctttaccacctcaacttgggattaataacctcaaagtatcgaacttaatgagcttggtttgtttgtggtgtatatgagttgagtagtttgttcagcaaacttgcacacaacacagcacctatcaatatgttgtgcatacttggctcagtaaacatatacgagccatcaatcatgaggtagttaatcggcgttgcatcatagttaccaaaccaaatgcatttttcatctgaaagagtttgttttagacagggatcaagtttcttcgaatttcttattgatgtgggctcccgtgtaaagctaaccttttcactaaaatacacaacttgatggaactttcttatcagacttcaaatctcatcattttaaagattttttcaaagttcaatggaaaaaagattttttagttcaatggcggaaactgaacgggtcagcttgtacaatgtggttggtttgataaaatacaatcatgcttatcgacattaatgcactccattagttgttgctataaatgcgataagatctcagcaaagtatgttccccttccatatttccgtattacactcccgcaaaattgccaaagctagttcgcccccttcgtagtttccatatcaactgccgcacagttgtagaacacgaaaaactttgcccttaccgtcaatatttaagttggttacgtcatcgtgacgtcaagacagcatcgcggcgagtattacaggcgccgttcattcattcccgtacatcggcaatgctgtgcccgtgtattcaacctattataaacttggtggccagattagcggtctacctaaagcgacgcggttttagcgacgggaaattacagtagcggtttcagcgaaagacgaccagaagtctttctggctttattggacccaactgcctgacagcaacagtattttataggtattccgatagattagtatagagccaactaattgctcagtgtttaacctgagttcttttactggtaaattctgaccatcgggttgttcctgaacatataaaatatgtgacagtcttgcaccagcgcaatggttatagtcgataaagtatattatagtttactaatgtatagtgtaaccacaaatgcagtgaaagcaaaatataacccatactttattctatgctgcgtgtaaaaaacaccagtgtcaccgtaagccaaaacccaacgaccgatatacaaaagtcacttggtaccagtttacccatgcaaggattaatctaacgttacaagcttcaccaaagctcgctaccgcttgctacaaaacgtgacaacaggatcgtgtatcttgtgtttatcaacaccatcgctatttcttactacgttgaatgcggtgtcactttccctcagggctcgatcgtatcttacttagagtattgtggctttaaattacagttcaaggttttctattcgggctaacatgtcaaaaaaaaacataaaacatgacacacttttcgttttcgtacatttttgttttgaaccgaaatgtttgaaataggcattcgtttcatatatatggactgttatcctgtattgaaactgacgagaaaatccgtttttctatgcaactttgccacgatgttttgtaaaacggtggcagactttatagttaagataggacgatgaaattaaaactatgatgacgtcacccaaaagtgacgtcatttgtgacgtaaccacccgactcaccgttgcttctcttgcagcaaacgccatgttgctgacatgctcatcactgagaaggtatccagggtgaagc

>Dmbx –3489/-2158 (Defcab -1798/-467) + basal promoter of FOG START

From Stolfi and Levine 2011

cactcatctgcctacattagcttagtattttagttgaataaataagttccaatattttcacaacccgttcttaactatagaacaagaatcgctacggctccgatgctcgatttcgagtacccggtcgccctagcaacccacttcacaatagtaacaatgagacccgaccctcccattgtgaggtaatgtttctttcattgttacgtgatatcgaaactccgccaatcgcaagggaaagaaaatcgacagggcacaaaatgaaatttttcgaagagttaaaaacgataaaaaaacaaactaaaaatactaaaagttgttgattgttataaaaacgcgacagaagcgtttattgttttatcacaatgacgattaaccggaccatactatgccattgaaaccgcctaattgaaatttcgtggtcggattcgctattttcgcaccctgcgttcctcaatttgattgtattctacatcgttaatatttaatgaaatattgacttgttaatcgacacttcgcgaaagaacaattaaatcgttatgacgtcatatgggtagacgattgttttaatcgcgatgtttcgacaccacctgttaaaatgtttaacgaaagaaaatactaaattaaactttagtactttaaacggcaattgtttcaacttcgtattcacttacgtcttaggagttgtctaatttgtcagtcatacaaaggggtaaagccaaaaaataatcactcacatagttacatacatatggtaactcgtaagctggcacgaggtgtatgaaacagaacactcgtgttatatacgacgttggtcataacgactgtcgttttccggccacgcgaggataaataagttgcatttactcggcccatttagtataaagaagtaaaaaacgtttttataataagcaaaaactaatcaattaaaagaagtttttaaacagatcaggttgatcgtatttaaccatggataagaatacaattcagccacataaagtttcgaatttaaagttgatttattttaatttattcgaattaacaagcagtcgccgataaatcgaagacacatttgtgacgtcacgccttaacaatccgctattgatgacgtaataaaggtgaatcatctgccattatgagctcacaattaacgcgtctgatctgcgacaaacgaagcgcgaaattgctcgacgcgctgggaataattaacctcgctttaacgccatctagcgagcacgaattcctgtatatttaggagcgtaaataaccagaatactcattcgtggttaacaatcagcattgttacgtcaataaggattgtaacgtcatagtactctcgagtatctgcaggtcgactctagaggatccggcaaagcttcgtgtattgtaccggcccattgtcaatcatgcaaacttgatattatattgacaagagaagaaggcagtttaaattaaaactctaaagtagagagacattaatctcagctgacaaggcaggtggtcacagtaagttcatttaaatagttggccaacaatagcctttccaagaaagtatttttgttccaggtctatacaaaaataacacacaacATG

>Dmbx -2657/-2158 (Defcab -1798/-1299) + basal promoter of FOG START

cggccacgcgaggataaataagttgcatttactcggcccatttagtataaagaagtaaaaaacgtttttataataagcaaaaactaatcaattaaaagaagtttttaaacagatcaggttgatcgtatttaaccatggataagaatacaattcagccaattaaagtttcgaatttaaagttgatttattttaatttattcgaattaacaagaagtcgccgataaatcgaagacacatttgtgacgtcacgccttaacaatccgctattgatgacgtaataaaggtgaatcatctgccattatgagctcacaattaacgcgtctgatctgcgacaaacgaagcgcgaaattgctcgacgcgctgggaataattaacctcgctttaacgccatctagcgagcacgaattcctgtatatttaggagcgtaaataaccagaatactcattcgtggttaacaatcagcattgttacgtcaataaggattgtaacgtcatagtactctcgagtatctgcaggtcgactctagaggatccggcaaagcttcgtgtattgtaccggcccattgtcaatcatgcaaacttgatattatattgacaagagaagaaggcagtttaaattaaaactctaaagtagagagacattaatctcagctgacaaggcaggtggtcacagtaagttcatttaaatagttggccaacaatagcctttccaagaaagtatttttgttccaggtctatacaaaaataacacacaacATG

>Seldom -3004/-12 (exon 3 start codon) from Johnson et al. 2023

ccatgatcgctgcaagtttaggttcgccgtttcgcactcatgttaccgaaccacctaacctcattacactggtgccataaatctgtaaagttgtaatttatctgtcaccaaaaatgcccctacgttttaggtaaacgggttatttagctataaaagaagtttttgcttgcggccgtcgtaaggaatccttgaccgacgtataataaagtttcttttcattttttaatatgtcatcgtaaaaacaaggatgaaactgtcgtaccacagctaaactcctatttttgccaatacagttaaaatctcctgaaaataggactgttttgccaattgatttaggagtcggacctacgcactaaaccgcgcttatatgtgccagcataatacactgcctttcaaagtcgtatcaacagaagcctatccctaaaggtagaattagcaaagcaaatattatttctgacgcggcctggtagcccttcttttacagacggtcaacattgtctttcaaataataataagtaagagcaaatacgaatcagatggatgacttagtgctgccatggatataaaatattgttctgcataagttgataaatataaagttttggtttctaaaacagaagtatgacttttttaatttttatattacaaattgttaataattttcgccgttcagtgtaaatatgggggcggggggaaacgggacaccttcagcagataatatccaaatatcataaataacacaaaataagatcacactttacatcaactaccaaaagttaaaggacaattagtccctagttatttagtaggttggggggagatggtacacgtttaacacattttattcaaatatcttgatcgtgatttaaacaactaacaacgttctatgagagtagcggggataaacggatacggttttataattatttgaatgtttcttgtttaccaccaaatgggacgagaaaatagaatgaaaacgtgtcccatcttcaccctgctgtcggttttaactttataataaaaactatttctccgaatgaccgaccgtcgcgcatctttactacgacgtcggttaaaaacttttttgctacatagttaacataactatcttcatgtatttcgagataatgagagataatatatcgccatagcttgttcatgaagtaacatgtagacaaaccagactactcaactcccgtaacctcacaaactgcaaacacttatgtcgttcaagtatgtaatcaattgtactgtgactttcggttaatgtaacgcttcgcctataggcctacagcggttaagtaatatctaatttcgggttaaatttgccgaatccaatatattcgtaaacgtgactcccagactgtcgcaagaagtcaatataaatatctcagtggccctgtaataattctaaatcgagttgaatcaatttggttcgcaaccttatcgctttttattctggataatctaatcactggcgtttgtgtacgctgtttgtgggcgaggaactcgctggcctttctataaaaacacaactcacaaagacctggcaagcggcgtcttttgttgctaggtcacattcagaccaccgcgcgcggtatacgtgttgcttttttctagtacagccaaagcgattctcttcaaaatacagtcgagctacacttcaattaaattcactttcaagcgaatctacatcgaaactaatgaattttcaacattaattagtagccaagtttgaaactaataggtcacacttgtattttggttagtttttatccgaacctttttaaataattttttggctttcaacgataaataatatgccgagagacgttttataatgaatccaataaagaactcaggcttataatagactaccaaccctaacagttcatgaatcatttatataacccgaatgtatacacacgacacacctatacgtatttaaaaacagctcactacattgtcagcgtttaaaacatttttaggcttagtttattacaactttgactaaatttgacttcacctttgattacatgtcttcaaattttggtcgttttcacagtaatagaatttcgataactccaaacgaatcagcaacataaatatgcgggctcactcgtctggggcacaactcgtatgcaaagtttaaaactacgttcaaatgttaacttgcatccggattgtgacgtcagagaaaaacgtaagttacgttttttgcttgtctatatttgttaaagcgtcacgtttttattcaatttgttgaaaacaatgagcgattaggtttagaaaagcaaacacgtaaagataaatttcgtttttttgaccaagtcaacgaaagttaaaatagtatatacataaaaatttgctaaaacacgctaaaaaggtataaaacttggatacagaacgtaggacaaccgttatacaataagaaattttattaatgtaacattattgggaaactgtggaaaaagagtttagacacatgactagtacaataatgggaactaccaaatttattttttttactaatgaaaaaatattatcacgaatgaattacgtaaacgagttttctccatacaattaatttcagatcaaaagcggaacggatagtcgtcattgcattaataattcatacacatggttgacttatacccgtcgtcaacatcaataattgatcttaactgtcgcggacataataaattcggttgctcgaattttttaaggtaattgactcccgtgttaaggtatgcgaatgttggctgtttagtgtctgctagcgtggaaataaattctttctcatgtttgctggttttcagtatagtaaattagggagattggattttgttgaataagctgaatgtgtaaattcagtcactaaatttaaataggatgtaatgtgtctaattcctaatatgttgtcgattattcagcgactttttaatgatttaaatttcagacatttattaacacggacaaaaacttagtactga

>Defcab -1792/+24 START coding sequence (from Gibboney et al. 2020)

atgacgttacaatccttattgacgtaacaatgctgattgttaaccacgaatgagtattctggttatttacgctcctaaatatacaggaattcgtgctcgctagatggcgttaaagcgaggttaattattcccagcgcgtcgagcaatttcgcgcttcgtttgtcgcagatcagacgcgttaattgtgagctcataatggcagatgattcacctttattacgtcatcaatagcggattgttaaggcgtgacgtcacaaatgtgtcttcgatttatcggcgacttcttgttaattcgaataaattaaaataaatcaactttaaattcgaaactttaattggctgaattgtattcttatccatggttaaatacgatcaacctgatctgtttaaaaacttcttttaattgattagtttttgcttattataaaaacgttttttacttctttatactaaatgggccgagtaaatgcaacttatttatcctcgcgtggccggaaaacgacagtcgttatgaccaacgtcgtatataacatgagtgttctgtttcatacacctcgtgccagcttacgagttaccatatgtatgtaactatgtgagtgattattttttggctttacccctttgtatggctgacaaattagacaactcctaagacgtaagtgaatacgaagttgaaacaattgccgtttaaagtactaaagtttgatttagtattttctttcgttaaacattttaacaggtggtgtcgaaacatcgcgattaaaacaatcgtctacccatatgacgtcataacgatttaattgttctttcgcgaagtgtcgattaacaagtcaatatttcattaaatattaacgatgtagaatacaatcaaattgaggaacgcagggtgcgaaaatagcgaatccgaccacgaaatttcaattaggcggtttcaatggcatagtatggtccggttaatcgtcattgtgataaaacaataaacgcttctgtcgcgtttttataacaatcaacaacttttagtatttttagtttgtttttttatcgtttttaactcttcgaaaaatttcattttgtgccctgtcgattttctttcccttgcgattggcggagtttcgatatcacgtaacaatgaaagaaacattacctcacaatgggagggtcgggtctcattgttactattgtgaagtgggttgctagggcgaccgggtactcgaaatcgagcatcggagccgtagcgattcttgttctatagttaagaacgggttgtgaaaatattggaacttatttattcaactaaaatactaagctaatgtaggcagatgagtgagtggacaagcagaggggttcgaatttagcaaaagtttgcagcgaaacaaggatttgggttgccgcaaaatggtttataaagaaacagtgcgattattagaagcagcaaaattgacatttaagaatttaaaggtagtatgcttatagtgcttaaggttataaagtcgtttttttattaaatttaatttataatatgtgatattatgaaagttaaatatttagaaatgcaacttactacagatttaggcgtcaataaaaaaaaagaaatgaattaaaaatccaaaaatatgataggcctacaaagtttgggctgttgtagttacgagatagattacgcaaaacagacattagttttgtgtagcatataacagctacggatgtttagaatataaagaaaataagcactaacttaatttttgttcaaaggttgcgcttaatttgggcgaaagtttcttaaaatagaattATGTCGCAAACCATACTCAGTAGA

>Ebf -2631/+15 “STOP” START from Stolfi and Levine 2011

attcttccgggaaataagagcggcagcgactttattcgaatttcaaatttccatgatattcgataccccacactccaccctataacggtgacctggtttaggaaatatgtgacgttgcttcacgacaattttagcagctatataagttaccattgcaaaaagatcagtctttttatttatcacaatttttaccagatttttaaatattttttagaaataacttaaattttgatcaaactgggttaaaattgcaaattttacatttttgtgattaatattaatctttttagtcgccatggcttaaaaatgtcatttgtttagtccgctgtattcaattttaccgtttgataaatcaaacggttaagtttcgcataaaaccgatttctcatatggatattactccgcgcgagttctaacaacgcgttctcttcgctggccaagatcgatactccgtgacgtcacaatcacatgcgcgcgcgtctgaaatatggtcgacagagttgcaacacgcaacccttttgtttgggttaatttccctctttgttttttaagtccgatcgcgtcgccacatctgttcgacgaatcttcgttctgtcttagcaactttgcggctctgtccattgtgaagtcgtaaacgaggcattgtcgtcactgctgtctcgcctacgtcacaaagcagtgcacagtgacgtcacggagacgaggtcgcgtgccttcgagttggaaaattcaaaccattgcactttttgccgaatgttaatttttacgacgggagaagttaggaaaagcataaaataacaaataaataaccaataaaaaacgtaaaaacaaacaagagaacggtattcaaaattagaatttcgaaaaaaatgttaaaaaataatacttagagtcgctgaacaatttaagccgacaaaggccacaaaactgcctaaaatttataaaaaaaaatgtaaaattattttgttttttttaaactacagttatcacctttaaaacaaacaaattagcaaacgttgtaattacttcacaactttcttgcgacgctaaaaggcggcgaattttattgctattgtgacgtcacaagcgctctcgtcacgcccggatacgattagaacaacgaaggattgtttgtttttaattatttctctgtttaatcatttgatttagcgcggcacaaattttgttttatataaaatgtatctatttatccattttatttctgtgcgttgttgtactattttttgaaaaatgtttgttaacctttagaaaatcgcgaaaccaacgaaattatttctaaaggctgtacaattctttttcgttaggttacgtgtttaagtataggaccaagttttaaatggcggacagagtttcgtattttgatatcttgaattattttgaaatcttgaaaaaaaataattgtacgctttacatagaataactaaccaaaatatctgaagcaaaaattacgtacaattttttaaaatgtaatttactttctagcttttaatttttgtgcttttctcaatatgtgtgccatattttaaaaacgtaaatttgcctgttgttaagcggaggaaaaagtaatctgcgtgaatgcgaaaaacacgattttggaatcagcgccggcaatggtgtttgtaaataggggtggcatacgcgtttcgtagcgaaagagagaatgaggcgaaagtcgacagatgcacgctccgatttatgagacaggaaccagtcgcgagggcacggaggaaaaagaaccttactccaagacatgcgccgccttttttctttctgtcctagtcaggaatactagagtatagaaggccacgcgtcgtggagtttaaaaccagcaagcgagtgtctcaacggacacatcaatacagagacttctctcagtggacaactcggacgattcgccactaacttggtggattcgtcccggacgacccatgggccgggtcccagcgcgctagttggccaccatacagtgtagaatcagctagatcgtctctgcggatttcgcaaatagattgagttggagataggttcccgaccgggatttcgactaatttgcaatgttagttattaatcaaggtgacagtcaggagttaaagttaatttaccttttgaaagggcaaaaagattttcgaagtaaattgattccgttaattgtaactcttaaacgcaaaccgataaacgacgccattttgcttttcattgagaaactaaccatttaggcattctataattaaaattaaatgtttttaaaatctgtaactatctgtaataaactagaattatttatttcagtttaaatttttatttaaaacaataattttattatttcctttattcatcctattgatattggtatagagaaaacgatgtttttatttccaaaaaattttgtttcataaaagtcattttgttcaatttaaaaaaataccttattctaaaaaatgcccaaaacaagttctttatttcttaaaaactatcctaaataaaaaaaaactccacgttttaataaaacctataaaattaaaaactataaaaagatgacttatttttttaccctaacgtgatttttcaccagataccttaagtgttattttatttgtaagtaatatccaaATGGCAACAATCGCGTAA

>Engrailed -2086/+24 START from Stolfi et al. 2011

ggaaagatgggacacgtttagcacataatatccgaatatcctgatcgtcttttaaagaactaacaacggtctgggagtcgtgaggatgcagttttataattctttgtatgtttttgtttactaccaagtgggtaccaactgggtcaagaaaattgaatgaaaaggtgtcccatcttcccccaccctactatatttatcaaacaattcgcgaaacacgtctatcataacacttagaacgaaccgtgaaaggtcacatttatcagacaaacacacgccgcctaaattcccacagcatcgtgttaaatcaatcgctaattactgctctaaatgtgaactgatctataggaaaagtggcaacgattgttccgggtcgttgttgggcgtgtgcgcttcgatagttcgcgataatgaacgatgatgtacgtcaaattattccacaagtttttttttttgtgggttttaaacacacacaatgtacatccgacaaatgagaatttaatgtagcgaaagttaactacagtttgaacaatcgctagatttctgaccatttgcaaattcaaaccgatctgctggcaaaattgtgatgaactccaatctataatgccagttgtgaactttggccaatattaattgaagcgtcgtttaattctcgattaattctgcggtcgttccgattgcgccacggtagggaattcaaaatccaatctggatcagaactcttaagttacttaatttcaaatttgtcactgagacaacacacaagaacaaagtcactccaactcccataggaatgatgtttaacatggctaaaattgaaccagaaagtgattaacgcagccagattgaatgttatacgcttgaatgaagtatttgaacatttccgaagtcagagttttgttggtgtattggagtgggcggtatataggagaaggcgatacactgctgagaatgaataccgtcagctaataatattctattccaatcgtattttaacaaataataacattttttggtggtggggggagacgggacaccttcagcccagaatatccgaataccctgatcttgttttaaacaattaccaacggtcaatggtagtcgtgagaatacggttttatatttctttgaatgttatttgattactactaaataagacgagaaaatagagtaaaacggtgtcccattttaccccaacactactatatacaaaacgggcacgccgctataacactgaacttagaaatctaaagatggaacaaactacacctccaatcaacaaaccttaagagcaccttacacgtattgattacagaccaccatcattagtgccacaacaataaattgatcagaaaaaagctcttattgtacaagcagctcaatccaaattacagtgttttgtatgtacccgagaataaagtttaaataatgtgtatttcgctacttaatttatacaaataattcaacgatttaatagcctagttcagcaacaaatattttcctttcaatttgttcagaattcacaaaatgacatttcacgcttcttgagaacaattacgcaatcggtttaaccgagtcaacgaattttacaaaatggcggaactcaagaaaaaggcggaagctcagattggaagcggcattcctagtgatcaagtcagttacagcatactcgtctgtgacgaagtatatttgtgtgaaggacgtaaatgatgtgtaaactttaagcgtaataataattattgcttaaagatttaagttgaatttatttaaattagatacttgtgaatcttggtaaaatatctgtctttgtattagatattgttggttggacttaatttatgattacttctgatttaatatattttgatccattaattctgaacgccaattattttttcgcgcaaaggcagagcgagtatattaccttattttttaataagaaaaatactatttttctactaaactatttcaaaatggtaactctttttttactttcgaaattattttagccgtttaagtagtaatcgtaaaacaacaccgctataaaagtgttttatttcaaaagacaaaggcgggcggtaaatttaaaATGAAGCGAAATACAAGGTTGCCC

>Pax3/7 -5873/+21 from Kim et al. 2022 START

tgacatacagtttctgattgtcctcataatttactgttaatttggatttaattttataaatctcgttaccgttgttttgtacggaagtaaaatcgcaattattttcacggtcaaagaatccgttctttcctaatcaagtgtcgtccatgtcgttaagcagaaacagctatctacgggctcggtcaccgggaatgaatgcgaggcgataacttttcaaactcgtcactaatttcagtaaatccaataaaacgagcgctaagaaggttcgggggtcggtcggccgggtcgctgagaggtgacgcgaaaaggacgacccagaaatccaacagacgcctcgcaatacgtcacagaggcattgtgtacgtcactgtgatcattgttcgcgttagagtgcttgcgcgccgtcactttacgctcggttaaatttacgtcgattttttttgcaatgatagcgaaacgattcgacgcgcacgcgctgctaatcagcgcttcggtcacggaaacgctaaacactcgtcagccacgcaacgaggcgagcctaaagagtccggaagagccggcttcgttaactcctttgtttaaacgcgttcgctatttcgaaaacggaagccccgacgacgtcgttcgcgggcccagcgtgtcgtcgaagtagcgaaggtaggtcgggcgtactaccattttaaaacaaggtagacggattttcgaatgatagaactgatcgaaaatcgattaaagactagaatacgtccggctgaggcgatattcgcacacaaaaccaacattaaatcagcgggtgttgtagaaaccgaatagaatcataatccgttcacataaattgattttccttccatatggtttaatagcgtggtcgtgcattcgattttcgattatgtgaacttgcctggcgttcccacgcatggagtgttggtcgaaaataaacgattgtgacatcagaatatgacaaagcgtcaagaaacaatagaaaacgcgattcccacaatctccagcttctgcgacgtttttaaaaattcgtcatcttaatttcacgacgattaacgaaactttgttaaagcgttagggttactttataagcttaaacgcaacaggaacaaacgagaaatccgtaaaacggcttcgaaactcatcggtgcgcaaacacattaagagatttaaatttgcaccgaatatattttgtgaccccaagttaactaggcgcccgatcatcctgttacgtcgaaaccagccgaagctggatcgaggtttgggccaatttaaatgtctgctgccacacaaagaatgctcggttgcagtttcccataaaacagtggtttaatcgggaacccgctgagcccgggcgccgtacgatcctgtcagtttcgatcaggcagcacccctggttcgaaggtcacctgaaaacacggagcggggtctcgcttggccagccatgccgaacgcttaaggaaacaaaatgattatacagcgagaagaagccatgttggctctggacgtggagtttggaaaccaggaattggtttcacgcaaagctttcacttttgttgtcaattagacgctcgctgacttgagtagctcacccgctgcgagatcgagcttcgaaactcgactgacaggcgaacaaaataattataaaggcgtcaccaaaaattcaattgggaaagaacacggaaaagttgaatttgcgcatttggaaaatatagaactttgcttgtattttaagagcagttgcacttttaagaccacattatcgacctgtatgatacatatgttgggaataatgtcgatattgccaactcttaaaacgttatataacaccctcattttaaccgcgccaaactcttttaaattcccacacccgattcgcgctacgaacaaccatgacaatgattatgattgatttacatcgcacatggaggtacagagatgacatacacgcttgcttgccttggtttataaccattgctattaggtcgggtaatgcgcaagcctttcatctagcacttgataagctgtagttctaattaattaattataaatacgtcaaatatgatgattttataatcgaaaccgaaaacggtttcaaaaaaatgtcgaactccgcagtaaatcagttaatacgaatatttagataataaaagtaaaatatgtacaaagtatctattaaagtcacaaatacacacggtataacaatattaataaaatatatatacatacctctaataatttacagttagcctatacatcgtttatttgattttcagaaaactgactcctacaatatttatctaaagatttgcttcctatttttgctttattttacgtgttgcttttttgctgtaacttcctatatttatgaggcaagcatgtatgactgtgtaaatctgcttctggccgaactgtgtcctttggttatcaaatccggtattgatgctctcttttatgaatcattctgcgtcactacagataacataacatcagttacctgcaccacacacaacaactataacactgaaaactgtggtatcttctttacccttatattcttcggcgccctaacgctatcggatttcaaatttaaattatcacgcgtcttagtttcaacgtttgcaaaatcacggtatagtgttaagacttcaaaacaccttcacagtgtaggttgcgtgaccaatcgagtcttagaaacctcacggcaggtcttccacggcgaataagttatctcaaatttaggtgcgaaaagctaaagggcttattttcataggtgcctccatagaactgtggcgaatgcattgctagcataggggcttacaccagccaaaccattagcttaccaattaggcttatggttatcgccaaaatctcagggccgataattttcagtttaattttagagttttttctgaacttgaatttaaacggtgcagttccagtaataaagttgccagttattcctaagttttgttttagttacaacttttattttctgcttatttgtatttttttgtataaagtttttggcatttaaaaagccatttcgtcggtctaaaaatttcacacatcgttctatctgttcaaataacagaacacaagcattctttactgcttgatatacattgtaacttgaattatagcatttttcatctctgttttaacactataatgtttttaagtgatttcatccaaaaatggattgtggttataacagatctagataaagttaagccggttactggaatactatgctcggtgcgccatttttagtgtattttgtttgttaaacctcgggtaaaatgtgttcggagtgcctttatgcttttaaagcgggccgtacattatacagcaacgatattgtgtttgggggagagttttagaaaacctctttcgttcaattgtaagtataccgtcccttctacgtgttgtttgcacacgctgaaatattgcaccattttaacaagcttgactgaagtataatttttaaacgccgttcaaatcgataattatacgccgcgtcctccgagacaccattttgtagcaagcgagccaacggcgaccggccggcccgggtcaggttgatgtctttcgccggaattgacgacgcaaagtaccgcaggcgtgggtgccgaagttgtaaattggacttttcggattttagttattttcccaaatttaagtgaattagttcaatacctaaactgcatgtaattgcgcacattgataactttacacatgctcgctattgtgttatactgttttgaagacacatattctacacgcctatcacctataagcctctaatgctatatagattgaataatttaaaaggtggtccaaggcagcggtatctcaacacacagtttacttcttaattggaatttaaaactgcattgcatgtcacttttctgctttatttcacggtgtatgattaccatattttgtagcacggcgttaacgttacgtgagagaagcatagatgtcgcactacatcggcccgtggcaacgtttagcccaacacgtttattaccgtcgtccatttttgaaaagcgaaggccgatccgaaatcgtgcagagttgcaatcctgcacgcgaattcgcctaattatgtcgctcgccgttaatcaagcgtggttagggatagagaaaaagatagactggtgatatagagtggctatgacgacgagcaaagcgaggcaacggcaactcagacgagctcggtagtctggcgagtcgctgcttttttcggagctcgaccgccgcgtcgtaatatcttgctcgacgacgcatcaaaaccaaggcaatcttattggcgtgtagtggtggacggaaacggtcagcgtggtccagcctttcgttatcgaatcgattttgggccagaaagcaagtttatgtaaaaccatgatatagatgggacttagagtattaaggcattcatcggggaggcatttttaggtgattggttttgatttaaagtcgagttatgatccaaagtatgattacaaaactaaaggttgaactgagctgaagcaccttcaccgtcgtaacttaaaagagataagcggaggggacgctggaaaattcagtttaaaattagatgaagttaaagataggaaaagaggattcgctacgccgtgggtttttgcttattttaaggcatcgagcttgaaaaatttatacagtgcaccgtatagtcaagccaaagttgcgttacagaaaaactacattgtttagctttgatcggccggcagcgtccttgctgtttgtttgttgatttaagcatttgtgagaatctgttgcgccattttgaaatataacaaccttaacgaaaatttattctaaaagaattttcaaaaatattttgcctagtagtttatctcttacaaaactacaagcatcgataattacttaaacttttcataaaatacgccgcaatattttccaatcacaaaattagctccttaataaatataagggttatagcttttataactctatataaaccagagcgttatattactcgttatacatattgctgtaaacttctgataaattccaatttttctgtcctcccgctaaaacatgttgttcaatctaagcgtcactttttcctatccgttctttcaatgagacccgtgcctgtaattcgatatagacggtggttttactaagacagcatgacagataactgatattctggttggagttagcggcgagaaaactgtagatgtcggtttttaatggctttttgcggaagataatttctaaatcacagggtagttaaaaatacttaacttacataaggggtttctgcacagtttagaatagggccaataattacataattatgtaaacttgtagtatattttaaactgtactacattcgtagatgcgcctaaatcaagtataaaaaataaaattgaaaaatttacagtcgttttgcgttaaatacttattttaaaaaacgccaggtgagcgttgcatgcggacgccattttaattttacctttcagatatctcatgcacttagcttatttttcgtttaaaatcaattatttttaaccgattacggtttaatgtactagggtattaatatatttattattgttatgtggtatatagtatagtagcatttttgtcaaaaatgaaataaattcagtcgcgttttagatgtatttgaaatgttgtaaattttatttatatgtttgagaaaaaagagtgtaattaagagatggaattcgttttatttttgaacgctttcattattttgcaggccccgaattgctttacaattgcaataaatcttataagagagaaagattaaattattgttgtaaagtagtctgacttctgaacacaacccaacaaacATGATGCATCCAGGGTCTAATTTCCGA

>Slc32a1 (VGAT) -2352/+9 disrupted start codon

Based on Yoshida et al. 2004

tttgtaagccatgtggtacaggccgtgaggtatgtcgttttcgacacttcacaacgagcgtgactcgattcgcaaagaatcacatttcaaaggttttttaataggcatgtttgcaattatatggttgggaggataatgtagagccctaaatgttacattatcgaaggatgtaacgctgctgtggtatttgtatgatcattttattaccggtacgaaatttacattagcgtgcatgtgttcctaattagtttgggtccagaattcgtagcgttccgcaacgaagacttattacatgaagcttataaccgggaaagttgattgattgcaccaggtgcatctatattgggtttcagtgaaggacttttaagtctcaattattcatcgatttcctgtttattccgctccataccttctactagccgcaataacccaacgatcaaacgacgttgtcggtccgccaagcttaaaattagtgtagaaagaataccagcgccgttcgtgttttgcgatgttttatcttgatttccgaatttatatgtggcgtaagacattggtcagaaaaaggcaatcctaaaacatttttgtgttaacgtcctccccactatagcgtcatctctattttgttatataaaagcggccattttttttcaacccaatactacgacttcggcaaccaaagcagaaagttatccaatatactggttgcggtcttcataacgtcttataagtagtcgccgaactcttacaggacaaataattattctgtattgtttcagaccgacccatccataatcacatcagccgcaaagtacttagcacgaatatctgaagagtgaaaatcgtatgtattttatcgctagtttgtatagtttatcatgaatttaaactacgtttcactcgccgttgattttatgatcgcatgatgaaaatatagtcgtcaaaaatttttcatgtgaaaaatatatgttacgaaatacgaatgagttactacaatgttattccgtgagctttaataatcccgtgcgatcccctataatctatatcatcaggcttgcaggataatacaggtttggaatgaacgaaaatcgatcgattggctacgtaataatcccatgtctttctatttcacgcttaaatgggcaatgttacgccaggggaacctttaaaacgatagccaatcagtgcttaacgcaccgcagctatgcaatcgctattagtgccccatcttaagcaacaattccgattgattcgcaattaaacaccgatttgttttattggttctaaatgaaagttaccactggagaaaaaatgtatggccatgtagtctatactgcttaaaaagcgcagctgaatcggctgcagtttgatttcgtttgtgtttcgcagtagaaaaaacgttgtacaagaaattcagtaattccggaaaagacttttagtgaatctgcagcgcattccaaattattgtaggtgaatcggctgcagtttgatttcgcttgattgtccatgacacgtagtttaaataagttgccaaggatcgattccggaaaaaaaaacatttggtgattctgcagcgcactaccaaatcatttattagaaattttacatgcgcctgttttgaaaatacaatattttcttgaatcgatacgcaaggcaggttctataaataattttgcaaaatacccaagcgatgaatgaagtaattactgcctggaatgatttacagaatcctgttacagtcagagcatttctttagaaaagcagtccgttgatgttgtcagtcgtaattacggagctttttggcccaagacattttcatcagcggtattcggagaacatggttcgcctctctagctgtcaacgcacgattccctggagataagtgcgtggctctacgcctcttgtcaatgtcgctataagtgggctaaatgagcgctttattggggtatctgcggtcgcagagattcacctgcatttcgacgcattcgcggacctccagctcgcagcgttgataccgatcgtccctggtgcggattagcgtaggcaagaatatgccttgcttattggtcgctcaggcgtgtgatcctgcaatcatacgatacgaagacattgatcgaccaatctcaggctatattaaacaatattaaacatactgcatttataatagttcagtccaggctgtatattgtgctaaatagaaccaaacttaaacaggtttttattgaaaaggtagcccagttcgtgcgataaaggcttagaaaagacatagtctaagaaatatttacttcaaccagacctgcgtggtggaataaaaggttattggaataaaaataagaataggtaccttctgttagtATAGCTGCA

>Ephrin A.b -1328/-699 + -355/-15

aacacactatacgtgactgagataaataaccctgtgtaaaagcggtaattaattcacgtccaagaaaaccagtgtctaattatactcctgtggaggttcgtgcagccacatggcttgttaactgtcgtcgggttcgctttgggcatgtgggcggtttgaacagcgatgaaagaagtgtttttgagcaggagcgtggagggcgaaaagtgtcggctgcatatggtacatcgctgattgctcaacgccactgctggctttgctaagaaggcggctgctgtggcaatgtattgcaatgcacaacagtttacagattgcagcttattttgctaacagggcaatcgagcaaaaactggtctaattttaaaaggaagcaaaacaaaagaacgcaatcgttaatttgatacggaaatgaggaattacacggcggtactataaagtatcccgctaggatctcttgtgttcttgttatagagttgctagctgtaaaacaatacgatttgacccgccgcgaagctcgcctttcaaatcgatgttacttattgttgtttggttttcttcgttcgcgacctctaggtcatgatgtaaacaaagtaaagcatagcagtttaacattatacgacgacgctgagt ctcgaggaagcatagcagtaaccttgggcgattgtgtcgacacaaacgcgattaagataagatctcagtgtagcgcacagtacagtgcactgtttgagtagtagtcggttagagtgtaaaatattatgtattataatggctttaattaaaaacctaattgtacatttattcatttttaataaagtaggattgtagtataataatgagaagcagatcaacagatttatagcctgcagaatacgttcagcatttaagtctataggattgtggtttatagaatgtcagattacgatgttgcatattcaaacaaatatgataaacctaatttccttgcagattgtatgtaa

>Foxf -836/-1 (epidermal driver) from Beh et al. 2007

gagcagcgctttaattaattaccgaggtggtcgccgctccgccgcgggcgaggtagcaattatacttaataaagagccgttaagaagtagcaactttcgccattcagtcagtcgtcggcgcagtaaatgtgtagcttcgtttatggggattattcacgaattaagtcaacaagtttatagcaaaattacgagtcgtaaaggttgccgattctattcgaggtattttggtttacaagccatgcccttttacggcgttttggtaaaacttaaaataaaataaaaatatgcttaaagttatccttagacaaaaatcacgagtttgtactaaatatagcttgaacatataaaatataaggatatgtatttttgtcgtgccagtttcacacgatcaaggcgtgttaaaaataaatacagcgactagatttagaaattggagctattcttactgagccatcttggattgtttattttaaaatacagacagagtttattttgaaagtaaacttggagaaacgtttgtagcatcaagacgtgatagacagaactgtttactacgcgctgaagttgtttaatgtttgagcgattcggtttgtttgctgtttataaaactctgatgtagcggttggcctaagactgttttgtttgcctgatgtaatctgtctgctatttaaataagaacagagaaagcaagcgttcttatttttgctaaatacaaaatattttattaagtatttaggagtatttataattttaacctgtgttctttatttaaaagatttcttttaagcgcttttaaattattacaagaccagttttacaacatcagttattgaagt

>VGLUT -2814/+1773(exon 3) exons (based on NCBI gene model) START

ccggtatgtccacagcattctctttaaactctttttcgtaatttctcttttcgtgctcttgcacggagcccaaaatctcttttttattcttattgagtggaaactcgaaaacttgttttacttcctcaggagtttccttctcaatccattcttcttcttcttcttcctctttcttcttgacctccactttcgtttccaaagaatctaaaccttccatcgcatcttccaatttatcggaactaaccttgattccaacatccgacctgcgatctttctcctccacgacagaatcatcactatcagtaatttcatctccgtttgggtgcgaggtatcgtctgtttcgtcatcgtttaaattttcctgagtctcttcttcctcgattcttggtactccgtgcacatcttgatcgtcttccgcagaagcatctattactcctccgttgtcatctttttccttttgtcttctgtcttcctctttcacgccgacatcagaaaaattttcaaattcccgttcactgctttctgattttggatcaaggaaaattggttcttcataaacgactttaatttgtggtctgtaaataaatatcttaaaatctcaacgttcctcacctaaaattaggcaaatataaaagattcgtgtccaaataagagcaacagcgaaattcggaatatttgaaaatctatggataaataaagcccatacacaaatacttacgtgctcgttatcgttgttgttgaagtaaaaacagcgatcgcgacagctatcatacaaaccaaacttgcgaccactattccacggtattttgaaaacgtcggtgttgatggtgtgttactgtggggtgagatggataccgttatcacataaggttcaaaatttttttaacgtggttttaagagtcatatatacacgggtatataattctctaaacattaccaaatgtgacgataataaaaaatgaaacatggaccatcttaccccgacttacttgttcggggtgtttagggacaaaatcccccaaaatatattttagttaggtttgccctttactcaattaacatgtaacctacatcgtatatgatatggtagccaacaaatgtaggtaacttgccttttcttttccctatcaacatactccttaatttgacctgcaggtttcgggtcatcgctggtcgcctgttttcccttctttgatttctttttcgccatcttaaccagctcagttcatcctaagtgtcgggtagcagttatttgctacttgctttgttttaaacttcagctgcaatatgtgtccagcaagcgatcgtgcgtttagcgagtgacacgttagtcaaaatcggccatcaccgaacttcatcgattcgttaagtgccattcaacacttaaaatctttctgaaaaacacagaccgtcctctttgcttcacacgttgttaatattgatacaaaacaaactgtgtatgcatgtttgacaacaattaaactttaaccaaaattttacgtttattcataaaaatgtgttagcacaaatatgactaatttgttatattgttgttaaattaagccaattacattaccatagaaatttctaattagctccaattaggcgccctttatactgcatcgcgtacaatactgtggggtaagatgggatatctttatcacataatgtccacatattctaattgtaatttaaacaattattaacggtgtatggaacccgttaggatacggttttataaatcattgtttactactaaataggacgagaaatagaataaaaaagtgccccatctccccccaccatactatatacgttggttttcttgaagcaagcccagaatttaggaatcacgaaaattataaatacgacctattgaacaaacttatacagatcattaatcttattgcagctaatagtgaagctgtagtgaaagttcattagttattgcaaaccactttacttcgatttttagtcttttttcaggaacagtttttcctgtttagccgttactaagtcgcatgtacagcgaactgtttttagtgattgcaaatcgttttgagtgctacacacggcgctctagcactcataacagaaatatgttgtaatgcgcaaaagcaattaactgtgattaccacaacagcagcacaagaaggattagtcgtcaataacatacaggtcgataatggtgtttgtaccccacatccccaagcgaggtctctcgagaccaaaacattgcaattcaaggaggttaatttgccttcaatggaattattgcgcccacttgagtatactgaataagataattaaacgtcattttcgtcgatttagcaacgaagaaagagttattacgagattgctttatatttgactcaattcatatactatatacttagatgaattggtttatattgtagttcaattatttttttaagcaattgtaaacggcgcggcggtaattggttttcaataatggacattccgagttaaataccgacagacagaaaaacggaacagttcagcgttttacaacgtcagctgtccagcactgcgtcgctaagtctgtgtcagtttcacctgccttctgcacaggacagtcactcggttgaacagctcgagtaaaataggaaagaatgaatttttcataacttgcctgcaatataacaaagacgcatctctgttgaagagaattagttatttttttgctttgggaattactataggaaaattgagtttgatattttgcattaggaattgataaagtatatttacaatttatttattgttccagagcaggagggtatttaggatattATGACAAACAATAAGGCGGATGGAGGATTCGCGACGGCATCACATGACCTTCTCAGAACAATTCAAAACGGGTTTCGAAGTGTTTTATACAGgtgagagcatacctaaaaatttatatagctttttgccaaaaaaattagaaatgtttttgtcgtttggatttttattatagttaaacatcgaaaatgtcgatttggtaccgaacccctatatatactttaatatagtgtcgcacactattacgtatttaaatttcactaattaaagaatttcgtttcgaatatccgttcgttagtcgcacccttgtgtatatagtaatgctcccaagaccacttacacggcaaaaagtgaaataaactagttcccgtaacgatatgacaagtagacaaacctaaattattcatggccccgaagcaggataacattgttagagagtgagtagttctattgtgtcgcgaaatttcccgcacccacgattctgttggtcctttctcatcacgaaataacttacgcagttataatatgcatttttatttttattttcccaattacccaatcttgaaactttttccaattcatatatagtagagtggggggaagatgggacacctttagcacataatatcttggaagtcgcgaagatacggttttatgattctttgactgttctttgtttattaccaaatgggacgaaaaaatagaacgaaaaggtgtcccatcttcccccaccctactatagttactttgggaaaacgctaataccgttaacacataatcacctatatttcctaacagttaacaacacaaaataacaacgccctttttgaggataaacgattatatagttttttatgtagtttcaccttaaaatgaagattatgtagtttcaccttaaaatgaattgttcttcaaaattgctcggtcgttgttttatcgtttagacttaaatggaactttcataaaactgttgtatctttttagtaataaggaagttttaaacgattacgaaaatattgtattgttatgaattttctaaaaatcctattaatgtgctacatatatagcctacgcatagatctaatgccagatacgaaattttagaatgtgtatatgtttagaaatccaggtcaaactagctaaaactggtttttatctttttttcttatcctgttttcctcaccgttacataaatgttcatgcaattttaataagacccctgtttttgaacaactatcctgttatagttatcccgtgttatattgtgtttgtaaaactctttcaaacataatcttatttttaccgattaagattatatcaccaatatcttcatcaaaatatcatattttaataacagGTTCACCGGAATCGGCAACCCACCATACCAGGAACAAACTTTGGACAGAAACTCAACTTTAAACGAAATTTCACGCAACTCCCAAGAGCATGGAGAAGACCATTTCGAAGAGGACGTCCCACAAACgtaatctaccttgacatggcttttgtggctccatgtgtcaccgctaatgcaccaatgtgcatcaaaaagtagcgcaacgaaacaaaaaatctattttccgttagtgttaatcttgtcacaaaacttaattatctattttgtgggtgaagttcttgtcagagacccgggatctgtgccaggtttagccttcaccccacaacttttttatgcagtatttaccctgcattttcttagcatggtttataatagatggcttatgttattttttctgcagTAACAACAAGTGGATGTGG

>Foxc -2132/-1 based on Wagner and Levine 2012

ccgcctacgtagggtaaataccgctcagggcgcgtacgttgcacacagcgggaaatgacaaagaaatggataaaaagatgcatggttttctaaattgtccggcatacagaaaacattcccctgcaaagttatatacgtggtaactcgtaagtgggaatgcggtgttataaaacaaaacacccatgttataacgaccgtcgttttctcggcacttgataataaataaatcgcattcattcaattaataatagatagccagcagcacgcgtatgttatttgagcaatcgcttgcaagaggcacgtccgggaagtccacaccttctacctgaaacgtctactttgacgttatgaacccgtgcaccgaaaagtgttagcgaaaccaagatggttaaactggaatagcttcctcaacacccacgtacactctttccttttgcgtcaagcggggtttcctttccctctagaagtgttgatataaatccttcatgacaccggacaccgcattaagcgcggttcaatcagaatatctatgcggcacccgcttcatgtagtaggccgggggaagatgggacacctttagcacataatacccaaacaacctaatcgtattttaaacagttaacaacggtctatggaagtcgtgaggatacggtttaataattctttaaatatttcttgtttactaccaaatgagacgagaaaatagaataaaaaagtgtcccatctccccccaccctactgtataaaccagaaaaagtggaaatgtccgaaaagagttttatcagcattttagtattttggcgatttaagctttagatacacaaggtgttagtaattgcggaaaggtgttttgttcagttaatctgacgaaaggagttggtttatatttttatttctaccaatatatatatattgctaccaggatttagtaaaaagcgttttatagttaatttaaaagtaaatattttaaacagtaagcaacaaatctgtacgattaattgctccgtaaacgtttactgcatgttcggttaaataaccacctcgcttttcatttttcaatcctactcgttaacgtcgttttgctaattccctttacttttttcaaggccaagttagttactataccaaacgcaatataacataacttaaactgttgtttagctatttaaatacagaaacaaaaataagattaattgaaatagcaaaccaagagaatcagcaacaaaactacacttgttaaaaatacgctatgaaggtaaaaaaaaactaagtaaaaatgtccaatatatttaataaaacgaagctatggtgggtggggaaaccttaagctaaatgctccaggaaaatatgaatcatcgacgcctaagttgccgccttagttgcataactcattgtatagcgagtcacggaaaatgctgcgcgtgtaaatttccgcatggtgtcgctgctgagccaaccggtctcgctcgtttcaaaaagtcgagttttaccgcaaaaaactcttcggttacatctcttatttataaacagcaacccgaggagtcacgctgtaaactgatcgggtcgtgacaaggttcggacaggagaggcagcttcagttataaccgctgaatatcaacggtgaacgttaaccgccatttttaatgaacgttggagttaaaaagttccaagattgagagattaatttaaaagttgtggtttatataaacagggctattgggtaaggctccatagtgagcggtgtcagcaggtgtttcgtaaggcggcgcgtgccaagttctctacttagagcttgtcaaaacacgatctaattactgcatcattagcgcgccattgttcctcgcgaaagttgattgggattatgacgctcctgctttccattgtttaaggggaagatgaactttttaccttcgctcaggctcgactcggtcgtgggcaggtaccggcagaaaacattcgattattgacacgaaggcagtgcgagtgttgtgagggaagtcgtttcggagcgacgtttgtttgcttgcagcgttggcgttcagattctaacttttatatatctcgggcagtgttagtgtaagttaagttacgttgaacacaggacaccgaatccttggtttgattctctata

>Eef1a -1939/-1 based on Sasakura et al. 2010

gtgacgggaaaacgatagtcgttataacacgagtattcatacacctcgtgcgagcttacgagctaccgtatatgttactttgtgggcgaataaaggttttataaatataacattggttttataaataaaacaacgccattttaaagtcggttacataattctgtaactagttcaaattgaacggtaaacgtaaataaaaaccttgaccgtcttacccaattatataaaaacactttgaacgctttttaagatggaagggtatggccatgcctagataattctgtggaccatctcaccccaacctattacagaacggtcgtaataatgaaaatggataccattttaggcatatagactggttcctctactttctagaaacgtaagcagtatacacagaaaaaatgaagtgtgtttctgtgcaaattaaaccgttctaaattcatagccgactgaatttctaattaggtgctgaatggctgacctagatttattgttaagtttagcaccaaatctgagccagcgataagcagtctaattaaattggctgctggcgataaaataggtcatcctgaaaaatcgtttgcgcctttatttaaaatatagtagagtgggggaagacgggacatattttcgttctattttctcgtcccatttggtagtaaacaaagaacattcaaaaaatataaaccataacttcaaaacttcaatagaccgttgtcaactgtttaaaacacaataagagaatttggatattatgtgctaaaggtgtcccatcttcccccaccctactatatctgtttatagttctgtggggtaagataagataccgttaacacctaaacatttttactttaaacaatcaaccacgttttttatagtcgtaatggacatgtggttacataattctgaaaatattttttgcccccgaccaaaagacgcgaagagtaaaaacatgtctcagcttatatttcccacataaatatatttttgtactgtttggtgaatttataaacttatattaccatgcatatacgttatgttactggtattttctcagtaggcaaattcatttgtccacgttttataggtttttaatatttatgatttttaaaatgctaaaaatgcggaaggggggttgaaagtacaatacaaacacacaaaacaatacaaactaaagatttatagttatgctaattcacctacacgatataacaagatgtgtaatgcaaccatgtgtttatgatgagcgctaacatattttgtaaccactcaaattccccgccacacgaggataatgaataggtgactctgtagtctgtacatcttctacgaaataaaataatttctgctcactgattatacttctgttttatagattagaaaacgtttctactaaatgacctaattcgctatacacacacgctgtgcgcgagataatcattctcgcaccccgtttattgtgttaaaattgccgcccagattcacaaagcgtgacggctagagccagcaacgtgtcgccttcaattacgcaacatccgggttgcgcaattctggatataaaagaactaacaaagatgacgtagtacctttttcagttcagacttacgaaagactcacgtgtcggcggtctacttgtccttttcgagctgtggcaatttggtgagtggttctatcttttatctgagtacatctctaaggaattatagtttgattagttaaatttttattgttaggaaagatgaaatcattaggttttacttagtttaagtatgttagtactggttaggcgtttgaattactgaaaaactcagttcgttaactgtagtagttctggtagcttagcaagtataccctgtatacgccttttggctttttaacaataacttaaacttattttacagcaaatttctgtgcattcggttaaccccaaccttccaaa

>FOG -2062/-1 from Rothbächer et al. 2007

gcaactattgtaacaccacacgggggcgcagtagagccagaatcttagtaaatgcggtttgattgtaaagtttttaacaatctctcgctttgtatattctagaatggggaaagatgggacccttaagcactcattgcccaatatttccaaacccaaaatctatgatggttttgtgtggtaccacgaattagtactatcatatacacttcaaagaaatatattaaaccatacgagaagtaatttatctattcgcacaacaggtgtaactttggaaattcgaacacaggcgagctagtttaatgacacgtaaactttatatatttcttttctaatttaaggcatattgaccactgcccatttttcttgttttaatatcccagtttttaaaatagcctgaaatatttcagcttattttattaaattacgatactggattgataattcaaattaacaaactaccaaaaaatgttgacggttttaatataatttaccgagcaccacgtgtaacagcatcatacaaatttaattgattttgtattacttactttataatttaaacatacggtatttagtaaacaaagaacattcaaagaattagaaaaccgtattctcacgactttcatagaccgttgttgattgtttaaaacatgatcataatatgtaggtattatatgtgccaaaggtgtcccgtttccccccacccatactatatatattataaaaaaatggcgttataaaaaaaactataccttgcatgtatttatagatttgttaaacgtatgatctcgtgttatgatctatccgtatagtggaccaagaatctttgggttgtgcatgtttaactgagaaatcgacaagcaattaaacgcggcagtataggtataagccaacacacaatgcgttaaatgcatactatatgttaacgacatgaacaatataattggcgcattataattagattgtataccacgcactacctaaaaagttaagctataactgacgtcgggtattatttattggtatccggcaacactaataaatgaaattttgtcaaaatgtaagccctcgtatctatttaaattttgacgctaaataatccatttatgcagtataataccaaacatacaaaatgtttcagtttgtatcacttttttatgcatcacgtttttgggataaatacacatcaggcatttaacaaaacatgttagatacaatatggcgctgttgctttcacttaaatgtgttaacaattaagatgggaaaaataaaacagtagatatattctttaaatcgaacttttgttgtaaatcttctattttaattatttattcatgtttaaaacgatgtttgtttcggcgtcatgctgctcgataggtataacttcactcgggacattttagtatcgagttgctcaaaattgttagggttgtcaactgaatagcgatgttttcgattgagttaatatttctatacatgatggcatcagctagtaagatagaaaacaaccttgttattactcgtgctgtaataatataattcatgaaaaacaaacatcgtcttatcaaatctattccgaagacaaacatacggctgatagaattcagtcagcgcggtttgggcgacatcactcagagctctagctcttatcttgcttatcgagcaagcgataaaacaagcagtaacgagaaaacaagagaaagaaggctagctttctggagaagaccaagataaagtatctcaaaattcaggaaacggtccaagaccgaagctccaaagctcttgtgttcagttaaactctgatagtgaataagcttcgtgtattgtaccgacccattgtcaatcatgcaaacttgatattatattgacaagagaagaaggcagtttaaattaaaactctaaagtagagagacattaatctcagctgacaaggcaggtggtcacagtaagttcatttaaatagttggccaacaacagcttttccaagaaagtatttttgtttcaggtctatacaaaaataacacacata

>Vsx introns + basal promoter of FOG(-1) exons

from Stolfi and Levine 2011

GGTTCAAATCGGATGAAACCGAACCTGCAGCCAAAAAACGAAAACTTTCTTCGGATGGTACAAAAGgtaattttgttgcattactgctattttcatattctgttattgtggttgcaatatacaatctatagacaacgcccttcattttgtatttttgacataagtgtggccatttaacaaataataccaatagtattttgccgggtaatgctcaaaaactcgtatactaagtatattctggcatgccgcgcagtgaggactcaaccatatatagtatgggagcttgttttgtgagatgtgataccgtgtcaaaatattacggaaatttatttaaagtttaacacaccgcgttatatagtcgttattgtaaatagaagcttataaatcacgtcgtgttacttcttttaaaaaagcaacataaaatattttaaagtcttatcatatagtgaaaaaaaatattccaatattgcttcaattaaaacgtaagcaaattaaacaactaaaaattttattggaaaattccctaaatcaaatataaatttttactgcaggtttctgcattttagattaaatcagcaaatacttacagtcgataaatggcttcaatattgttcatgggattacattatttgaaaaaggggttggcaaaaatttatattttgcatattccatttgctagattttgatccgaatttaatcccgctttgctcgattccatcatgttcacctcattgagcaatgcaccctgcggtaatttgtatcgtttacatttgcgaccgtctggcatatgttccttataccgacgttcacaattgcttgtgttgcgaagatatctttaataacaaattaatcgcagtttcattcgctcaatcggctctcgtggcgaatcaactgctgtcgcttgaaaagaacgcgcatagctccgacaagaggcattatctactcacaataagtgcatcggaaacgattcgcttaattcaccaatcgcttcgctgctttagaataatagaaaacaaaatggcggcgctagcaatgtatcaatgtcatgttgtaacacttacaaatgacagaacgttttcttaatcgaactctctgccttctgtttaggtaataatgaaaacaaaaacactttagtatctactaatacacatgaaaacctttattcgtaaaattaacaaatgattacagttttcaactaaaaaatctttaaaatgaattgaattaaaaaagcaacgcgttttttacagAAGTTACCCAATCTAGAAACAACAGTAAGGAACTCAGTGATAAACTAAAGAAACGACGTAGCAGgtaactttcccaacgtgtggttttaaacaaagtactttctcgacaaatgctctctatttcttattatactcgctatattgatgtattttattctgtatgggtggggcccggcagcgtggcacgccattggtcctaggatttaggccaagattggcccattactctataacccccaaggtgctataccgcatggacactaccagattttggatcctgctacttccccggaggtataagccgccttaatggtttacatattgtatatatttatgcttacgtagatatccatggggtatccatggggtatagaggtcggtagttaaatcaaggatttttaattaatgtactataatttgcggcagAACAATTTTCACAAGCGCTCAACTTGAACATTTGGAAAAAGCATTTGATCTTGCTCATTACCCGGATGCAGAAACGAGAGAAAAATTAGCGAAAAACGCAAATTTACAAGAAGACAGAATTCAGgtaaaaacaaatgtttagtagaatggggtaagatggcctacgtatcattttatttcattgtcccatttagtatttaatagagtgtattaacagaatcttcacaacccttcagagtgttagtgttagttatttaaatcccgttcaggaaatacgagacataattatgcgataacgctacccatcttaccccactgtatagggcctatactatagggcctacttcggtttaaaaaacaacagcagtgttgatngtttaaatttacgaaagccntccaaattaaagcctgtttgtcattttcgttccttggttattgatttgaaataataggccgtaaacgtaaagtcagtcattgaattgattaaagcaatttatctattggttcaatcgctgttggcttcgaattaattatcgaccacacatgtgtggaatcactttttaatctgcgatctacgtacataattatggtaacattggttgtcgcttattcccagtgggtaccaagctcgattaaactgacgcgattaaattagttacaccattataggtgagacctgtacttcttctgggttaactttgtatttctaaatgttggaccaattctattcaaatccattatctattttaacgaacatgatttctttcccttcagGTTTGGTTTCGGAACCGCCGCGCAAAATGGCGGAAACGAGAGAATTGTTGGGGCAAGAGTAGCGTTATGGCAGAATACGGCTTGTACGGTGCAAtggttcgccattctatacctctacccgatactgttattaatgcctgttacaaggacggcctcaaaaacgatgcatttgcaccctggcttattggtaaggcgtaaaccgattatagattactggtgaacgtgtaacacggtgcttaattacgttatattgtattgtaggcatgcataagaaaaccgttcagtcaaaagttactcatgagtcacagcaaagtcaatgctcgaaggaagaatctcgagcagctgaagcttgcatgcctgcaggtcgactctagaggatccggcaaagcttcgtgtattgtaccggcccattgtcaatcatgcaaacttgatattatattgacaagagaagaaggcagtttaaattaaaactctaaagtagagagacattaatctcagctgacaaggcaggtggtcacagtaagttcatttaaatagttggccaacaatagcctttccaagaaagtatttttgttccaggtctatacaaaaataacacacaac

>Pou4 coding sequence START STOP

ATGTTTACTAACATGCTTGCTCCACACCATCACCACCGCCCTCCTTCACCACCTACTACAATGACACCACTCAATACAAGTTACGGTGAACAGAAATACGCCCCGCTTCATAACGCTGCAGATGCCGTCCGTCGACACTGCATGGCATCGCCAGTGGCATATTCCGGCAATCTGTTTGCTGGATTCGATGAGAGTTTTCTTGCTCGCCAGGCCGAAGCGTTGGCCGCTGTAGAATATCCCGGAAAAAGTCAGTTTAAACCCGAACCGACCAACCACAGTTATTACCCCCCAGCCCCCCACTACGTTACGCAACAACACACAACATCAGCCGTACCACCACCAGTTTACAGTTCAATGCACCCTCCACCCGCTTTCCACCACTCAGCGGCGCTGCAAACAAGTCTTGACAGCAGTGACGTCATCGACCAACTTGGGTCTTCTCACTGTGGCCTTCCAACTTATCAACCGCCAACATTTAACTCGTCGGTGATGATGACGTCAGACGTGACGTCACAGCCCCATTACATCACCCACCCCGCCCCAACCTACAACCCCCCGAATCATTTCCATCCTACGTCATCAGCAATGCACTCGACTGTATCAGTCACGCCCATCTCTGCACCAATCATCACCCAAGCCGGCTCCCCTCCGCCAAGCGTGCCAAGTTTTACCTCCAACGAGTACGACGCCGATCCCCGTGAACTAGAAGCCTTCGCAGAAAGATTCAAGCAGCGACGCATTAAACTTGGAGTGACGCAGAATGATGTTGGGCAAGCTCTTGCCAACCTCAAGCTACCGGGTGTTGGATCGCTGAGCCAAAGCACAATTTGCAGGTTTGAGTCCTTGACACTCAGTCACAACAACATGGTCGCCTTGAAACCAATCTTGATCACGTGGTTAGAAAAAGCGGAGGAAGAGCACAGGAGGAAGATGGAGAATGCTGCTATGGGAGAAAAGAAGAGAAAAAGAACTTCCATCGCTGCTCCGGAGAAACGTTCGTTGGAGGCTTACTTCGCCGTGCAGCCCCGACCTTCGTCTGAGAAAATCGCTGCAATCGCGGAAAAGCTCGACTTAAAGAAAAACGTCGTGCGTGTGTGGTTTTGCAATCAAAGGCAGAAACAAAAGAGAATGAAATTTTCAGCATTTAATGGGGACGTGATTATGTAG

>Lhx1/5 coding sequence START STOP

ATGGAAGGACCTTTATGTAGCAGATGTAACCACGTGATCAGCAACAAGTTTGTATTTCAGGTGGACGGGAAATTCTGGCACGGCGATTGCGTCGTTTGTTCCGATTGTGAATGCCCATTGTCTAACCGTTGTTACGTCAGAAATGAAGAATTATTTTGCAGCGACGACTTTTGTCGTCGCTTCGGTAAGAAATGTGGGGGATGCGGGGGTGCATTGTACCCCAACGACCTCGTGCACAAAGTCGGGCAGATAAACTCCGTGTTTCACGTTCGATGTTTCGTATGCTATGTTTGCAACAATAACCTTGAACCTGGGAAGCCTTTTCAAGCTAATGAAAGAGGGCTGCTCTGTAAGGATGATTACGTAGCGATGACGTCATCAGGAGATTCCGAAGAAGAAGTTTGGGGAGAAAGTAATGCACACGTCATCAGTGACGTCACAGGGGTGGCATCGGAAGTGGAAATGACGAAACAATTGATCGACGAGGGTGAGGCTAAACCTGGGGTGTGGGAAAATGAAGGAGGCGAAGTATCAAAGTTGGCGGGGTTTGAGAACCGACATCCTGGCCAAGATGATGAAGATTCCTTGATGTCGCAACAAACGAAAGATTCGAGCGCGTCGACCGCGATTGACACGTCACCACCCCAACAAGGATGCATCGCGATTGCAGCGACGAAGCGTCGTGGACCGCGGACGACGATAAAAGCAAAACAACTGGAAACGCTGAAAAATGCTTTCCTTAGCACCCCTAAACCCACACGTCATATCCGGGAGAAGTTGGCGCAAGATACAGGCCTCTCGATGCGGGTGATTCAGGTTTGGTTCCAGAATCGTCGAAGCAAAGAAAGAAGAATTAAACAATTAAACACGATGGTAGCGCGACGTCAGTACTTTGATCACGTGCCCAATATTGACGTCACAAATGAACAAAGATTTTATGCAGATCCTCACTACTACCCACAACAACAACCCAACTATTACCCAACCGCCGCCAACTTCAACGACAGTAGAATCTCACCAACAGAAAGTGAAACCTGGATTCATAGAATAGAAGACAACCATCATGTGTATGATGACATCATGTATGATGACGTCACTCAAACTCAGAACGGGGGGTTCCCTGGGTTCGCGCCCCGAACCGCGGAAGGCGCGATTTGCGAACAAAGCGGAAAATCTATTGTTTACTGGAATGCAGCGTGA

>Ddpep coding sequence START STOP

ATGACGTCACAAAGAAGTACAACCAGAACATCGGTGGTTGTCTCCGTTCTAGTATTGCTATTCTGGTGTCAGGTGTTCTATACGAGCAAAGCTGATGGCGCTACTGTGGAGCTAAATCGGGAGGCAAAAGTACGAAGCAAACGCCATGTTGCTGACATGCTCATCACTGAGAAGGTATCCAGGGTGAAGCGGGGACATAATAAAAATCTTCTGTTGAGTCGAATCCTACAACTAACTGATGCTTCTACTGGAGGTCACTGTCAATGCATTCTGCAACAAAAAGCAGTAAAGCAAGGCTGCGAATGCAACGAACCAATTCTATATCTTCGCATGATACGGAAGATTACAGATCCCAAAATGAAACTTGGCCATGGATCTGATGCCACTGAGTTGGATCGACTGCTCGCCCGTATGAAGCCAAATATGAAAAGAGCGGTGCTTCACCTAGCGATTAATGAATTTCAGCGACTGCTCCGCGAGGTGGAGGTAGAGCGCACCAGGCGTGGGAGAATCATTCGCCAACTGATAAGACGTCGGCGGAAGTTACGTCATCATCCCCGTAGCGCGTTTGACGGAAAACAATTATTTTCGTTCTAA

>Ddpep::GFP

ATGACGTCACAAAGAAGTACAACCAGAACATCGGTGGTTGTCTCCGTTCTAGTATTGCTATTCTGGTGTCAGGTGTTCTATACGAGCAAAGCTGATGGCGCTACTGTGGAGCTAAATCGGGAGGCAAAAGTACGAAGCAAACGCCATGTTGCTGACATGCTCATCACTGAGAAGGTATCCAGGGTGAAGCGGGGACATAATAAAAATCTTCTGTTGAGTCGAATCCTACAACTAACTGATGCTTCTACTGGAGGTCACTGTCAATGCATTCTGCAACAAAAAGCAGTAAAGCAAGGCTGCGAATGCAACGAACCAATTCTATATCTTCGCATGATACGGAAGATTACAGATCCCAAAATGAAACTTGGCCATGGATCTGATGCCACTGAGTTGGATCGACTGCTCGCCCGTATGAAGCCAAATATGAAAAGAGCGGTGCTTCACCTAGCGATTAATGAATTTCAGCGACTGCTCCGCGAGGTGGAGGTAGAGCGCACCAGGCGTGGGAGAATCATTCGCCAACTGATAAGACGTCGGCGGAAGTTACGTCATCATCCCCGTAGCGCGTTTGACGGAAAACAATTATTTTCGTTCACTAGTACCATGGTGAGCAAGGGCGAGGAGCTGTTCACCGGGGTGGTGCCCATCCTGGTCGAGCTGGACGGCGACGTAAACGGCCACAAGTTCAGCGTGTCCGGCGAGGGCGAGGGCGATGCCACCTACGGCAAGCTGACCCTGAAGTTCATCTGCACCACCGGCAAGCTGCCCGTGCCCTGGCCCACCCTCGTGACCACCCTGACCTACGGCGTGCAGTGCTTCAGCCGCTACCCCGACCACATGAAGCAGCACGACTTCTTCAAGTCCGCCATGCCCGAAGGCTACGTCCAGGAGCGCACCATCTTCTTCAAGGACGACGGCAACTACAAGACCCGCGCCGAGGTGAAGTTCGAGGGCGACACCCTGGTGAACCGCATCGAGCTGAAGGGCATCGACTTCAAGGAGGACGGCAACATCCTGGGGCACAAGCTGGAGTACAACTACAACAGCCACAACGTCTATATCATGGCCGACAAGCAGAAGAACGGCATCAAGGTGAACTTCAAGATCCGCCACAACATCGAGGACGGCAGCGTGCAGCTCGCCGACCACTACCAGCAGAACACCCCCATCGGCGACGGCCCCGTGCTGCTGCCCGACAACCACTACCTGAGCACCCAGTCCGCCCTGAGCAAAGACCCCAACGAGAAGCGCGATCACATGGTCCTGCTGGAGTTCGTGACCGCCGCCGGGATCACTCTCGGCATGGACGAGCTGTACAAGTAA

>GFP::Nckap5

ATGGTGAGCAAGGGCGAGGAGCTGTTCACCGGGGTGGTGCCCATCCTGGTCGAGCTGGACGGCGACGTAAACGGCCACAAGTTCAGCGTGTCCGGCGAGGGCGAGGGCGATGCCACCTACGGCAAGCTGACCCTGAAGTTCATCTGCACCACCGGCAAGCTGCCCGTGCCCTGGCCCACCCTCGTGACCACCCTGACCTACGGCGTGCAGTGCTTCAGCCGCTACCCCGACCACATGAAGCAGCACGACTTCTTCAAGTCCGCCATGCCCGAAGGCTACGTCCAGGAGCGCACCATCTTCTTCAAGGACGACGGCAACTACAAGACCCGCGCCGAGGTGAAGTTCGAGGGCGACACCCTGGTGAACCGCATCGAGCTGAAGGGCATCGACTTCAAGGAGGACGGCAACATCCTGGGGCACAAGCTGGAGTACAACTACAACAGCCACAACGTCTATATCATGGCCGACAAGCAGAAGAACGGCATCAAGGTGAACTTCAAGATCCGCCACAACATCGAGGACGGCAGCGTGCAGCTCGCCGACCACTACCAGCAGAACACCCCCATCGGCGACGGCCCCGTGCTGCTGCCCGACAACCACTACCTGAGCACCCAGTCCGCCCTGAGCAAAGACCCCAACGAGAAGCGCGATCACATGGTCCTGCTGGAGTTCGTGACCGCCGCCGGGATCACTCTCGGCATGGACGAGCTGTACAAgccggcggccgcaaccATGGAATTAAAGTCTGAGTTTGACAACATGGAAGAATCTCTTCGTGACGCTTTACGTCATACAGAACATCACACAGATCTTGGCCGCTATGTAACTTTACAAGAACAGTCTCAACCATTGAAACCATCTATATATGTTGGTGCAACTGAGCAAAAGAGGAATATGGAATTACAAAACTTCACTTCAACATCTAATAACAATCAAATGCATTGTCAGAGAGATATGCGGCAAGCAGTCCTGGACCTTGATGATTTTCGAAGACACTGCATTTTGTTAGCTGAAAGAGTTCAAAAAATGGAAGAACAAAATTATCAGCTTATTAAACTAGCTGAAAAACAGAAAGTGGAATATAATAAGGTTGCAGCTCAGGTGGTCCAATCGGAACTCATTCAACGTCAGTTGCAATCTGAGTGCTTCAATTTAAAACGACAAGTTGACAATCTTTCCAAATATAACAAAATATTGCAACAACAAAATGAAACTTCTTTTTCCGGAGTTGAAAAAGAACATGGTAAAAACAATAGACAAGTTTTGATAGAAGATTCAAACGACCCAATGAGAGATTTGCACAGCAGCGTTGAGAGAAAACCCAAAAATGGGTTAACAGATAAGCTTTTCACCAAAGCGATGGGACGTCTTCAGTCTTCACCTAACACTGAAGATTCTGATCCTTACAAAGCAACTGATGTTGCTACAGTCGTAGAACTATATTCTGCTTTACAGAAGTCAACAGTTTCCGATGTTAATTCGGAACCAATAAAGACACCTAATATCCCAAACAGTGATAATGCATCAGTTGGTTCACTTAATATAATACATGAGCATCCTGAACCTTATACAACGAACGAACATGCTGATTACGCCACACCGTCCAGCAAATGTGCATTACAGTCTTGCTCATCTTGCCCCCAAAGACATTCCTCCTCCTCCAAGCCTCCACTGAGTCCTTCAAGATCATGTTCGGGTGAATCCCTGGAACAGAGAAAAGAATCGTGCAAAAAAACGAACAACCGAAAACTTTTTAAAATTAAAAAAAGCACAAGCGGAAGTAACTTAGAAAATAGTCAGTCTTTCGACAATAATTCCTACATTGAAGTTGAGCTCAGAAGATACAAAGAGGATTCGACTAAAAATAACGATCGATTGAATCCTGCATCTCAAAACAAAAACTCTCATAGTAAACGAACCACACAAAGACCGTCGAGTTTGGTTTTACAATCCGATTCTTCAGACACGTTGAACCAAACCTCAGCTTGTAGTGACTCTCAACCCTCGGCTTCATTCTATGAAAATGAGCCGTTTAACCAGAATTCTAAAAATACCAATAACAACAATGGATCTCAGCATCAGCAGAAAACACAAGCTCACAGAGAAATCTCATTGCGTTTAGAAGAAAAGTCGCTTTCACCTGATTCTCTCAGTTCTAGATCATCCTCGTGGAACAAATCTTCTCCAGGTTTAATAAGAAGATCACATTCCAAAACACCATTATTTACAGATCTTCCACCTCCACCTCCTCCCCCTCCATGCGAGGATCAAACGGAAGATGGAGCAAACAGGAGATCCAAAGAATTTTCTCGTTCTGAGTCGTTAAACAAAGATGACAGTGTTTTTTCCACCGACACGCAACCAAGAAATCAAACCGAAGGAAGCAGACCTTTACCTAAACTCAGTGTATGGAAGGAATCGGCTGACAATGAATCAGGAAAATCACCAAACCAGGACACCAGTGGGTGGAGAAGGAAGTCATATTCAGCTGCTTGCAGAAAGTCACAGTCGTTGGATCATATATACGAACTTGAACAATTTGATAATAACAAGAAAAATATGGAAGAAAAGTTGGACGTCTGCTCCATGGTATTCAGCACTCTACAAACTCCCAAAAGATGCTCAAAGCCGCAACCTTCTTTTTCTCCACAATCTCCAGATGCTTTTGTAGAAAAATCGAAAGAACAATCTCTGGAAGATGAGTTTCCTCCTCCCCCACCTCCCCACATCCTCCAGGAGGTGATGAAAGGATTTGCTCCCATTGAAGAGGATGGTTATGTTCTCATGGACCCCATCGAGAATGATAAAAACGCCAAATTTAAAGATCTAAGTCCAAAGCAAAATTCTCAGTTTGAATTAACAAAGAAGAACAGCAAAAGTGAATCGTCAATTCACCAACGCAGGTCTTTATCAAAAGATCATTCAAAATCTTCTGACATAGGTGGGAGACCAAGTAAAAACGAAGAGGAAAGTTTCAAACCTGAAATTGGTCGACCTAAAAACCGAAGAATGAGTGAAGGCTCCGCTGTGAGACGAAGATCCGACCGACAGTCCCTTGAAGACGATGGTGGTATGAGAAAAATTTCTCGTCATCGGAGCATTAGCAGTGACCACTCGTTAAGAAAGCGAGAAAACTATGTCCCTGAAAACCAAAATGATTTTACGAAAGCAAATCTCCAGCCCCGATTTCCATCAACCAACCCCTATGTTAAGTTAACTTCAACTCCTAAAGCAAGATTATCAAATGCAGAAGAGGAAAAATCCCCCATGCCTAGAAGTGTAAAAGAGTCAAAACCTCTGCAAGGACGACAACCCACAAAGCTTACACCCAACGTAAAGGTAACGACAAATTCTGCAACAAACGGAAAGAAGAAGTCAAGTTCTCCATCATCATTTGTGAATCTTATGAAAGCAAAGTTTGGAAAATCAAATTTAGGAAACAAATCTGGAAGTAAGAAGTCTGAAGAAGAAAATGTTTTAGAGAAAACCGAATCTTCCCTTCCGGTTAAAGCTGCAAGCTCCTTGATTGATCCGCGTACTCTTTCGTTAGAACCAAACTCATATTTAAAACCAACCGTTGATGGTGTACTTGATGATAAAAGATCCAAACTTAAACTTGGAAACAACAGCGACGCAGCTGTAGGAAGAAAACTAAGTCGTCCCTTTCCAAATCAACCACTTGCGAATCAACAATCAAAGGGACAATTATTGAACGTGGTTGAAGAAGTGGTTTCCGATGAAGGATTTGCTGAAGGAAGCACAATTGAACGTGCTGAAAAAGAACAAACTTCCGAAAAAGACGTGAATGAAACAAACTGTGAGTCACGATATGTAGATATCACAGTTCAAAAACAAGAACAGCGTAATTGCTTAAACAAAAATCCCGCTGAGATTGGGATCAAAAATCAACCTAAAAAAAACAATTCTTATTGGTATGACAATGTAGTTAGCATAGGAAGCGGTAGTAACAAACAATTTAAAGTTTTAGTGACCGAAACAACTTTTGATGAAGATGGAATAACAAGAAGCAACTGTGTGAAGCAGAAACCCACAAGGTTAAGAATGAAGGAAGAACCTCCATATGAAAATTACCCATTAGTAACTAGTTCAATGGGCCAGCTAGCCCCACCGTCATACGAACATGTTATTCTCACGAGGTCACAAAAAAGTGACAACCCCAGACCGACATCTGTGAGTTACAACTCCCAAAACCAAAATCGTGAAGTGATAGCCACAGGCGTGACTATTGTGCGGTCACCTCAACAAAAAGCTTTCAAACCAAAAGTTGAAAGTAAACCGACCAAGACTACTACAGCGTTTGACAGGATAATGAAATCTGTTGGTCATCATTATTCAGGAAGTAAAGAAGACCTGAGTGGGAAAAGAGAATTAGATTTAAAAAATACTGCCCAAAAATCACACAGAAATACAGGCAAGAACGAAAGTCCACAAGGAAAACTTAATGCTGAACATCAAAAAGTGTTTGAAAAAGCTTTTATTAGCTCGGTCGTTGAAGCGGAACTTTCACAAACATCTCATAACTTGATGGAAGGTGGAAATGATCAGAAAACATCGAGTGAGTTGCCATGTCTTCCTCTACCCCCATCGAGTGGTGGAAACAGCACTGGAACAGGTTCCAGTTTCGGATCTCTACCTTCTGATGGCATTGCGGACGTCACAGAGTCTCGTACTAGGACGTCTTCTGCGTCTTGCCCCTCACTAAAGTCGCAATTGAGATTAAGCAAGACACAAAGGAACAGTATTACTTCGTTTGGTTATTCCGCTCTTCAAGGGACAGTGGATTCCAGCACATCGACGACTAGTTCTGAATCGAGTTCAAGAGGTCGCGATAAAAATTCAGTTGGAGGATGTTTGGAGTCGAACTCATCGTTTGGGTCAGTGGAGCGACCAGTGACCTCATCCGGGATTACGAAATCAAAAATCAACCCCATGACACAATCACGACTTTCAAAATCACCAAAAAATCCGAGTTTTGAACAAGAAACGCAAACAAAACTTGCTCCAAACCGTAGATCTCAGTCAGGATCACGAATATCTCGACTTAGCACGCCAAGCAAACTGCCCGCACCAGGAAAATCTCCTGACTCAAGATCTACCACTCCTAAAAGTGTGTTTTCACCTTCCACTCCAAATGAATTACCTCCTACATCAGCAAACTCCAAGCAAAGATTAAAGCTGGCTCCTCTAGGCTTATTACCCCCTAGCCCATCTTACACCAATCAAACCACTGAACCAGTTTATGAAAATGCTGCTGCCTTTAAACAACACAGAGTTAATGAAAAGCAAATACCCGCCCTCAAAAGGATCCCATCGTCCATGAGGCCATCTGGTTCCAGCATCGGCATGCCATCTGTTTCAGCGCGAACAAATAGTTGTGCCTCCAATCATATAGACACCCGACGTTCGAGGAAAGGTAACCCATCAGCATCTGGACTGCATCGCCCCCGTCCAATCTACCCAGGAGTTGGAGCTGGTTAA

>nls::Cas9::nls from Stolfi et al. 2014

ATGGCTAGCCCCAAAAAGAAGAGGAAAGTGGACAAGAAGTATTCTATCGGACTGGACATCGGGACTAATAGCGTCGGGTGGGCCGTGATCACTGACGAGTACAAGGTGCCCTCTAAGAAGTTCAAGGTGCTCGGGAACACCGACCGGCATTCCATCAAGAAAAATCTGATCGGAGCTCTCCTCTTTGATTCAGGGGAGACCGCTGAAGCAACCCGCCTCAAGCGGACTGCTAGACGGCGGTACACCAGGAGGAAGAACCGGATTTGTTACCTTCAAGAGATATTCTCCAACGAAATGGCAAAGGTCGACGACAGCTTCTTCCATAGGCTGGAAGAATCATTCCTCGTGGAAGAGGATAAGAAGCATGAACGGCATCCCATCTTCGGTAATATCGTCGACGAGGTGGCCTATCACGAGAAATACCCAACCATCTACCATCTTCGCAAAAAGCTGGTGGACTCAACCGACAAGGCAGACCTCCGGCTTATCTACCTGGCCCTGGCCCACATGATCAAGTTCAGAGGCCACTTCCTGATCGAGGGCGACCTCAATCCTGACAATAGCGATGTGGATAAACTGTTCATCCAGCTGGTGCAGACTTACAACCAGCTCTTTGAAGAGAACCCCATCAATGCAAGCGGAGTCGATGCCAAGGCCATTCTGTCAGCCCGGCTGTCAAAGAGCCGCGGACTTGAGAATCTTATCGCTCAGCTGCCGGGTGAAAAGAAAAATGGACTGTTCGGGAACCTGATTGCTCTTTCACTTGGGCTGACTCCCAATTTCAAGTCTAATTTCGACCTGGCAGAGGATGCCAAGCTGCAACTGTCCAAGGACACCTATGATGACGATCTCGACAACCTCCTGGCCCAGATCGGTGACCAATACGCCGACCTTTTCCTTGCTGCTAAGAATCTTTCTGACGCCATCCTGCTGTCTGACATTCTCCGCGTGAACACTGAAATCACCAAGGCCCCTCTTTCAGCTTCAATGATTAAGCGGTATGATGAGCACCACCAGGACCTGACCCTGCTTAAGGCACTCGTCCGGCAGCAGCTTCCGGAGAAGTACAAGGAAATCTTCTTTGACCAGTCAAAGAATGGATACGCCGGCTACATCGACGGAGGTGCCTCCCAAGAGGAATTTTATAAGTTTATCAAACCTATCCTTGAGAAGATGGACGGCACCGAAGAGCTCCTCGTGAAACTGAATCGGGAGGATCTGCTGCGGAAGCAGCGCACTTTCGACAATGGGAGCATTCCCCACCAGATCCATCTTGGGGAGCTTCACGCCATCCTTCGGCGCCAAGAGGACTTCTACCCCTTTCTTAAGGACAACAGGGAGAAGATTGAGAAAATTCTCACTTTCCGCATCCCCTACTACGTGGGACCCCTCGCCAGAGGAAATAGCCGGTTTGCTTGGATGACCAGAAAGTCAGAAGAAACTATCACTCCCTGGAACTTCGAAGAGGTGGTGGACAAGGGAGCCAGCGCTCAGTCATTCATCGAACGGATGACTAACTTCGATAAGAACCTCCCCAATGAGAAGGTCCTGCCGAAACATTCCCTGCTCTACGAGTACTTTACCGTGTACAACGAGCTGACCAAGGTGAAATATGTCACCGAAGGGATGAGGAAGCCCGCATTCCTGTCAGGCGAACAAAAGAAGGCAATTGTGGACCTTCTGTTCAAGACCAATAGAAAGGTGACCGTGAAGCAGCTGAAGGAGGACTATTTCAAGAAAATTGAATGCTTCGACTCTGTGGAGATTAGCGGGGTCGAAGATCGGTTCAACGCAAGCCTGGGTACCTACCATGATCTGCTTAAGATCATCAAGGACAAGGATTTTCTGGACAATGAGGAGAACGAGGACATCCTTGAGGACATTGTCCTGACTCTCACTCTGTTCGAGGACCGGGAAATGATCGAGGAGAGGCTTAAGACCTACGCCCATCTGTTCGACGATAAAGTGATGAAGCAACTTAAACGGAGAAGATATACCGGATGGGGACGCCTTAGCCGCAAACTCATCAACGGAATCCGGGACAAACAGAGCGGAAAGACCATTCTTGATTTCCTTAAGAGCGACGGATTCGCTAATCGCAACTTCATGCAACTTATCCATGATGATTCCCTGACCTTTAAGGAGGACATCCAGAAGGCCCAAGTGTCTGGACAAGGTGACTCACTGCACGAGCATATCGCAAATCTGGCTGGTTCACCCGCTATTAAGAAGGGTATTCTCCAGACCGTGAAAGTCGTGGACGAGCTGGTCAAGGTGATGGGTCGCCATAAACCAGAGAACATTGTCATCGAGATGGCCAGGGAAAACCAGACTACCCAGAAGGGACAGAAGAACAGCAGGGAGCGGATGAAAAGAATTGAGGAAGGGATTAAGGAGCTCGGGTCACAGATCCTTAAAGAGCACCCGGTGGAAAACACCCAGCTTCAGAATGAGAAGCTCTATCTGTACTACCTTCAAAATGGACGCGATATGTATGTGGACCAAGAGCTTGATATCAACAGGCTCTCAGACTACGACGTGGACCACATCGTCCCTCAGAGCTTCCTCAAAGACGACTCAATTGACAATAAGGTGCTGACTCGCTCAGACAAGAACCGGGGAAAGTCAGATAACGTGCCCTCAGAGGAAGTCGTGAAAAAGATGAAGAACTATTGGCGCCAGCTTCTGAACGCAAAGCTGATCACTCAGCGGAAGTTCGACAATCTCACTAAGGCTGAGAGGGGCGGACTGAGCGAACTGGACAAAGCAGGATTCATTAAACGGCAACTTGTGGAGACTCGGCAGATTACTAAACATGTCGCCCAAATCCTTGACTCACGCATGAATACCAAGTACGACGAAAACGACAAACTTATCCGCGAGGTGAAGGTGATTACCCTGAAGTCCAAGCTGGTCAGCGATTTCAGAAAGGACTTTCAATTCTACAAAGTGCGGGAGATCAATAACTATCATCATGCTCATGACGCATATCTGAATGCCGTGGTGGGAACCGCCCTGATCAAGAAGTACCCAAAGCTGGAAAGCGAGTTCGTGTACGGAGACTACAAGGTCTACGACGTGCGCAAGATGATTGCCAAATCTGAGCAGGAGATCGGAAAGGCCACCGCAAAGTACTTCTTCTACAGCAACATCATGAATTTCTTCAAGACCGAAATCACCCTTGCAAACGGTGAGATCCGGAAGAGGCCGCTCATCGAGACTAATGGGGAGACTGGCGAAATCGTGTGGGACAAGGGCAGAGATTTCGCTACCGTGCGCAAAGTGCTTTCTATGCCTCAAGTGAACATCGTGAAGAAAACCGAGGTGCAAACCGGAGGCTTTTCTAAGGAATCAATCCTCCCCAAGCGCAACTCCGACAAGCTCATTGCAAGGAAGAAGGATTGGGACCCTAAGAAGTACGGCGGATTCGATTCACCAACTGTGGCTTATTCTGTCCTGGTCGTGGCTAAGGTGGAAAAAGGAAAGTCTAAGAAGCTCAAGAGCGTGAAGGAACTGCTGGGTATCACCATTATGGAGCGCAGCTCCTTCGAGAAGAACCCAATTGACTTTCTCGAAGCCAAAGGTTACAAGGAAGTCAAGAAGGACCTTATCATCAAGCTCCCAAAGTATAGCCTGTTCGAACTGGAGAATGGGCGGAAGCGGATGCTCGCCTCCGCTGGCGAACTTCAGAAGGGTAATGAGCTGGCTCTCCCCTCCAAGTACGTGAATTTCCTCTACCTTGCAAGCCATTACGAGAAGCTGAAGGGGAGCCCCGAGGACAACGAGCAAAAGCAACTGTTTGTGGAGCAGCATAAGCATTATCTGGACGAGATCATTGAGCAGATTTCCGAGTTTTCTAAACGCGTCATTCTCGCTGATGCCAACCTCGATAAAGTCCTTAGCGCATACAATAAGCACAGAGACAAACCAATTCGGGAGCAGGCTGAGAATATCATCCACCTGTTCACCCTCACCAATCTTGGTGCCCCTGCCGCATTCAAGTACTTCGACACCACCATCGACCGGAAACGCTATACCTCCACCAAAGAAGTGCTGGACGCCACCCTCATCCACCAGAGCATCACCGGACTTTACGAAACTCGGATTGACCTCTCACAGCTCGGAGGGGATGAGGGAGCTCCCAAGAAAAAGCGCAAGGTAGGTTAA

>nls::Cas9::nls::CionaGeminin-Nterminus (from Song et al. 2022)

nls::Cas9::nls (described in Stolfi et al. 2014)

Ciona robusta Geminin N-terminus

ATGGCTAGCCCCAAAAAGAAGAGGAAAGTGGACAAGAAGTATTCTATCGGACTGGACATCGGGACTAATAGCGTCGGGTGGGCCGTGATCACTGACGAGTACAAGGTGCCCTCTAAGAAGTTCAAGGTGCTCGGGAACACCGACCGGCATTCCATCAAGAAAAATCTGATCGGAGCTCTCCTCTTTGATTCAGGGGAGACCGCTGAAGCAACCCGCCTCAAGCGGACTGCTAGACGGCGGTACACCAGGAGGAAGAACCGGATTTGTTACCTTCAAGAGATATTCTCCAACGAAATGGCAAAGGTCGACGACAGCTTCTTCCATAGGCTGGAAGAATCATTCCTCGTGGAAGAGGATAAGAAGCATGAACGGCATCCCATCTTCGGTAATATCGTCGACGAGGTGGCCTATCACGAGAAATACCCAACCATCTACCATCTTCGCAAAAAGCTGGTGGACTCAACCGACAAGGCAGACCTCCGGCTTATCTACCTGGCCCTGGCCCACATGATCAAGTTCAGAGGCCACTTCCTGATCGAGGGCGACCTCAATCCTGACAATAGCGATGTGGATAAACTGTTCATCCAGCTGGTGCAGACTTACAACCAGCTCTTTGAAGAGAACCCCATCAATGCAAGCGGAGTCGATGCCAAGGCCATTCTGTCAGCCCGGCTGTCAAAGAGCCGCAGACTTGAGAATCTTATCGCTCAGCTGCCGGGTGAAAAGAAAAATGGACTGTTCGGGAACCTGATTGCTCTTTCACTTGGGCTGACTCCCAATTTCAAGTCTAATTTCGACCTGGCAGAGGATGCCAAGCTGCAACTGTCCAAGGACACCTATGATGACGATCTCGACAACCTCCTGGCCCAGATCGGTGACCAATACGCCGACCTTTTCCTTGCTGCTAAGAATCTTTCTGACGCCATCCTGCTGTCTGACATTCTCCGCGTGAACACTGAAATCACCAAGGCCCCTCTTTCAGCTTCAATGATTAAGCGGTATGATGAGCACCACCAGGACCTGACCCTGCTTAAGGCACTCGTCCGGCAGCAGCTTCCGGAGAAGTACAAGGAAATCTTCTTTGACCAGTCAAAGAATGGATACGCCGGCTACATCGACGGAGGTGCCTCCCAAGAGGAATTTTATAAGTTTATCAAACCTATCCTTGAGAAGATGGACGGCACCGAAGAGCTCCTCGTGAAACTGAATCGGGAGGATCTGCTGCGGAAGCAGCGCACTTTCGACAATGGGAGCATTCCCCACCAGATCCATCTTGGGGAGCTTCACGCCATCCTTCGGCGCCAAGAGGACTTCTACCCCTTTCTTAAGGACAACAGGGAGAAGATTGAGAAAATTCTCACTTTCCGCATCCCCTACTACGTGGGACCCCTCGCCAGAGGAAATAGCCGGTTTGCTTGGATGACCAGAAAGTCAGAAGAAACTATCACTCCCTGGAACTTCGAAGAGGTGGTGGACAAGGGAGCCAGCGCTCAGTCATTCATCGAACGGATGACTAACTTCGATAAGAACCTCCCCAATGAGAAGGTCCTGCCGAAACATTCCCTGCTCTACGAGTACTTTACCGTGTACAACGAGCTGACCAAGGTGAAATATGTCACCGAAGGGATGAGGAAGCCCGCATTCCTGTCAGGCGAACAAAAGAAGGCAATTGTGGACCTTCTGTTCAAGACCAATAGAAAGGTGACCGTGAAGCAGCTGAAGGAGGACTATTTCAAGAAAATTGAATGCTTCGACTCTGTGGAGATTAGCGGGGTCGAAGATCGGTTCAACGCAAGCCTGGGTACCTACCATGATCTGCTTAAGATCATCAAGGACAAGGATTTTCTGGACAATGAGGAGAACGAGGACATCCTTGAGGACATTGTCCTGACTCTCACTCTGTTCGAGGACCGGGAAATGATCGAGGAGAGGCTTAAGACCTACGCCCATCTGTTCGACGATAAAGTGATGAAGCAACTTAAACGGAGAAGATATACCGGATGGGGACGCCTTAGCCGCAAACTCATCAACGGAATCCGGGACAAACAGAGCGGAAAGACCATTCTTGATTTCCTTAAGAGCGACGGATTCGCTAATCGCAACTTCATGCAACTTATCCATGATGATTCCCTGACCTTTAAGGAGGACATCCAGAAGGCCCAAGTGTCTGGACAAGGTGACTCACTGCACGAGCATATCGCAAATCTGGCTGGTTCACCCGCTATTAAGAAGGGTATTCTCCAGACCGTGAAAGTCGTGGACGAGCTGGTCAAGGTGATGGGTCGCCATAAACCAGAGAACATTGTCATCGAGATGGCCAGGGAAAACCAGACTACCCAGAAGGGACAGAAGAACAGCAGGGAGCGGATGAAAAGAATTGAGGAAGGGATTAAGGAGCTCGGGTCACAGATCCTTAAAGAGCACCCGGTGGAAAACACCCAGCTTCAGAATGAGAAGCTCTATCTGTACTACCTTCAAAATGGACGCGATATGTATGTGGACCAAGAGCTTGATATCAACAGGCTCTCAGACTACGACGTGGACCACATCGTCCCTCAGAGCTTCCTCAAAGACGACTCAATTGACAATAAGGTGCTGACTCGCTCAGACAAGAACCGGGGAAAGTCAGATAACGTGCCCTCAGAGGAAGTCGTGAAAAAGATGAAGAACTATTGGCGCCAGCTTCTGAACGCAAAGCTGATCACTCAGCGGAAGTTCGACAATCTCACTAAGGCTGAGAGGGGCGGACTGAGCGAACTGGACAAAGCAGGATTCATTAAACGGCAACTTGTGGAGACTCGGCAGATTACTAAACATGTCGCCCAAATCCTTGACTCACGCATGAATACCAAGTACGACGAAAACGACAAACTTATCCGCGAGGTGAAGGTGATTACCCTGAAGTCCAAGCTGGTCAGCGATTTCAGAAAGGACTTTCAATTCTACAAAGTGCGGGAGATCAATAACTATCATCATGCTCATGACGCATATCTGAATGCCGTGGTGGGAACCGCCCTGATCAAGAAGTACCCAAAGCTGGAAAGCGAGTTCGTGTACGGAGACTACAAGGTCTACGACGTGCGCAAGATGATTGCCAAATCTGAGCAGGAGATCGGAAAGGCCACCGCAAAGTACTTCTTCTACAGCAACATCATGAATTTCTTCAAGACCGAAATCACCCTTGCAAACGGTGAGATCCGGAAGAGGCCGCTCATCGAGACTAATGGGGAGACTGGCGAAATCGTGTGGGACAAGGGCAGAGATTTCGCTACCGTGCGCAAAGTGCTTTCTATGCCTCAAGTGAACATCGTGAAGAAAACCGAGGTGCAAACCGGAGGCTTTTCTAAGGAATCAATCCTCCCCAAGCGCAACTCCGACAAGCTCATTGCAAGGAAGAAGGATTGGGACCCTAAGAAGTACGGCGGATTCGATTCACCAACTGTGGCTTATTCTGTCCTGGTCGTGGCTAAGGTGGAAAAAGGAAAGTCTAAGAAGCTCAAGAGCGTGAAGGAACTGCTGGGTATCACCATTATGGAGCGCAGCTCCTTCGAGAAGAACCCAATTGACTTTCTCGAAGCCAAAGGTTACAAGGAAGTCAAGAAGGACCTTATCATCAAGCTCCCAAAGTATAGCCTGTTCGAACTGGAGAATGGGCGGAAGCGGATGCTCGCCTCCGCTGGCGAACTTCAGAAGGGTAATGAGCTGGCTCTCCCCTCCAAGTACGTGAATTTCCTCTACCTTGCAAGCCATTACGAGAAGCTGAAGGGGAGCCCCGAGGACAACGAGCAAAAGCAACTGTTTGTGGAGCAGCATAAGCATTATCTGGACGAGATCATTGAGCAGATTTCCGAGTTTTCTAAACGCGTCATTCTCGCTGATGCCAACCTCGATAAAGTCCTTAGCGCATACAATAAGCACAGAGACAAACCAATTCGGGAGCAGGCTGAGAATATCATCCACCTGTTCACCCTCACCAATCTTGGTGCCCCTGCCGCATTCAAGTACTTCGACACCACCATCGACCGGAAACGCTATACCTCCACCAAAGAAGTGCTGGACGCCACCCTCATCCACCAGAGCATCACCGGACTTTACGAAACTCGGATTGACCTCTCACAGCTCGGAGGGGATGAGGGAGCTCCCAAGAAAAAGCGCAAGGTAATGGCCACGAAAAATATTCTTCAAAATATAAATGCACAATGGAAGGAGAATGACAACAGATCACCAAGTAGAAAGCGACGGTTAGATGACGTCACTGAAGAATCACAATTACCTTCCACGACCAAACGACGTCATCTTCAAACAAATACAAACGTTGTAAATTCCACAGGATTGAAACAAGGCCTGACAAATGTGAAAAATTCAATAAATCCAAAGAACAAATCAATAAAAAATTTCTTTTCTGATATTCCACGTGTGTCATGTACTAAATCTGAAAAGATTCAAATTTTTAAAGAAGCTAAGAAAACTCCAAAAAAGAATGCAACCACTCAGACAAGGAGTGAAGCTGAAGAATTGGTCTGCAGTGATCAACCCAGTGAAAAATATTGGGAACTCTTAGCCGAGGAGCGAAGGAAAGGGTTGTAA

>Styela clava Ddpep (NCBI Reference Sequence: XP_039257153.1)

MTQKVFKFNAYWIIAGFLFGILLHVGPSKEAKLEKSAIKQVLKNYEKVVDAQSSLTDASSEVLFRGKRHT

ADHLMNDKYSRTLKSHARKQILSTLLKLVGSTAGGPCQCESHSHLDDQTSRLKSRQDAISKRVHTEPSFD

CNCDMYSLVSYLAERLSQGGEDEESTGSDPLSGYLSRLSPSVKARLVHFLNEESMRIKRNEMLRKLRRMR

LHRQFLHRRRLLRNDPNAAFRGKQFLSF

>Halocynthia roretzi Ddpep (Harore.g00004091)

MWQTRLGSNLVYVALLCFLLCKPVMVASKRQVNAKPQKGDLQNGIQQFHTDSQATIIADEAGKTTEFSLGESGRSRQKRHTADHLISDKYSRSLKSQARKQILSKLLNLVGSSPGNPCECLFAQSTSDKDSSSRLIQRDVFKRLDSISSSSFKCNCDISSLIKFILSKLKFTKNSNTDPLGRYMSMLSEATKTRLAHFLHEEELRLKKNEILRKLRIRRMHRMMMHRRRLLHKDPRAAFRGKQFFSF

>Halocynthia aurantium Ddpep (Haaura.g00003261)

MWQTRLSSNLVYVALLCFLLRKPVMVASKHEVSAKPQKDDLQNGIQQFHTDSQATIINDETGKTAEFSRGESGRSREKRHTADHLISDKYSRSLKSQARKQILSKLLNLVGSSPDNPCECLFAQSTSDKDSSSRLIERDVFKRLDSISSSSFKCNCDISSLIKFILSKLKFTKKSNTDPLGRYMSMLSEATKTRLAHFLHEEELRLKKNEILRKLRIRRMHRMMMHRRRLLHKDPRAAFRGKQFFSF

>Ciona robusta Ddpep (Cirobu.g00002781

MTSQRSTTRTSVVVSVLVLLFWCQVFYTSKADGATVELNREAKVRSKRHVADMLITEKVSRVKRGHNKNLLLSRILQLTDASTGGHCQCILQQKAVKQGCECNEPILYLRMIRKITDPKMKLGHGSDATELDRLLARMKPNMKRAVLHLAINEFQRLLREVEVERTRRGRIIRQLIRRRRKLRHHPRSAFDGKQLFSF

>Ciona robusta Seldom (Cirobu.g00006902)

MRRRRISSTNLQSAVDVDDYRRKPAVRCGNLVKLGNNVFRLQDLPEDSWLCRSLEIIGLFVVVLLVYVTYFHYESLHFHVAKGYGHLGYAPAQHVVGQRYLTGRGAPKNETEAMKWFKYAADQGHAEASFNLAVGHIHGIKTGLRPGQPTRLLHHAKKNGVEEAEHALSLCARRGCDM

>Styela clava Seldom (NCBI Reference Sequence: XP_039249433.1)

MELVVNISVSLLLVLLAERRPRSALDSRHYKTTMTGKTLRHRKSSNKIVEEQAGDEYAADATSSHNDMER

YVKLGHSVYRMQGSKRDPMWVTALEVFGVLLAFSLIYTSYYYYDHLHFHVSKGYAHLGYSSAQHVVGQRY

LAGKGTDKNDTLAMQWFRAASDQGHAEASYNLAVGHMHGSQTNLKRGEPEKLLRFAADKGVKAAHHALNL

CARRGCD

>Molgula oculata Seldom (Moocul.g00007273)
MSIRRRRSEQKLLSKEGRNDDEPSYEETPLANRKWVRIGKNVFRVSRPHQDSWWCTMMEVLAVTTSLLLMYGILYHYEMMHYHVSKGYAQLGYPSAQHVVGQRLLAGKGVQQNDSKAMEWFRYAADQGHPEAGYHLAIGQMQGVHSMLKRGESERLLRHAADNGVSAAQHALDLCPKRGCD

>Choloepus didactylus (Southern two-toed sloth) Seldom?

MELLMPQSRKGRSKANDIHASGANLSRQYKVHTLDSWNKWELLAIIGTMIFLLYIWLYSQSFHFHVAHLY

AHFGYPSAQHIVGQRYLKGAGVVKDEEMAMHWFRRASQQNHPYASFNLAVGKMKNMTGSMEVGDVEMLLN

VAASQGIQDAQELLENVIWMKSKLLPTKRMQL

>Varanus komodoensis (Komodo dragon) Seldom?

MRQAVLRWSRRLVSKKRARVRKNMKSVMAQRKQAGRSQEGTEQVREANLSPRIKKPSPTKWSKWENSLLP

FQLLAITGSIFLLLYIMVCYENFHFHVVHMYAHLGYPNAQHIVGQRYLKGAGVEKNEEKAMQWFRQAAEK

GHPHSSFNLAVGKLKNMTAMLEEGDVEKLLNVAAGQGVQEAQNLLENIRNRHLP

>Gigantopelta aegis (deep sea snail) Seldom?

MATSGLEGASQAELRHRYQPELDPEWNPNLPYGGKVYLARKKKSDPMYVRVLEAGLIIFTFAMVYYAFFY

FDNLHFHVVHAYAHLGYAHAQHQVGQRYLHGKGVEKHDDKAMEWFRKAADQGHAHASYNLAIGHLRGMNA

GLKPGEAHTLIHHAASKGVEEADKVLNTICTQGGCD

>Ciona robusta Saxo (Cirobu.g00001698)

MKSQKWWHRPRPAGFLNFGHQAGSGTEYVDEYVRHRVPPTESFKPAEEMRKSDAKVSDETTFRLDYIPHQLSKHEPHPKEVYNPPGTPMEGVSMYRQDYPGHNTGPAQLAKRSEARSVPLVKFEAHPTYASDFKRWTIPPRVKLGPDNDYKKPVVKMENTSTFQQDYIHRFAPPRESARPPDKAFQSDVPLESHTVHRVSFIPHQIQPRLQREKEKYCAPTVPMNSETTFKQDFTGPRALPAESMRPSQAPFVSTDPLASSSEFRDSFVAWPVVRPYRKEPLKYQGPQGDMELLSTQRLDFRSLNGRPASAKRPAVRRGKVMPFEGVTNYSSDFKKWNVPRTLGKPRPEAIQQTGRFEGLSTARQHFITHSGALPARSCKPDNRAFLSDSALEDKTIYKVSYVPKSMREVERYPTPDWLKQQEDIWRKTGMMTQKTRGMTPEATAAYIRAAA

>Homo sapiens SAXO1

MKTKCICELCSCGRHHCPHLPTKIYDKTEKPCLLSEYTENYPFYHSYLPRESFKPRREYQKGPIPMEGLTTSRRDFGPHKVAPVKVHQYDQFVPSEENMDLLTTYKKDYNPYPVCRVDPIKPRDSKYPCSDKMECLPTYKADYLPWNQPRREPLRLEHKYQPASVRFDNRTTHQDDYPIKGLVKTISCKPLAMPKLCNIPLEDVTNYKMSYVAHPVEKRFVHEAEKFRPCEIPFESLTTQKQSYRGLMGEPAKSLKPLARPPGLDMPFCNTTEFRDKYQAWPMPRMFSKAPITYVPPEDRMDLLTTVQAHYTCPKGAPAQSCRPALQIKKCGRFEGSSTTKDDYKQWSSMRTEPVKPVPQLDLPTEPLDCLTTTRAHYVPHLPINTKSCKPHWSGPRGNVPVESQTTYTISFTPKEMGRCLASYPEPPGYTFEEVDALGHRIYKPVSQAGSQQSSHLSVDDSENPNQRELEVLA

**Single-chain guide RNAs (sgRNAs) used in this study**

Pax3/7.2.1 (from Kim et al. 2022)

**GTAGTGGAGATGGCAGCTCA (G+N19)**

Pax3/7.4.1 (from Kim et al. 2022)

**GGACTAATAGAACTGACCGA** **(G+N19)**

Pou4.3.21 (from Johnson et al. 2023)

**GCTGAGTGGTGGAAAGCGGG (G+N19)**

Pou4.4.106 (from Johnson et al. 2023)

**GAGGATGGAAATGATTCGGG (G+N19)**

Lhx1/5.3.21

**GTGGACGGGAAATTCTGGCA (G+N19)**

Lhx1/5.4.137

**GCAACAATAACCTTGAACCT (G+N19)**

Ddpep.1.32

**GAGAAGTACAACCAGAACAT (G+N19)**

Ddpep.1.122

**GACTGTGGAGCTAAATCGGG (G+N19)**

Saxo.2.95

**GGTTTAAACGATTCTGTAGG (G+N19)**

Saxo.3.112

**GTGTACAGACAAGACTACCC (G+N19)**

Nckap5.4.121

**GGCATTGTCAGAGAGATATG** **(G+N19)**

Nckap5.8.520

**GGAGCTCAGAAGATACAAAG (G+N19)**

Control (from Stolfi et al. 2014)

**GCTTTGCTACGATCTACATT (G+N19)**

**Primers for NGS validation of sgRNAs**

| **Gene + exon** | **Forward primer** | **Reverse primer** |
| --- | --- | --- |
| Lhx1/5 exon 3 | GTAGCAGATGTAACCACGTG | ATACGAACCGACAAAAGTCG |
| Lhx1/5 exon 4 | TTTGCAGCGACGACTTTTG | TCATGACGTACCTCCTGATG |
| Ddpep exon 1 | AAGTTATCCGGGCCTGTTTC | GGGATAGTGAGTCACTGGTAC |
| Saxo exon 2 | TCGTATTGGTTTAAATGCCGTA | GAAATTAACGAGTAAGGCAAGCA |
| Saxo exon 3 | TTTATCCTGCGTTTGACAGTG | CGACTTTCGCAATAAAGGGTA |
| Nckap5 exon 4 | CCGACATAACGTTTTAAAGGAAA | TATCATGTCTGATGCCATCAATG |
| Nckap5 exon 8 | ACAGTTTCCGATGTTAATTCGG | GTTTTGAGATGCAGGATTCAATC |

**ID numbers of gene sequences used in this study**

| **Gene name** | **KH ID** | **KY21 ID** | **ANISEED ID** |
| --- | --- | --- | --- |
| Fgf8/17/18 | C5.5 | Chr5.501 | Cirobu.g00007390 |
| Pax3/7 | C10.150 | Chr10.288 | Cirobu.g00001350 |
| Hand-related | C1.1116 | Chr1.1701 | Cirobu.g00000132 |
| MRF | C14.307 | Chr14.750 | Cirobu.g00003733 |
| Saxo | C10.475 | Chr10.763 | Cirobu.g00001698 |
| Ddpep | C12.244 | Chr12.988 | Cirobu.g00002781 |
| Defcab | C1.1218 | Chr1.2072 | Cirobu.g00000242 |
| Seldom | C4.78 | Chr4.267 | Cirobu.g00006902 |
| Dmbx | C1.1212 | Chr1.2074 | Cirobu.g00000237 |
| Pou4 | C2.42 | Chr2.456 | Cirobu.g00004616 |
| Lhx1/5 | L107.7 | Chr11.1275 | Cirobu.g00010599 |
| Engrailed | C7.431 | Chr7.541 | Cirobu.g00008277 |
| Vsx | C11.689 | Chr11.1087 | Cirobu.g00002541 |
| Ebf | L24.10 | Chr1.422 | Cirobu.g00012207 |
| VGLUT | C3.324* | Chr3.1172* | Cirobu.g00005496 |
| VGAT | C2.526 | Chr2.793 | Cirobu.g00004733 |
| Ephrin A.b | C3.202 | Chr3.875 | Cirobu.g00005364 |
| FOG | C10.574 | Chr10.450 | Cirobu.g00001805 |
| Eef1a | C14.52 | Chr14.174 | Cirobu.g00003963 |
| Foxc | L57.25 | Chr12.158 | Cirobu.g00012813 |
| Foxf | C3.170 | Chr3.989 | Cirobu.g00005328 |
| Nckap5 | C9.229 | Chr9.632 | Cirobu.g00009755 |

*Severely truncated gene models. Refer to NCBI gene model instead (NCBI Reference Sequence: NM_001128885.1)

**Electroporation mix recipes**

Fgf8>Pax3/7 for RNAseq analysis:

60 ug Fgf8/17/18 -4835/+12 “sec”>HA::Pax3/7

50 ug Fgf8/17/18 -2401/+3>TagRFP

25 ug MRF -906/-1>eGFP

25 ug Hand-related -2053/+30>eGFP

Negative control to compare to Fgf8>Pax3/7 in RNAseq analysis:

60 ug Fgf8/17/18 -4835/+12 “sec”>nls::lacZ

50 ug Fgf8/17/18 -2401/+3>TagRFP

25 ug MRF -906/-1>eGFP

25 ug Hand-related -2053/+30>eGFP

Pax3/7 CRISPR to assay Ddpep reporter expression:

70 ug Fgf8/17/18>Cas9

25 ug Fgf8/17/18>H2B::mCherry

100 ug Ddpep>Unc-76::GFP

40 ug U6>Pax3/7.2.1

40 ug U6>Pax3/7.4.1

Negative control CRISPR to compare to Pax3/7 CRISPR (Ddpep reporter):

70 ug Fgf8/17/18>Cas9

25 ug Fgf8/17/18>H2B::mCherry

100 ug Ddpep>Unc-76::GFP

80 ug U6>Control

Pax3/7 CRISPR to assay Saxo reporter expression:

70 ug Fgf8/17/18>Cas9

25 ug Fgf8/17/18>H2B::mCherry

70 ug Saxo intron 1 + bpFOG>Unc-76::GFP

40 ug U6>Pax3/7.2.1

40 ug U6>Pax3/7.4.1

Negative control CRISPR to compare to Pax3/7 CRISPR (Saxo reporter):

70 ug Fgf8/17/18>Cas9

25 ug Fgf8/17/18>H2B::mCherry

700 Saxo intron 1 + bpFOG>Unc-76::GFP

80 ug U6>Control

Pax3/7 CRISPR to assay Seldom reporter expression:

70 ug Fgf8/17/18>Cas9

25 ug Fgf8/17/18>H2B::mCherry

70 ug Seldom>Unc-76::GFP

40 ug U6>Pax3/7.2.1

40 ug U6>Pax3/7.4.1

Negative control CRISPR to compare to Pax3/7 CRISPR (Seldom reporter):

70 ug Fgf8/17/18>Cas9

25 ug Fgf8/17/18>H2B::mCherry

700 Seldom>Unc-76::GFP

80 ug U6>Control

Pax3/7 overexpression (Ebf driver) to assay Ddpep reporter:

70 ug Ebf>HA::Pax3/7

50 ug Ebf>Unc-76::mCherry

80 ug Ddpep>Unc-76::GFP

Negative control to compare to Ebf>Pax3/7 above (Ddpep reporter):

70 ug Ebf>lacZ

50 ug Ebf>Unc-76::mCherry

80 ug Ddpep>Unc-76::GFP

Pax3/7 overexpression (Ebf driver) to assay Saxo reporter:

70 ug Ebf>HA::Pax3/7

50 ug Ebf>Unc-76::mCherry

80 ug Saxo intron 1 + bpFOG>Unc-76::GFP

Negative control to compare to Ebf>Pax3/7 above (Saxo reporter):

70 ug Ebf>lacZ

50 ug Ebf>Unc-76::mCherry

80 ug Saxo intron 1 + bpFOG>Unc-76::GFP

Pax3/7 overexpression (Fgf8/17/18 driver) to assay Saxo reporter:

70 ug Fgf8/17/18>HA::Pax3/7

50 ug Fgf8/17/18>H2B::mCherry

80 ug Saxo intron 1 + bpFOG>Unc-76::GFP

Negative control to compare to Fgf8/17/18>Pax3/7 (Saxo reporter):

50 ug Fgf8/17/18>H2B::mCherry

80 ug Saxo intron 1 + bpFOG>Unc-76::GFP

Pax3/7 overexpression (Ebf driver) to assay Seldom reporter:

70 ug Ebf>HA::Pax3/7

50 ug Ebf>Unc-76::mCherry

80 ug Seldom>Unc-76::GFP

Negative control to compare to Ebf>Pax3/7 above (Seldom reporter):

70 ug Ebf>lacZ

50 ug Ebf>Unc-76::mCherry

80 ug Seldom>Unc-76::GFP

Pou4 CRISPR to assay Ddpep reporter expression:

70 ug Fgf8/17/18>Cas9

25 ug Fgf8/17/18>H2B::mCherry

100 ug Ddpep>Unc-76::GFP

40 ug U6>Pou4.3.21

40 ug U6>Pou4.4.106

Negative control CRISPR to compare to Pou4 CRISPR (Ddpep reporter):

70 ug Fgf8/17/18>Cas9

25 ug Fgf8/17/18>H2B::mCherry

100 ug Ddpep>Unc-76::GFP

80 ug U6>Control

Pou4 CRISPR to assay Seldom reporter expression:

70 ug Fgf8/17/18>Cas9

25 ug Fgf8/17/18>H2B::mCherry

70 ug Seldom>Unc-76::GFP

40 ug U6>Pou4.3.21

40 ug U6>Pou4.4.106

Negative control CRISPR to compare to Pou4 CRISPR (Seldom reporter):

70 ug Fgf8/17/18>Cas9

25 ug Fgf8/17/18>H2B::mCherry

70 ug Seldom>Unc-76::GFP

80 ug U6>Control

Pou4 CRISPR to assay Saxo reporter expression:

70 ug Fgf8/17/18>Cas9

25 ug Fgf8/17/18>H2B::mCherry

70 ug Saxo intron 1 + bpFOG>Unc-76::GFP

40 ug U6>Pou4.3.21

40 ug U6>Pou4.4.106

Negative control CRISPR to compare to Pou4 CRISPR (Saxo reporter):

70 ug Fgf8/17/18>Cas9

25 ug Fgf8/17/18>H2B::mCherry

70 ug Saxo intron 1 + bpFOG>Unc-76::GFP

80 ug U6>Control

Pou4 CRISPR to assay Dmbx/Defcab reporter expression:

70 ug Fgf8/17/18>Cas9

25 ug Fgf8/17/18>H2B::mCherry

70 ug Dmbx –3489/-2158 + bpFOG>Unc-76::YFP

40 ug U6>Pou4.3.21

40 ug U6>Pou4.4.106

Neg. ctrl. CRISPR to compare to Pou4 CRISPR (Dmbx/Defcab reporter):

70 ug Fgf8/17/18>Cas9

25 ug Fgf8/17/18>H2B::mCherry

70 ug Dmbx –3489/-2158 + bpFOG>Unc-76::YFP

80 ug U6>Control

Pou4 overexpression (Fgf8/17/18 driver) to assay Ddpep reporter:

70 ug Fgf8/17/18>Pou4

25 ug Fgf8/17/18>H2B::mCherry

100 ug Ddpep>Unc-76::GFP

Negative control to compare to Fgf8/17/18>Pou4 (Ddpep reporter):

25 ug Fgf8/17/18>H2B::mCherry

100 ug Ddpep>Unc-76::GFP

Pou4 overexpression (Fgf8/17/18 driver) to assay Seldom reporter:

70 ug Fgf8/17/18>Pou4

25 ug Fgf8/17/18>H2B::mCherry

70 ug Seldom>Unc-76::GFP

Negative control to compare to Fgf8/17/18>Pou4 (Seldom reporter):

25 ug Fgf8/17/18>H2B::mCherry

70 ug Seldom>Unc-76::GFP

Pou4 overexpression (Fgf8/17/18 driver) to assay Saxo reporter:

70 ug Fgf8/17/18>Pou4

25 ug Fgf8/17/18>H2B::mCherry

70 ug Saxo intron 1 + bpFOG>Unc-76::GFP

Negative control to compare to Fgf8/17/18>Pou4 (Saxo reporter):

25 ug Fgf8/17/18>H2B::mCherry

70 ug Saxo intron 1 + bpFOG>Unc-76::GFP

Pou4 overexpression (Fgf8/17/18 driver) to assay Dmbx/Defcab reporter:

70 ug Fgf8/17/18>Pou4

25 ug Fgf8/17/18>H2B::mCherry

70 ug Dmbx –3489/-2158 + bpFOG>Unc-76::YFP

Negative control to compare to Fgf8/17/18>Pou4 (Dmbx/Defcab reporter):

25 ug Fgf8/17/18>H2B::mCherry

70 ug Dmbx –3489/-2158 + bpFOG>Unc-76::YFP

Pou4 overexpression (Fgf8/17/18 driver) to assay Defcab-only reporter:

70 ug Fgf8/17/18>Pou4

25 ug Fgf8/17/18>H2B::mCherry

70 ug Defcab -1792/+24>Unc-76::GFP

Negative control to compare to Fgf8/17/18>Pou4 (Defcab-only reporter):

25 ug Fgf8/17/18>H2B::mCherry

70 ug Defcab -1792/+24>Unc-76::GFP

Saxo mPou4 site 1 test

70 ug Saxo intron 1 + bpFOG>Unc-76::mCherry

70 ug Saxo intron 1 mPou4 site 1 + bpFOG>Unc-76::GFP

Saxo mPou4 site 2 test

70 ug Saxo intron 1 + bpFOG>Unc-76::mCherry

70 ug Saxo intron 1 mPou4 site 2 + bpFOG>Unc-76::GFP

Saxo mPou4 site 1+2 test

70 ug Saxo intron 1 + bpFOG>Unc-76::mCherry

70 ug Saxo intron 1 mPou4 site 1+2 + bpFOG>Unc-76::GFP

Saxo wild-type (control) test

70 ug Saxo intron 1 + bpFOG>Unc-76::mCherry

70 ug Saxo intron 1 + bpFOG>Unc-76::GFP

Lhx1/5 CRISPR to assay Dmbx/Defcab reporter expression:

70 ug Fgf8/17/18>Cas9

25 ug Fgf8/17/18>H2B::mCherry

70 ug Dmbx –3489/-2158 + bpFOG>Unc-76::YFP

40 ug U6>Lhx1/5.3.21

40 ug U6>Lhx1/5.4.137

Neg. ctrl. CRISPR to compare to Lhx1/5 CRISPR (Dmbx/Defcab reporter):

70 ug Fgf8/17/18>Cas9

25 ug Fgf8/17/18>H2B::mCherry

70 ug Dmbx –3489/-2158 + bpFOG>Unc-76::YFP

80 ug U6>Control

Lhx1/5 CRISPR to assay Saxo reporter expression:

70 ug Fgf8/17/18>Cas9

25 ug Fgf8/17/18>H2B::mCherry

70 ug Saxo intron 1 + bpFOG>Unc-76::GFP

40 ug U6>Lhx1/5.3.21

40 ug U6>Lhx1/5.4.137

Negative control CRISPR to compare to Lhx1/5 CRISPR (Saxo reporter):

70 ug Fgf8/17/18>Cas9

25 ug Fgf8/17/1>H2B::mCherry

70 ug Saxo intron 1 + bpFOG>Unc-76::GFP

80 ug U6>Control

Lhx1/5 CRISPR to assay Seldom reporter expression:

70 ug Fgf8/17/18>Cas9

25 ug Fgf8/17/18>H2B::mCherry

70 ug Seldom>Unc-76::GFP

40 ug U6>Lhx1/5.3.21

40 ug U6>Lhx1/5.4.137

Negative control CRISPR to compare to Lhx1/5 CRISPR (Seldom reporter):

70 ug Fgf8/17/18>Cas9

25 ug Fgf8/17/1>H2B::mCherry

70 ug Seldom>Unc-76::GFP

80 ug U6>Control

Lhx1/5 CRISPR to assay Ddpep reporter expression:

70 ug Fgf8/17/18>Cas9

25 ug Fgf8/17/18>H2B::mCherry

100 ug Ddpep>Unc-76::GFP

40 ug U6>Lhx1/5.3.21

40 ug U6>Lhx1/5.4.137

Negative control CRISPR to compare to Lhx1/5 CRISPR (Ddpep reporter):

70 ug Fgf8/17/18>Cas9

25 ug Fgf8/17/1>H2B::mCherry

100ug Ddpep>Unc-76::GFP

80 ug U6>Control

En>Lhx1/5 looking at extended Dmbx/Defcab reporter:

80 ug En>Lhx1/5

50 ug En>Unc-76::mCherry

70 ug Dmbx –3489/-2158 + bpFOG>Unc-76::YFP

Negative control looking at extended Dmbx/Defcab reporter:

50 ug En>Unc-76::mCherry

70 ug Dmbx –3489/-2158 + bpFOG>Unc-76::YFP

En>Lhx1/5 looking at smaller Dmbx/Defcab reporter:

80 ug En>Lhx1/5

50 ug En>Unc-76::mCherry

70 ug Dmbx –2657/-2158 + bpFOG>Unc-76::GFP

Negative control looking at smaller Dmbx/Defcab reporter:

50 ug En>Unc-76::mCherry

70 ug Dmbx –2657/-2158 + bpFOG>Unc-76::GFP

En>Lhx1/5 looking at Saxo reporter:

80 ug En>Lhx1/5

25 ug En>Unc-76::mCherry

70 ug Saxo>Unc-76::GFP

Negative control looking at Saxo reporter:

25 ug En>Unc-76::mCherry

70 ug Saxo>Unc-76::GFP

En>Lhx1/5 looking at Seldom reporter:

80 ug En>Lhx1/5

25 ug En>Unc-76::mCherry

70 ug Seldom>Unc-76::GFP

Negative control looking at Seldom reporter:

25 ug En>Unc-76::mCherry

70 ug Seldom>Unc-76::GFP

En>Lhx1/5 looking at Ddpep reporter:

80 ug En>Lhx1/5

25 ug En>Unc-76::mCherry

100 ug Ddpep>Unc-76::GFP

Negative control looking at Ddpep reporter:

25 ug En>Unc-76::mCherry

100 ug Ddpep>Unc-76::GFP

Pax3/7 CRISPR in ACIN lineage (Ddpep reporter):

80 ug EprhinA.b>Cas9::Geminin-Nter

25 ug EphrinA.b>H2B::mCherry

100 ug Ddpep>Unc-76::GFP

40 ug U6>Pax3/7.2.1

40 ug U6>Pax3/7.4.1

Negative control CRISPR in ACIN lineage (Ddpep reporter):

80 ug EprhinA.b>Cas9::Geminin-Nter

25 ug EphrinA.b>H2B::mCherry

100 ug Ddpep>Unc-76::GFP

80 ug U6>Control

Overexpression of Pou4 in epidermis looking at Saxo reporter:

70 ug Foxf>Pou4

25 ug Foxf>H2B::mCherry

70 ug Saxo intron 1 + bpFOG>Unc-76::GFP

Negative control to compare to Foxf>Pou4 (Saxo reporter):

25 ug Foxf>H2B::mCherry

70 ug Saxo intron 1 + bpFOG>Unc-76::GFP

Pou4 CRISPR in epidermis to assay Saxo reporter expression:

25 ug Fog>Cas9

15 ug Fog>H2B::mCherry

70 ug Saxo intron 1 + bpFOG>Unc-76::GFP

40 ug U6>Pou4.3.21

40 ug U6>POu4.4.106

Neg. ctrl. CRISPR to compare to epidermis Pou4 CRISPR (Saxo reporter):

25 ug Fog>Cas9

15 ug Fog>H2B::mCherry

70 ug Saxo intron 1 + bpFOG>Unc-76::GFP

80 ug U6>Control

Pou4 CRISPR in epidermis to assay VGLUT reporter expression:

25 ug Fog>Cas9

15 ug Fog>H2B::mCherry

70 ug VGLUT>Unc-76::YFP

40 ug U6>Pou4.3.21

40 ug U6>POu4.4.106

Neg. ctrl. CRISPR to compare to epidermis Pou4 CRISPR (Saxo reporter):

25 ug Fog>Cas9

15 ug Fog>H2B::mCherry

70 ug VGLUT>Unc-76::YFP

80 ug U6>Control

Ddpep overexpression to look at ddN and MGIN2 axon trajectories:

80 ug Ebf>Ddpep

90 ug Vsx intron + bpFOG>Unc-76::mCherry

70 ug Dmbx –3489/-2158 + bpFOG>Unc-76::YFP

Negative control for ddN/MGIN2 axon trajectories above:

90 ug Vsx intron + bpFOG>Unc-76::mCherry

70 ug Dmbx Dmbx –3489/-2158 + bpFOG>Unc-76::YFP

Ddpep CRISPR to look at ACIN midline crossing:

80 ug EphrinA.b>Cas9::Geminin-Nter

25 ug EphrinA.b>H2B::mCherry

100 ug Ddpep>Unc-76::GFP

40 ug U6>Ddpep.1.32

40 ug U6>Ddpep.1.122

Negative control for ACIN midline crossing experiment above:

80 ug EphrinA.b>Cas9::Geminin-Nter

25 ug EphrinA.b>H2B::mCherry

100 ug Ddpep>Unc-76::GFP

80 ug U6>Control

Saxo CRISPR to look at ddN axon length:

70 ug Fgf8/17/18>Cas9

25 ug Fgf8/17/18>H2B::mCherry

70 ug Dmbx –3489/-2158 + bpFOG>Unc-76::YFP

40 ug U6>Saxo.2.95

40 ug U6>Saxo.3.112

Negative control to compare to Saxo CRISPR axon length above:

70 ug Fgf8/17/18>Cas9

25 ug Fgf8/17/18>H2B::mCherry

70 ug Dmbx –3489/-2158 + bpFOG>Unc-76::YFP

80 ug U6>Control

Nckap5 CRISPR to look at ddN axon length:

70 ug Fgf8/17/18>Cas9

25 ug Fgf8/17/18>H2B::mCherry

80 ug Dmbx –3489/-2158 + bpFOG>Unc-76::YFP

50 ug U6>Nckap5.4.121

50 ug U6>Nckap5.8.520

Negative control to compare to Nckap5 CRISPR axon length above:

70 ug Fgf8/17/18>Cas9

25 ug Fgf8/17/18>H2B::mCherry

80 ug Dmbx –3489/-2158 + bpFOG>Unc-76::YFP

100 ug U6>Control

Saxo CRISPR to look at metamorphosis:

40 ug Foxc>Cas9

10 ug Foxc>H2B::mCherry

40 ug U6>Saxo.2.95

40 ug U6>Saxo.3.112

Pou4 CRISPR to compare to Saxo CRISPR in metamorphosis:

40 ug Foxc>Cas9

10 ug Foxc>H2B::mCherry

40 ug U6>Pou4.3.21

40 ug U6>Pou4.4.106

Negative Control CRISPR to compare to Saxo CRISPR in metamorphosis:

40 ug Foxc>Cas9

10 ug Foxc>H2B::mCherry

80 ug U6>Control

Ddpep::GFP fusion localization:

70 ug Ebf>Ddpep::GFP

50 ug Ebf>Unc-76::mCherry
